# Inhibitors of membrane associated serine proteases block replication of coronavirus SARS-CoV-2 and influenza virus H1N1

**DOI:** 10.1101/2025.05.09.652710

**Authors:** Victoria Banas, Kuljeet Seehra, Matthew Mahoney, Yevhenii Kyriukha, Elizabeth Weiland, Carolina Alisio, Michelle Fath, Zhenfu Han, Traci Bricker, Michael Tartell, Sean Whelan, Adrianus C.M. Boon, James W. Janetka

**Author notes:** Corresponding author James Janetka, Washington University School of Medicine, 660 S Euclid Avenue, Campus Box 8231 St Louis MO 63110 USA. Authors contributed equally (Victoria Banas, Kuljeet Seehra).

## Abstract

TMPRSS2 is a membrane associated serine protease which is important in the viral pathogenesis of coronaviruses and influenza viruses. We developed mechanism-based covalent α-ketobenzothiazole (kbt) inhibitors using substrate specificity PS-SCL screening of TMPRSS2 as a rational guide for inhibitor design. Three distinct focused libraries of tetrapeptide kbts were synthesized and evaluated for their inhibition of TMPRSS2, matriptase and other serine proteases. We also investigated different capping groups for the previously reported tripeptide inhibitor Ac-QFR-kbt (MM3144) to increase its selectivity over the blood coagulation protease factor Xa. The most potent compounds were tested for their ability to inhibit viral replication of SARS-CoV-2 coronavirus and the H1N1 influenza virus. The most active compounds were profiled for their pharmacokinetics (PK) in mice. Several promising new compounds were identified with improved potency, selectivity, and drug-like properties including Bz-QFR-kbt (CA1043) and Cbz-QFR-kbt (ZFH9141) with an IC_50_ of 150 nM and 60 nM for H1N1, respectively.

## Introduction

TMPRSS2 is a transmembrane serine protease which is involved in the pathogenesis of cancer, in addition to coronavirus and influenza infections. The COVID-19 pandemic, caused by the SARS-CoV-2 coronavirus, has been an eye-opening experience for both the general and scientific community across the world. It has impacted all aspects of human life and the threat of future outbreaks via infections from another virus or bacteria for which currently available drugs will be ineffective, like SARS-CoV-2 at the outset. Incredibly, vaccines as well as several new drug candidates targeting this virus were developed at unprecedented speed. This was made possible by the combined efforts of scientists worldwide who elucidated details about the makeup and pathogenesis of SARS-CoV-2 infection. This work resulted in the identification of several potential therapeutic targets such as two viral proteases MPro and ClPro, as well as several host cell proteases, most importantly TMPRSS2.

The role of TMPRSS2^1^ in SARS-CoV-2^2^ infection and other viral infection from coronaviruses and influenza is related to proteolytic processing of the RNA virus’s outer glycoproteins Spike and Hemagglutinin, respectively. The Spike and HA proteins require partial proteolysis^3^ to bind to their host cell receptors, namely ACE2 in SARS-CoV-2 (**Figure 1**)^4^. TMPRSS2 is advantageously located on the surface of lung epithelial cells and is highjacked by these viruses to perform this function and thus is essential for host cell viral binding and entry in the lung. We and others have demonstrated that inhibition of TMPRSS2 with small molecule inhibitors^1, 5, 6^ also results in decreased viral replication^7^.

**Figure 1.**
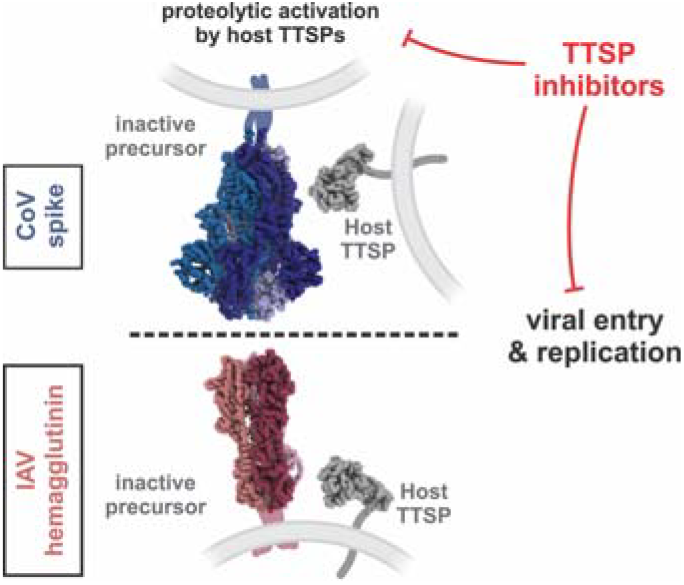
Activation of viral glycoproteins by host proteases is an essential step in the replication of multiple human viruses, including influenza and coronaviruses. Selective and potent inhibitors of these proteases would provide broad protection against current and emerging viral threats.

TMPRSS2 has a trypsin-like serine protease domain and belongs to the family of Type II Transmembrane Serine Protease (TTSP)^8^ proteolytic enzymes or membrane associated serine proteases (MASPs)^3^. In addition to SARS-CoV-2, TMPRSS2 has previously been shown to be important in other coronavirus infections^9^ caused by SARS-CoV^10, 11^ and MERS^12-14^. Furthermore, TMPRSS2^15-18^ and the other TTSPs^8, 19^, including matriptase^20, 21^ and TMPRSS11D/HAT (human airway trypsin-like protease)^16, 22^, have been demonstrated to process the Hemagglutinin (HA) protein on the surface of Influenza A viruses, allowing viral cell adhesion and entry in these infections. Furthermore, there are reports of TMPRSS4^23-25^ and TMPRSS13^26-28^ also being involved in coronavirus and influenza viral cell entry and pathogenesis. Various inhibitors of matriptase have been developed which have demonstrated activity against influenza viruses^29-31^.

We recently reported on the first peptidyl α-ketobenzothiazole (kbt) covalent reversible inhibitors of TMPRSS2^5^ and have shown they have potent antiviral effects both *in vitro* ^5^ and *in vivo*^7^ against SARS-CoV-2 infection. Here, we report on our continuing efforts to develop structure activity relationships (SAR) and optimize initial lead compounds of this class of inhibitors based on the substrate specificity of TMPRSS2 as gleaned from positional scanning of combinatorial substrate libraries (PS-SCL) screening^32^.

## Design and Synthesis

Our previously identified tetrapeptide kbt inhibitors of TMPRSS2^5^, Ac-IQFR-kbt (MM3116)-1, Ac-GQFR-kbt (MM3122)-2, and Ac-PQFR-kbt (MM3123)-3 have exquisite sub-nanomolar inhibition of TMPRSS2 (Table 5) . PS-SCL studies^32^ revealed the amino acid sidechain preferences of TMPRSS2 tetrapeptide substrates to be I in the P4, Q in the P3, and F in the P2 position of the peptide (MM3116). A P1 Arg is highly preferred like all other trypsin-like serine proteases. While this information is integral to substrate and inhibitor design, it should be noted that PS-SCL only records individual preferences for one protease subpocket (S) binding relative to the control peptide and does not account for) any cooperativity between the different protease subpockets and peptide sidechains. Thus, it is imperative to perform SAR studies evaluating several of the top amino acid sidechains and not just the best one identified from PS-SCL. For example, while MM3116 was predicted to be optimal, it turned out that replacing the P4 I with a G resulted in an almost 10-fold enhancement in potency. In the case of TMPRSS2^32^, I, G, P, M, and L are preferred in the P4 position, Q, E, M, S, and T in the P3 position and F, W, A, V, T, and S in the P2 position. Therefore, to determine the best combination of P4, P3, and P2 sidechains, we synthesized 3 focused libraries of peptide ketobenzothiazoles (kbts) keeping two of the amino acids constant based on the top preferred sidechains for those positions while changing the third position (**Table 1**). We also synthesized the physiologically relevant peptide Ac-PSKR-kbt (MM3194)-17 based on the SARS-CoV-2 Spike protein sequence which gets cleaved by TMPRSS2.

**Table 1.** SAR of P4, P3, and P2 libraries based on tetrapeptide Ac-IQFR-kbt (MM3116) from PS-SCL.

| Compound Name | Structure | TMPRSS2 IC<sub>50</sub> (nM) | Matriptase IC<sub>50</sub> (nM) | Hepsin IC<sub>50</sub> (nM) | HGFA IC<sub>50</sub> (nM) | Thrombin IC<sub>50</sub> (nM) | Factor Xa IC<sub>50</sub> (nM) |
| --- | --- | --- | --- | --- | --- | --- | --- |
| MM3116 (1)* | Ac-IQFR-kbt | 1.7 | 0.92 | 0.14 | 30 | >20,000 | 792 |
| MM3122 (2)* | Ac-GQFR-kbt | 0.03 | 0.31 | 0.19 | 33 | >20,000 | 700 |
| MM3123 (3)* | Ac-PQFR-kbt | 0.06 | 0.32 | 0.13 | 75 | >20,000 | 199 |
| MM3131 (4) | Ac-LQFR-kbt | 0.03 | 1.8 | 0.25 | 98 | >20,000 | 219 |
| MM3130 (5) | Ac-MQFR-kbt | 0.14 | 0.55 | 0.13 | 36 | >20,000 | 342 |
| ZFH7063* | Ac-FLFR-kbt | 0.13/3.0 | 7.2/414 | 0.13/2.9 | 228 | >20,000 | 26 |
| ZFH7064 (6)* | Ac-WLFR-kbt | 0.09 | 12 | 0.69 | 266 | >20,000 | 2.0 |
| MM4037 (7) | Ac-IEFR-kbt | 0.11 | 2.4 | 0.35 | 276 | >20,000 |  |
| MM4037-1 | Ac-IEFdR-kbt | 0.38 |  |  | >20,000 | >20,000 |  |
| MM4028 (8) | Ac-ITFR-kbt | 0.06 | 4.7 | 0.86 | 182 | >20,000 |  |
| MM4028-1 | Ac-ITFdR-kbt | 1.42 |  |  | 1,119 | >20,000 |  |
| MM4027 (9) | Ac-IMFR-kbt | 3.0 | 0.51 | 1.3 |  | >20,000 |  |
| MM4009 (10) | Ac-ISFR-kbt | 0.07 | 4.7 | 0.86 | 182 | >20,000 |  |
| MM4027-1 | Ac-IMFdR-kbt | 5.63 | 16.1 | 3.16 | 234 | >20,000 |  |
| MM4038 (11) | Ac-IdWFR-kbt | 0.04 | 25 | 12 | 66 | >20,000 |  |
| MM3187 (12) | Ac-IQWR-kbt | 2.2 | 1.2 | 0.32 | 1,450 | >20,000 | 129 |
| MM3186 (13) | Ac-IQTR-kbt | 0.17 | 16 | 0.98 | 18,400 | >20,000 | >20,000 |
| MM3180 (14) | Ac-IQVR-kbt | 0.03 | 0.76 | 0.11 | 799 | >20,000 |  |
| MM3180-1 | Ac-IQVdR-kbt | 2.89 | 22.9 | 1.72 | 14,600 | >20,000 | >20,000 |
| MM3178 (15) | Ac-IQAR-kbt | 0.23 | 0.10 | 1.3 | >20,000 | >20,000 | 170 |
| MM3177 (16) | Ac-IQSR-kbt | 4.5 | 1.3 | 2.4 | 19,900 | >20,000 | 1,728 |
| MM3194 (17) | Ac-PSKR-kbt | 0.29 | 0.29 | 2.4 | >20,000 | >20,000 |  |

Shown in **Table 1** are the results from screening the inhibitors for their inhibition of the six serine proteases TMPRSS2, HGFA, matriptase, hepsin, Factor Xa, and thrombin. We found most analogs displayed sub-nM activity with the most active being Ac-LQFR-kbt (4), Ac-IQVR-kbt (14), and Ac-IdWFR-kbt (11) with IC_50_s ranging from 3-4 nM while the least active were Ac-IQSR-kbt (16), Ac-IMFR-kbt (9), and Ac-IQWR-kbt (12) which however are still potent with IC_50_’s from 2.2 to 4.5 nM…Ac-WLFR-kbt (ZFH7064-6) ^32^

We first tested the most active analogs for their ability to block viral entry (**Table 2**) of the chimeric VSV SARS-CoV-2 virus (ref) into Calu-3 human lung epithelial cells. We found previous inhibitors MM3122 and MM3123 still performed the best compared to any new compounds with EC_50_’s below nM. Next, we tested the best analogs for their effect on inhibiting replication of SARS-CoV-2 and Influenza strain H1N1 viruses. MM3122 still performed the best against both SARS-CoV-2 and H1N1 in blocking replication in Calu-3 cells (4.1 log and 4.3 log fold reduction for CoV-2 (@0.1 µM) and H1N1 (@10 µM) respectively). Compounds performing almost as well relative to MM3122 against H1N1 were MM4009, MM4038, with MM3123, MM3130, MM3180, and MM3178 showing a little less effect. MM4038 (Ac-IdWFR-kbt) however performed to a much lesser extent for SARS-CoV-2 only reducing the virus by 0.43 log fold. This is consistent with the viral entry data indicating this compound is much less effective with an EC_50_ of 226 nM. Interestingly MM3194, which is based on the actual S2 site cleavage of the Spike protein, only showed a 0.6 log fold reduction of the H1N1 virus at 10 µM but it was not tested for inhibition of SARS-CoV-2 since it did not effectively inhibit viral entry (EC_50_ = 613 nM).

**Table 2.** Antiviral cell entry and replication inhibition data.

| Compound Name | Structure | VSV SARS-CoV-2 Chimera EC<sub>50</sub> | SARS-CoV-2 titer Log fold reduction | H1N1 titer Log fold reduction | H1N1 titer Log fold reduction |
| --- | --- | --- | --- | --- | --- |

| | | (nM) | (0.1 μM) | (10 μM) | (1 μM) |
| --- | --- | --- | --- | --- | --- |
| MM3116 (1)* | Ac-IQFR-kbt | 3.6 | 1.3 |  |  |
| MM3122 (2)* | Ac-GQFR-kbt | 0.43 | 4.1 | 4.3 | 3.4 |
| MM3123 (3)* | Ac-PQFR-kbt | 0.86 | 1.3 | 2.6 | 1.3 |
| MM3131 (4) | Ac-LQFR-kbt | 11 |  |  |  |
| MM3130 (5) | Ac-MQFR-kbt | 48 | 1.1 | 2.7 | 1.4 |
| ZFH7064 (6)* | Ac-WLFR-kbt |  | 1.7 |  |  |
| MM4037 (7) | Ac-IEFR-kbt | 24 | 1.5 |  |  |
| MM4028 (8) | Ac-ITFR-kbt | 103 |  |  |  |
| MM4027 (9) | Ac-IMFR-kbt | 92 |  |  |  |
| MM4009 (10) | Ac-ISFR-kbt | 40 |  | 3.4 | 0.62 |
| MM4038 (11) | Ac-IdWFR-kbt | 226 | 0.48 | 3.0 | 2.1 |
| MM3187 (12) | Ac-IQWR-kbt | 22 | 2.0 |  |  |
| MM3186 (13) | Ac-IQTR-kbt | 19 | 0.53 |  |  |
| MM3180 (14) | Ac-IQVR-kbt | 12 | 2.4 | 2.6 | 2.1 |
| MM3178 (15) | Ac-IQAR-kbt | 11 | 1.2 | 2.4 | 2.0 |
| MM3177 (16) | Ac-IQSR-kbt | 36 | 0.42 |  |  |
| MM3194 (17) | Ac-PSKR-kbt | 613 |  | 0.60 | 0 |

In previous studies^5^, we identified a tripeptide, Ac-QFR-kbt (MM3144)-**18** and another group reported on MeSO_2_ -QFR-kbt (N-0385)-**26**^6^ which both had potency (IC_50_ 0.3 nM) equivalent to MM3122-**2** (Ac-GQFR-kbt, IC_50_, but both are not selective over Factor Xa, the coagulation protease which is essential to get selectivity over as well as thrombin. One reason for the required selectivity is to prevent any blood thinning in infected patients in which the marketed Factor Xa inhibitor apixaban is used for. The second reason is that Factor Xa has been shown to proteolytically cleave the Spike protein of SARS-CoV-2 into and form that is incapable of binding the ACE2 receptor^33^. Here, we explored different N-terminal capping groups of MM3144-17 in place of the acetyl to find improved inhibitors with better potency and selectivity over Factor Xa. These included substitutions on the amine with alkyl, acyl, and sulfonamide groups.

Shown in **Table 3** is the biological activity (IC_50_s) of these inhibitors against TMPRSS2 and the other 5 serine proteases. We found the benzoyl group in **23** and the benzyl sulfone in **28** were the most potent against TMPRSS2 and improved over the previously reported methyl sulfone N-0385 (**26**)^6, **29**^. When the amine is alkylated as in **19** (dimethyl) and **24** (piperidine), and **21** (benzyl) a significant loss in potency was observed. Interestingly, this decreased inhibition was also found with the phenyl sulfone derivative **27**, suggesting the additional methylene spacer of the benzyl is necessary for optimal interaction with TMPRSS2. All but 2 of these 12 capped tripeptides were tested for their antiviral activity against SARS-CoV-2 by measuring the fold-reduction in viral load (replication) at a 0.1 µM concentration of inhibitor. The best 2 compounds were MM3144 (**18**) and ZFH9141 (**25**) with a 3.9- and 3.7-fold reduction in virus, respectively. When tested against influenza H1N1 at a concentration of 10 and 1 µM, these showed a 2.5- and 2.8-fold reduction. Better compounds against H1N1 included CA1043 (**23**), MF1101 (**27**), and MF1104 (**29**) with a 3.3-fold reduction being the best displayed by **27**.

**Table 3.**
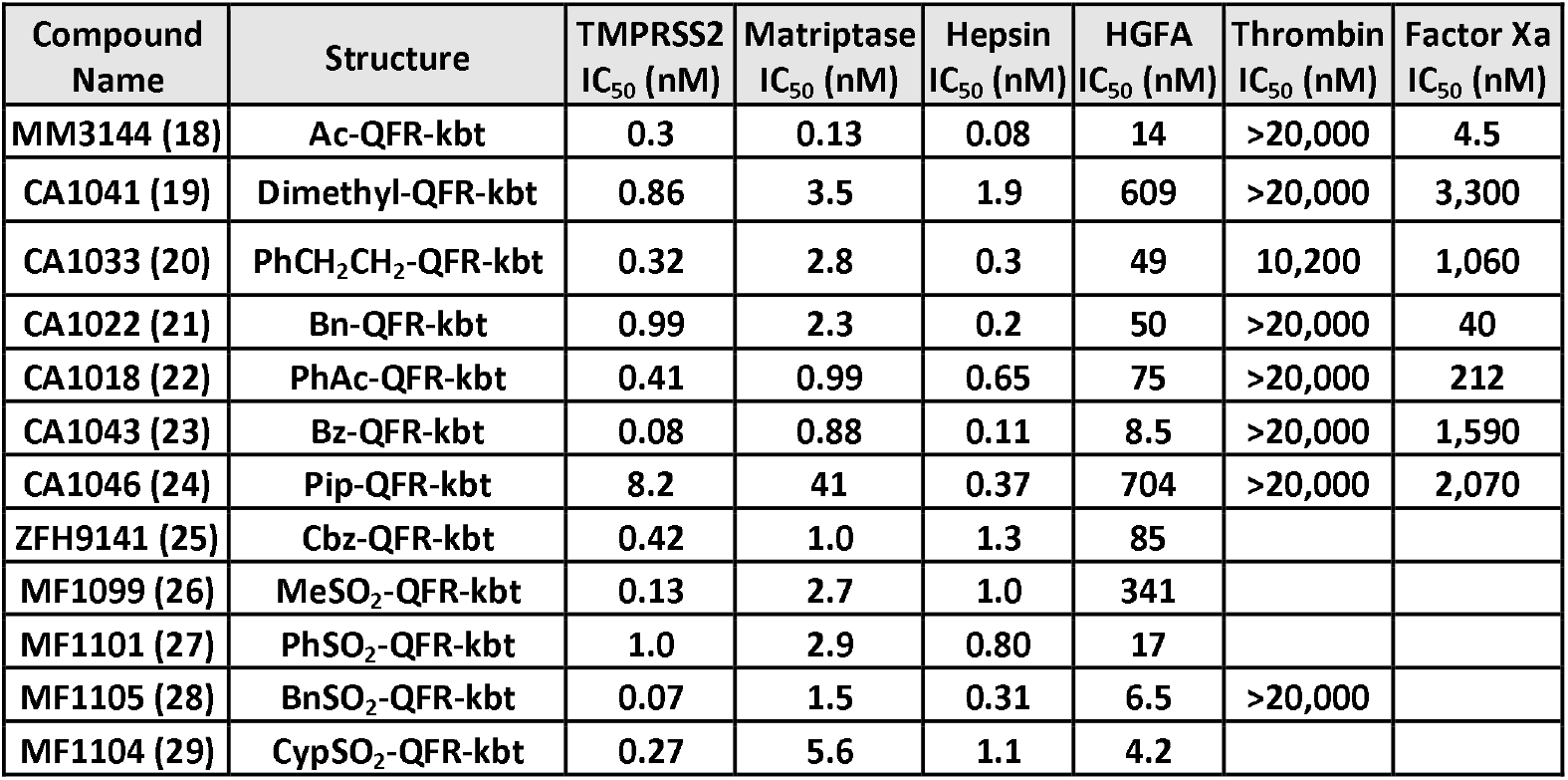
Enzyme SAR of N-terminal capped QFR-kbt.

| Compound Name | Structure | TMPRSS2<br>IC<sub>50</sub> (nM) | Matriptase<br>IC<sub>50</sub> (nM) | Hepsin<br>IC<sub>50</sub> (nM) | HGFA<br>IC<sub>50</sub> (nM) | Thrombin<br>IC<sub>50</sub> (nM) | Factor Xa<br>IC<sub>50</sub> (nM) |
| --- | --- | --- | --- | --- | --- | --- | --- |
| MM3144 (**18**) | Ac-QFR-kbt | 0.3 | 0.13 | 0.08 | 14 | >20,000 | 4.5 |
| CA1041 (**19**) | Dimethyl-QFR-kbt | 0.86 | 3.5 | 1.9 | 609 | >20,000 | 3,300 |
| CA1033 (**20**) | PhCH<sub>2</sub>CH<sub>2</sub>-QFR-kbt | 0.32 | 2.8 | 0.3 | 49 | 10,200 | 1,060 |
| CA1022 (**21**) | Bn-QFR-kbt | 0.99 | 2.3 | 0.2 | 50 | >20,000 | 40 |
| CA1018 (**22**) | PhAc-QFR-kbt | 0.41 | 0.99 | 0.65 | 75 | >20,000 | 212 |
| CA1043 (**23**) | Bz-QFR-kbt | 0.08 | 0.88 | 0.11 | 8.5 | >20,000 | 1,590 |
| CA1046 (**24**) | Pip-QFR-kbt | 8.2 | 41 | 0.37 | 704 | >20,000 | 2,070 |
| ZFH9141 (**25**) | Cbz-QFR-kbt | 0.42 | 1.0 | 1.3 | 85 |  |  |
| MF1099 (**26**) | MeSO<sub>2</sub>-QFR-kbt | 0.13 | 2.7 | 1.0 | 341 |  |  |
| MF1101 (**27**) | PhSO<sub>2</sub>-QFR-kbt | 1.0 | 2.9 | 0.80 | 17 |  |  |
| MF1105 (**28**) | BnSO<sub>2</sub>-QFR-kbt | 0.07 | 1.5 | 0.31 | 6.5 | >20,000 |  |
| MF1104 (**29**) | CypSO<sub>2</sub>-QFR-kbt | 0.27 | 5.6 | 1.1 | 4.2 |  |  |

**Table 4.** Antiviral replication data.

| Compound Name | Structure | SARS-CoV-2 titer<br>Log fold reduction<br>(0.1 μM) | H1N1 titer Log<br>fold reduction<br>(10 μM) | H1N1 titer Log<br>fold reduction<br>(1 μM) |
| --- | --- | --- | --- | --- |
| MM3144 (18)<br>(N-0386)* | Ac-QFR-kbt | 3.9 | 2.5 | 2.4 |
| CA1033 (20) | PhCH<sub>2</sub>CH<sub>2</sub>-QFR-kbt | 0.81 |  |  |
| CA1022 (21) | Bn-QFR-kbt | 0.95 |  |  |
| CA1018 (22) | PhAc-QFR-kbt | 2.6 |  |  |
| CA1043 (23) | Bz-QFR-kbt | 2.6 | 3.0 | 3.1 |
| ZFH9141 (25) | Cbz-QFR-kbt | 3.7 | 2.8 | 2.8 |
| MF1099 (26)<br>(N-0385)* | MeSO<sub>2</sub>-QFR-kbt | 2.0 |  |  |
| MF1101 (27) | PhSO<sub>2</sub>-QFR-kbt | 2.3 | 3.3 | 2.9 |
| MF1105 (28) | BnSO<sub>2</sub>-QFR-kbt | 2.5 |  |  |
| MF1104 (29) | CypSO<sub>2</sub>-QFR-kbt | 1.9 | 3.0 | 2.9 |

For the most active compounds we determine the IC_50_ values against H1N1 as shown in Table 5 and found the most potent compound to be ZFH9141 (**25**) with an IC_50_ of 0.06 µM followed closely by CA1043 (**23**) having an IC_50_ of 0.15 µM which are significantly improved over previously reported inhibitors MM3122 (**2**) and MM3144 (**18**). Interestingly a report of a compound similar to ZFH9141 (**25**) but with a ketothiazole warhead (N-0920) was just published^34^. The sulfonamide derivatives MF1101 (**27**) and MF1104 (**29**) also showed excellent potencies with IC_50_s of 0.67 and 0.87 µM, respectively. Interestingly, the compound based on the actual SARS-CoV-2 peptide cleavage sequence, MM3194 (**17**), was essentially inactive with an IC_50_ >10 µM.

**Table 5.** Concentration response of inhibitors and corresponding IC50 values.

| Compound Name | H1N1 Fold inhibition @ indicated μM concentration |  |  |  |  | No Drug control | IC<sub>50</sub> (μM) |
| --- | --- | --- | --- | --- | --- | --- | --- |
|  | 10 | 1 | 0.1 | 0.01 | 0.001 |  |  |
| ZFH9141 (25) | 578 | 578 | 483 | 6 | 2 | 1.1 | 0.06 |
| CA1043 (23) | 774 | 968 | 5 | 1 | 1 | 1.4 | 0.15 |
| MM3144 (18) | 298 | 239 | 34 |  |  | 0.8 | 0.4 |
| MM3122 (2) | 2952 | 2216 | 27.5 | 4.2 | 2.1 | 1 | 0.64 |
| MF1101 (27) | 1321 | 1014 | 4 | 1 | 1 | 1 | 0.67 |
| MF1104 (29) | 916 | 617 | 1 | 2 | 1 | 1.3 | 0.87 |
| MM3178 (15) | 238 | 91 | 2 |  |  | 1.1 | 1.3 |
| MM3180 (14) | 359 | 135 | 2 |  |  | 0.8 | 1.3 |
| MM3130 (5) | 510 | 27 | 1 |  |  | 1 | 2.8 |
| MM4009 (10) | 743 | 3 | 2 |  |  | 1.1 | 3 |
| MM3194 (17) | 4 | 1 | 1 |  |  | 0.8 | >10 |

We tested CA1043 (**23**) for its pharmacokinetics (PK) in mice (**Fig. 2**) and found it had significantly improved compound exposure (AUC) when dose IP (20 mg/kg) compared to MM3122 which had been previously tested^7^. Increased antiviral replication of CA1043 in H1N1 coupled with its improved PK suggests it will show increased efficacy relative to MM3122 when tested in animals^7^. We are currently testing ZFH9141 (**25**) for its PK in mice.

**Figure 2.**
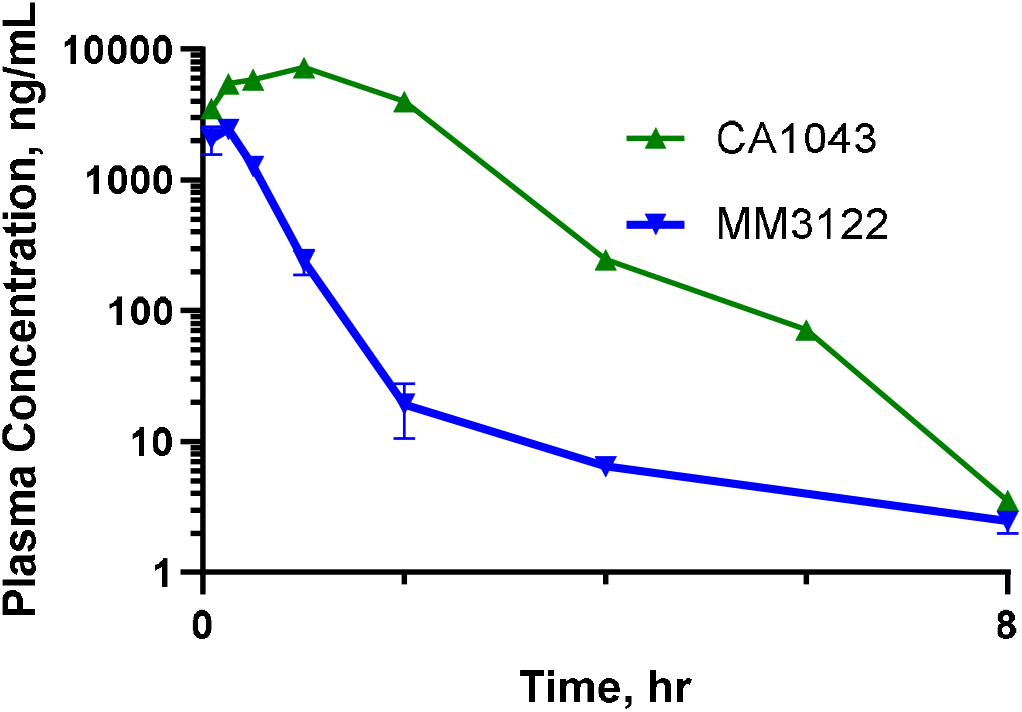
Pharmacokinetics of MM3122 (2) and CA1043 (23) in mice after IP dosing.

## Conclusions

We have significantly improved upon earlier reported inhibitors of TMPRSS2 and matriptase which potently block SARS CoV-2 coronavirus and H1N1 influenza virus replication. This demonstrates the ability to produce broad-spectrum antiviral drugs from one compound which targets TMPRSS2 and matriptase, the major proteolytic activators of viral glycoproteins, Spike and HA, necessary for host cell entry and replication in all coronaviruses and influenza A viruses. We identified ZFH9141 and CA1043 as the 2 most promising inhibitors which have significantly improved pharmacokinetics relative to previous lead MM3122. We are currently testing these lead compounds for their efficacy against SARS-CoV-2 and H1N1 in animal models. We are also engaged in further optimization of ZFH9141 and will report on our progress in a future communication.

### Experimental and Methods

#### Matriptase Protease Domain Protein Expression and Purification

The catalytic domain of Human Matriptase was purified as published elsewhere^35, 36^ with minor modifications. Briefly, the DNA constructs encoding for the sequence ranging from residue 596 to 855 was purchased as a g-block from IDT (Coralville, Iowa), amplified by PCR and cloned in pET28a vector at NdeI/XhoI restriction sites to place an N-terminal His6-tag. The DNA sequence was confirmed by sanger sequencing performed by Genewiz (South Plainfield, NJ), then the plasmid transformed in E. Coli Rosetta2(DE3)pLysS cells (EMD Chemicals, Novagen, Gibbstown, NJ) for protein over-expression. Transformed cells were grown in LB media at 37°C until OD600 reached 0.6-0.8, then chilled at 4°C for 30 mins, and subsequentially induced with 0.7mM IPTG and incubated at 16°C overnight. Finally, cells were harvested and responded in lysis buffer (20 mM sodium phosphate (pH 7.3), 400 mM NaCl, 10% (v/v) glycerol, 5 mM BME). Cells were lysed by sonication, and the inclusion bodies spun down and washed three times with lysis buffer, then dissolved overnight in lysis buffer supplemented with 8M urea and 5 mM imidazole. The resulting mixture was spun to remove cells debris and the supernatant loaded on HisPur Ni-NTA column (Thermo Scientific, Waltham, MA), followed by 3 volume wash with lysis buffer containing 8M urea and 5 mM imidazole, 3 volume wash with lysis buffer with 8M urea and 20 mM imidazole, and finally the protein eluted with lysis buffer containing 8M urea and 250 mM imidazole. Matriptase was refolded by removing the urea sequentially through the following dialysis steps: 1) lysis buffer containing 2M urea and 0.5 M arginine; 2) lysis buffer containing 1M urea and 0.5 M arginine; 3) lysis buffer containing 0.5 M arginine and; 4) twice against lysis buffer. During refolding, Matriptase self-activated and cleaved-off the pro-peptide (residues 596-614) together with the His6-tag. Finally, protein aggregates were removed by centrifugation, the protein concentrate to about 10 mg/ml by ultracentrifugation using vivaspin turbo 15 devices (Sartorius, Göttingen, Germany) and injected on HiLoad 16/600 superdex 75 pg (GE Healthcare, Salt Lake City, UT) size exclusion column using 20 mM Hepes (pH 7.0), 150 mM NaCl and 5% (v/v) glycerol as running buffer. Purified fractions were combined and concentrated to the desired working concentration for crystallographic studies or dialyzed against running buffer having 40% (v/v) glycerol for storage. Enzymatic activity was confirmed by fluorometric cleavage assay as reported below. The matriptase construct carrying the C731S^36^ mutation was generated by site-specific mutagenesis using the wild-type gene cloned in pET28a as a template and purified as reported above. Sequences of the g-block DNA and primers are reported in the supplementary material.

#### Full-length human TMPRSS2 expression and purification

TMPRSS2 was expressed and purified as described previously with some modifications^59^. Briefly, plasmid containing the TMPRSS2 ectodomain 109-492 plus EFVEHHHHHHHH (C-terminus) and honey bee mellitin tag (N terminus) was purchased from Addgene (176412). Plasmid was transformed into Escherichia coli D10Bac cells to generate the baculovirus bacmid following the protocol in the Bac-to-Bac ® Baculovirus Expression System (Gibco). The resultant bacmid was transfected into Gibco™ Sf9 cells in Sf-900™ II SFM (Cat.# 11496015) following the jetOPTIMUS protocol for “Reverse Transfection” (Polyplus Ref. # 101000051). The generated p0 was then used to make a p1 then a p2. Four liters of sf9 insect cells grown in Gibco™ Sf-900™ II SFM (Cat.# 10902104) at a density of between 2.5-4×10^6^ cells/mL. were infected with 8mL per 500mL of the p2 virus. Two and 3 days after infection, cells were harvested. The cell debris was spun down, 10x PBS pH 7.4 added to the supernatant to create 1X PBS, and the pH brought to 7.4 with NaOH. After spinning down the resulting precipitate, 60ml Ni-NTA XPure Agarose Resin (UBPBio) was added to the supernatant, and batch bound for 2 hours at 16°C with shaking at 120 rpm.After binding, the resin was then added to a gravity flow column at 4°C and washed with multiple column volumes of 1X PBS pH 7.4. After washing, the protein was eluted with 1X PBS pH 7.4 with 500mM Imidazole. Fractions containing TMPRSS2 were concentrated in Amicon® Ultra Centrifugal Filters, 30 kDa MWCO (Cat.# UFC9030) and buffer exchanged to less than 5mM Imidazole with SEC buffer (50 mM Tris pH 7.5, 250 mM NaCl) to allow for TMPRSS2 activation. Activation was achieved by leaving the protein on ice overnight. After activation, the protein was further purified via a HiLoad 16/600 Superdex 75 pg (Cytiva) size exclusion column at 4°C using SEC buffer. Fractions containing the TMPRSS2 were concentrated in Amicon® Ultra Centrifugal Filters, 30 kDa MWCO (Millipore) to 10mg/ml and stored in SEC buffer with 25% Glycerol at -80°C.

#### Fluorogenic Kinetic Enzyme Inhibitor Assays of HGFA, matriptase, TMPRSS2, and hepsin

Inhibitors (11-pt serial dilutions, concentrations ranged from 2 µM to 0.15 pM final concentration in reaction) were serially diluted in DMSO (2% DMSO final concentration) and then mixed with recombinant HGFA (1514-SE-010, R&D Systems), matriptase, TMPRSS2 or hepsin* (4776-SE-010, R&D Systems) in black 384 well plates (Corning # 3575). The final assay concentration for HGFA, matriptase, TMPRSS2 and hepsin 7.5 nM, 0.2 nM, 3 nM, and 0.3 nM, respectively in TNC buffer (25 mM Tris, 150 mM NaCl, 5 mM CaCl_2_, 0.01% Triton X-100, pH 8). After thirty minutes incubation at room temperature, Boc-QLR-AMC substrate (K_m_ = 37 µM) was added to the HGFA assays and Boc-QAR-AMC substrate was added to the matriptase (K_m_ = 93 µM), TMPRSS2 (K_m_ = 16 µM), and hepsin (K_m_ = 156 µM) assays. The final substrate concentrations for all assays were at the K_m_ for the respective enzymes. Changes in fluorescence (excitation at 380 nm and emission at 460 nm) were measured at room temperature over time in a Biotek Synergy 2 plate reader (Winnoski, VT). Using GraphPad Prism version 6.04 software program, (GraphPad Software, San Diego, CA, www.graphpad.com), a four-parameter curve fit was used to determine the inhibitor IC_50_s from a plot of the mean reaction velocity versus the inhibitor concentration. The IC_50_ values represent the average of three separate experimental determinations.

##### *Hepsin Activation

Recombinant Hepsin (10 μg, 0.44mg/mL) was diluted to 2.4 μM in TNC buffer (25 mM Tris, 150 mM NaCl, 5 mM CaCl_2_, 0.01% Triton X-100, pH 8) and incubated at 37°C. After twenty-four hours, the hepsin was diluted in glycerol to 50%. This stock hepsin (1.2 μM) was stored in a -20°C freezer and diluted in TNC buffer for use in assays.

#### Chromogenic Kinetic Enzyme Assay of Thrombin and Factor Xa

Inhibitors (11-pt serial dilutions, 0-20 µM final concentration) were serially diluted in DMSO (2% DMSO final concentration) and then mixed with recombinant thrombin (0.15 nM final concentration) or Factor Xa (0.35 nM final concentration) in TNC buffer (25 mM Tris, 150 mM NaCl, 5 mM CaCl_2_, 0.01% Triton X-100, pH 8) using clear 384 well plates. After incubating for 30 minutes at 25* C, the chromogenic substrate (S2238; D-Phe-Pip-Arg-pNA) for thrombin (K_m_ = 14.5 µM) (#1473-SE-010, R&D Systems) or (S2222; Bz-Ile-Glu-Gly-Arg-pNA) for Factor Xa (K_m_ = 200 µM) was added to a final concentration of K_m_ (4 x K_m_ (50 µM) for thrombin) in a final reaction volume of 40 microliters. Changes in absorbance at 405 nm were measured over time in a Biotek Synergy 2 plate (Winnoski, VT). Using GraphPad Prism version 6.04 software program, (GraphPad Software, San Diego, CA, www.graphpad.com), a four-parameter curve fit was used to determine the inhibitor IC_50_s from a plot of the mean reaction velocity versus the inhibitor concentration.

#### Cells and Viruses

Vero cells expressing human angiotensin converting enzyme 2 (ACE2) and transmembrane protease serine 2 (TMPRSS2) (Vero-hACE2-hTMPRSS2^25, 37^, gift from Adrian Creanga and Barney Graham, NIH) were cultured at 37°C in Dulbecco’s Modified Eagle medium (DMEM) supplemented with 10% fetal bovine serum (FBS), 10⍰mM HEPES (pH 7.3), 100⍰U/mL of Penicillin, 100 µg/mL of Streptomycin, and 10 µg/mL of puromycin. Vero cells expressing TMPRSS2 (Vero-hTMPRSS2)^25^ were cultured at 37°C in DMEM supplemented with 10% fetal bovine serum (FBS), 10⍰mM HEPES (pH 7.3), 100⍰U/mL of Penicillin, 100µg/mL of Streptomycin, and 5 µg/mL of blasticidin. Calu-3 cells were cultured in DMEM media supplemented with 1.0 mM sodium pyruvate, non-essential amino-acids (NEAA), 100 U/mL of penicillin, 100 µg/mL streptomycin, 2.0 mM L-glutamine, 10 mM HEPES, and 10% Fetal Bovine Serum (FBS). SARS-CoV-2 (strain WA1/2020) was propagated on Vero-hTMPRSS2 cells. The virus stocks were subjected to next-generation sequencing, and the S protein sequences were identical to the original isolates. The infectious virus titer was determined by plaque and focus--forming assay on Vero-hACE2-hTMPRSS2 or Vero-hTMPRSS2 cells. The influenza A virus strain A/California/4/2009 (pdmH1N1) was propagated in 10-day old embryonated chicken eggs, aliquoted, and stored at -80°C. The infectious virus titer was determined by focus-forming assay (FFA) on Madin-Darby canine kidney (MDCK) cells.

#### In vitro TMPRSS2 inhibition assays on SARS-CoV-2

Calu-3 cells (5 x 10^5^ cells/well) were seeded in 24-well culture plates in infection medium (DMEM + 1.0 mM Sodium pyruvate, NEAA, 100 U/mL of penicillin, 100 µg/mL streptomycin, 2.0 mM L-glutamine, 10 mM HEPES, and 2% FBS) and incubated overnight at 37°C and 5% CO_2_. After 24 h, media was removed and fresh 250 µL media was added to each well containing the TMPRSS2 inhibitors or starting at 20 µM concentration and diluted 10-fold to 2, 0.2, 0.02 and 0.002 µM. Media alone, and DMSO were included as negative controls. Next, the cells were transferred to the BSL3 laboratory and 250 µL of media containing 4,000 PFU of SARS-CoV-2 was added for 1 h at 37°C and 5% CO_2_. Note, that the final concentration of MM3122 and Remdesivir is 10, 1, 0.1, 0.01, and 0.001 µM. After 1 h, the virus inoculum was removed, the cells were washed twice with infection media and fresh infection media containing the inhibitor or controls were added to each well. At 48 h post--infection, culture supernatant is collected and used to quantify virus titers by RT-qPCR.

#### SARS-CoV-2 RT-qPCR assay

To quantify the amount of virus in the supernatant of SARS-CoV-2 infected and treated Calu-3 cells, RNA was extracted from 100 µL supernatant using the MagMax Viral Pathogen Kit (Thermo Fisher Scientific) on the KingFisher Flex Purification System following the manufacturer’s protocol and eluted with 50 µL of water. Four microliters RNA was used for real-time RT-qPCR to detect and quantify N gene of SARS-CoV-2 using TaqMan™ RNA--to-CT 1-Step Kit (Thermo Fisher Scientific) as described^38^ using the following primers and probes: Forward: GACCCCAAAATCAGCGAAAT; Reverse: TCTGGTTACTGCCAGTTGAATCTG; Probe: ACCCCGCATTACGTTTGGTGGACC; 5’Dye/3’Quencher: 6-FAM/ZEN/IBFQ. Viral RNA was expressed as N gene copy numbers per mL based on a standard included in the assay, which was created via *in vitro* transcription of a synthetic DNA molecule containing the target region of the N gene. These values were then plotted in GraphPad Prism 10.4 and used to calculate the fold inhibition compared to the control treated cells.

#### In vitro TMPRSS2 inhibition assays on Influenza A virus

Calu-3 cells (2.5 x 10^5^ cells/well) were seeded in 24-well culture plates in infection medium (DMEM + 1.0 mM Sodium pyruvate, NEAA, 100 U/mL of penicillin, 100 µg/mL streptomycin, 2.0 mM L-glutamine, 10 mM HEPES, and 2% FBS) and incubated overnight at 37°C and 5% CO_2_. After 24 h, media was removed and 250 µL of infection media (MEM + 0.1% bovine serum albumin, 100⍰U/mL of Penicillin, and 100 µg/mL of Streptomycin) containing 25 focus forming units (FFU) of pdmH1N1 (A/California/4/2009) was added for 1 h at 37°C and 5% CO_2_. After 1 h, the virus inoculum was removed, the cells were washed twice with PBS and fresh infection media containing 10, 1, 0.1, 0.01, or 0.001 µM of TMPRSS2 inhibitors or controls were added to each well. At 72 h post-infection, culture supernatant is collected and used to quantify virus titers by focus forming assay as described below.

#### Influenza virus titration assays

FFA’s were performed on MDCK cells in 96-well plates. The virus stock of tissue culture supernatant from treated or control Calu-3 cells were diluted serially by 10-fold, starting at 1:10, in infection medium. One hundred microliters of the diluted virus were added to a two wells per dilution per sample. After 1 h at 37°C, the inoculum was aspirated, the cells were washed with PBS, and a 1% methylcellulose overlay in MEM plus 100⍰U/mL of Penicillin, 100 µg/mL of Streptomycin, 25mM HEPES (Corning), 0.75% NaHCO_3_ (S8761, Sigma Aldrich), 0.1% BSA, 1x Vitamins (Corning), and 1.0 µg/mL of TPCK Trypsin was added. Twenty-four hours after virus inoculation, the cells were fixed with 4% formalin for 30 minutes. To visualize infected cells, the monolayer was incubated with 100 µL/well of permeabilization buffer (PBS with 0.5% Saponin and 2% FBS) for 15 minutes at room temperature. Following two washes with PBS, 50 µl of permeabilization buffer containing an anti-influenza virus monoclonal antibody (CR9114) was added for 1 hour at 20°C. After two washes with PBS + 0.05% Tween-20 (PBST), a secondary HRP conjugated anti-human IgG (1:1000 in permeabilization buffer, A6029) was added to each well for 1 h at 20°C. Next, the cells were washed two times with PBST, and the assay was developed with TMD substrate kit (SK-4400, Vector Laboratories) according to manufacturer’s instructions. The plates were analyzed on a BioSpot reader (Cellular Technology Limited ImmunoSpot) and used to determine the focus forming units per mL (FFU/mL) in each sample. These values were then plotted in GraphPad Prism 10.4 and used to calculate the fold inhibition compared to the control treated condition and the 50% inhibitory concentration using non-linear regression analysis.

#### Mouse pharmacokinetic (PK) studies

Compounds CA1043 (**23**) and ZFH9141 (**25**) were tested for their PK in mice (single IP dosing, 20 mg/kg). Three animals (male, CD-1) were dosed and then blood was drawn at eight different sampling time points (0.08, 0.25, 0.5, 1, 2, 4, 8, 24 h) post dosing. Bioanalysis of plasma samples was performed using standard LC/MS/MS techniques.

## Supplementary Material

Details of chemical synthesis of compounds and spectral data (NMR, HPLC, MS) are provided. The procedure used for the K_M_ determination of TMPRSS2 is also included.

