## Supplementary Material for "Inhibitors of membrane associated serine proteases block replication of coronavirus SARS-CoV-2 and influenza virus H1N1"

### General synthesis, purification, and analytical chemistry procedures.

Starting materials, reagents, and solvents were purchased from commercial vendors unless otherwise noted. 1H NMR spectra were measured on a Varian 400 MHz or Bruker Avance III 600 MHz NMR instrument. The chemical shifts were reported as δ ppm relative to TMS using residual solvent peak as the reference unless otherwise noted. The following abbreviations were used to express the multiplicities: s = singlet; d = doublet; t = triplet; q = quartet; m = multiplet. High-performed liquid chromatography (HPLC) was carried out on GILSON GX-281 using Waters C18 5 μM, 4.6×50 mm and Waters Prep C18 5 μM, 19×150 mm reverse phase columns, eluted with a gradient system of 5:95 to 95:5 acetonitrile:water with a buffer consisting of 0.05% TFA. Mass spectra (MS) were performed on HPLC/MSD using electrospray ionization (ESI) for detection. All reactions were monitored by thin layer chromatography (TLC) carried out on Merck silica gel plates (0.25 mm thick, 60F254), visualized by using UV (254 nm) or dyes such as $KMnO_4$, *p*-Anisaldehyde, and CAMA (Cerium Ammonium Molybdate or Hanessian's Stain). Silica gel chromatography was carried out on a Teledyne ISCO CombiFlash purification system using pre-packed silica gel columns (12 g to 330 g sizes). All compounds used for biological assays are greater than 95% purity based on NMR and HPLC by absorbance at 220 nm and 254 nm wavelengths.

Compounds MM3116 (1)[1], MM3122 (2)[1], MM3123 (3)[1], ZFH7064 (6)[1], MM3144 (18)[1] and MF1099 (26)[2] were prepared as described earlier.

#### General procedure for synthesis of peptides[3]

**Solid phase peptide coupling and deprotection:** Into a reaction vial (with a fritted glass filter) under nitrogen containing 2-chlorotrityl resin (0.5 mmol) modified with corresponding amino acid (Phe, Trp, Thr, Val, Ala, Ser or Lys) was added DMF/$CH_2Cl_2$ (15/15 ml). The mixture was shaken at RT for 30 min and then filtered. The resin was washed with DMF (10 ml) 2 times. A mixture of Fmoc-AA-OH (2.5 mmol) in DMF (20 ml), HBTU (0.853 g, 2.25 mmol), and $iPr_2NEt$ (0.87 ml, 5 mmol) was stirred at RT for 10 min and then added to the resin. The resultant heterogeneous mixture was shaken at RT overnight and then filtered. The resin was washed with DMF (4×20 ml), dried, and then piperidine/DMF (20% v/v, 30 ml) was added. The mixture was shaken for 1-4 h at RT, then filtered and washed with DMF (4×10 ml). Following the Fmoc deprotection, the peptide is carried on to the next step, or coupling of another Fmoc-AA-OH was performed identically as described above, and then subsequently a final Fmoc deprotection to the tripeptide.

**Acetyl capping and cleavage from resin**: The peptide-containing resin was suspended in 30 ml of 0.5M $Ac_2O$ and 1M $iPr_2NEt$ in DMF and shaken at RT for 1 h. The reaction was filtered, and the resin was washed with DMF (4×10 ml) followed by $CH_2Cl_2$ (4×10 ml). The resin was then suspended in 30 ml of 25% v/v HFIP/$CH_2Cl_2$ and shaken for 1 h. The reaction was filtered, and the filtrate was concentrated and dried in a vacuum.

**Coupling of Arg-(Pbf)-kbt:HCl and final deprotection**: To crude peptide acid (400 mg, 1.0 mmol) dissolved in dry DMF (10 ml) under a nitrogen atmosphere at 0 °C was added HATU (456 mg, 1.20 mmol) followed by stirring for 15 min. Next, Arg-(Pbf)-kbt:HCl (638 mg,

1.1 mmol) and $iPr_2NEt$ (0.87 ml, 5.0 mmol) were added to the reaction at 0 °C. The reaction was allowed to reach room temperature and then stirred for an additional 2-3 h. DMF was removed under vacuum and water (250 ml) was added to the residue. The precipitate was filtered and washed with water (2×50 ml) and then dried under vacuum. The precipitate was suspended in 10 ml of TFA/thioanisole/water (92/4/4 v/v/v) and stirred for 2 h at RT. The solvent was removed, and cold diethyl ether (100 ml) was added. The resulting precipitate was collected by centrifugation and the crude product was purified by HPLC (C18 5 μM, 19×150 mm column; eluent: acetonitrile/water (0.05% TFA)) to give the final compound.

**General procedure for the synthesis of $H_2N$-Q(Trt)-F-COOMe dipeptide:** Into a round bottom flask containing Boc-Gln(Trt)-OH (5 g, 10 mmol) and HATU (4.57 g, 12 mmol) in DMF (20 ml) was added Phe-OMe×HCl (2.37 g, 11 mmol) and $iPr_2NEt$ (9 ml, 50 mmol) at 0°C and stirred overnight under argon atmosphere. The reaction mixture was poured into 300 ml of water and the precipitated peptide was collected by filtration and dried in a vacuum. The Boc protecting group was removed by adding 10 ml of 4M HCl solution in dioxane and 5 ml of dioxane to peptide stirring for 2 h under argon. The reaction mixture was concentrated and dried in a vacuum to get crude amine $H_2N$-Q(Trt)-F-COOMe.

**General procedure for the synthesis of $RSO_2$-$H_2N$-Q-R-kbt tripeptide:** Into a round bottom flask containing Boc-Gln(Trt)-OH (5 g, 10 mmol) and HATU (4.57 g, 12 mmol) in DMF (20 ml) was added Phe-OMe×HCl (2.37 g, 11 mmol) and $iPr_2NEt$ (9 ml, 50 mmol) at 0°C and stirred overnight under argon atmosphere. The reaction mixture was poured into 300 ml of water and the precipitated peptide was collected by filtration and dried in a vacuum. To the residue (5.3 g, 8.2 mmol) 50 ml of MeOH was added, followed by 50 ml of water, and 20 ml of THF. LiOH (364 mg, 15.1 mmol) was added to the reaction mixture and stirred for 36 h. Solvents were removed by evaporation and 40 ml of water was added to the residue. Then, 1M HCl solution was added dropwise to pH 3 at 0°C. The reaction mixture was incubated for 30 min at 0°C, the formed precipitate was collected by filtration, washed with water, and dried under a vacuum. The crude peptide acid residue (581 mg, 0.9 mmol) was dissolved in DMF (5 ml). To the solution under argon atmosphere at 0°C was added HATU (410 mg, 1.08 mmol) and stirred for 5 min. Next, Arg(Pbf)-kbt×HCl (530 mg, 0.94 mmol) and $iPr_2NEt$ (323 μl, 1.8 mmol) were added to the reaction and stirred overnight. The reaction mixture was poured into 100 ml of ice-cold water. The precipitate was collected by filtration, washed with water, and dried in a vacuum. The crude peptide was purified on a Silica column using Ethyl acetate/hexane gradient as an eluent. To the tripeptide (600 mg, 0.52 mmol) 10 ml of 4M HCl solution in dioxane was added and stirred for 2h under Argon. Solvents were removed under reduced pressure. Free amine (100 mg, 0.092 mmol) was dissolved in DCM at 0°C. Then $Et_3N$ (52 μl, 0.36 mmol) was added to the reaction mixture, followed by corresponding sulfochloride (0.18 mmol, 2 eq). The reaction mixture was stirred 1h at rt under Argon. Then, the reaction mixture was diluted with DCM, the organic layer was washed with $NaHCO_3$ sat., dried over sodium sulfate and the solvent was removed under vacuum. To the residue, 10 ml of TFA/thioanisole/water mixture (92/4/4 v/v/v) was added and stirred for 30 min. The solvent was evaporated, and the residue was dried in a vacuum. The crude product was purified by HPLC to give the final compound.

### Synthesis of tetrapeptidyl ketobenzothiazoles, 1-18

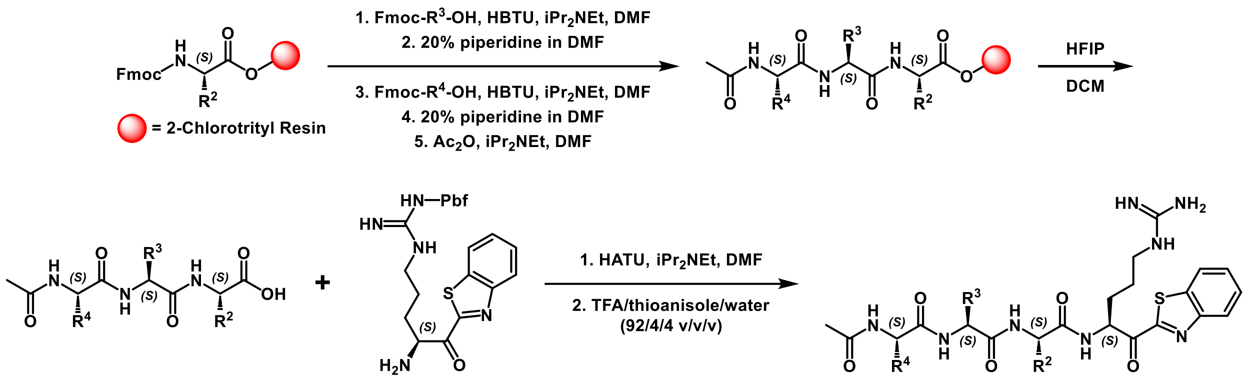


*Scheme S1. Synthesis of tetrapeptide ketobenzothiazoles (kbts) 1-18.*

We have previously published the methodology for the synthesis of the kbt peptide inhibitors[3]. As shown in Scheme S1. Synthesis of tetrapeptide ketobenzothiazoles (kbts) 1-18.Scheme S1, we first construct the tripeptide on the 2-chlorotrityl resin using standard Fmoc-solid phase peptide synthesis protocols including HBTU for amide bond coupling and 20% piperidine for Fmoc deprotection steps. The peptide was then capped with an acetyl group using acetic anhydride. Cleavage of the 2-chlorotrityl resin without removal of the protecting groups is accomplished using HFIP, then H-Arg(Pbf)-kbt is installed with HATU or EDC/HOBt in DMF. Final deprotection of the amino acid sidechains using TFA:water:thioanisole (95:2.5:2.5 %v/v) generates the target compounds which are purified by reverse-phase prep HPLC.

**MM3116 (1) or Ac-IQFR-kbt**

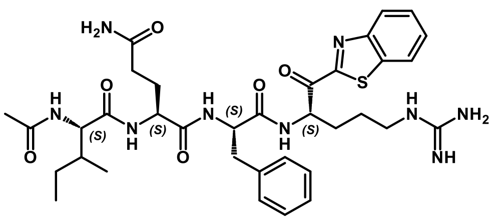

**(2S)-2-((2S)-2-acetamido-3-methylpentanamido)-N1-((S)-1-(((S)-1-(benzo[d]thiazol-2-yl)-5-guanidino-1-oxopentan-2-yl)amino)-1-oxo-3-phenylpropan-2-yl)pentanediamide**

Yield: 20 mg (25%). ESI-MS $[M+H]^+$ calcd. $C_{35}H_{48}N_9O_6S$ 722.89, found 722.6.

$^1$H NMR (400 MHz, dmso) δ 8.56 (d, *J* = 6.8 Hz, 1H), 8.32 – 8.22 (m, 2H), 8.15 (d, *J* = 7.5 Hz, 1H), 7.95 (t, *J* = 9.4 Hz, 2H), 7.68 (dd, *J* = 6.6, 3.6 Hz, 2H), 7.52 (d, *J* = 5.9 Hz, 1H), 7.26 (s, 1H), 7.24 – 7.10 (m, 5H), 6.81 (s, 1H), 5.55 – 5.46 (m, 1H), 4.65 – 4.51 (m, 1H), 4.21 – 4.12 (m, 1H), 4.08 (t, *J* = 7.6 Hz, 1H), 3.22 – 3.10 (m, 2H), 3.09 – 2.96 (m, 1H), 2.78 (dd, *J* = 14.0, 9.7 Hz, 1H), 2.13 – 1.90 (m, 3H), 1.87 (s, 3H), 1.85 – 1.72 (m, 2H), 1.72 – 1.56 (m, 4H), 1.52 – 1.30 (m, 1H), 1.14 – 1.02 (m, 1H), 0.88 – 0.70 (m, 6H).

**MM3122 (2) or Ac-GQFR-kbt**

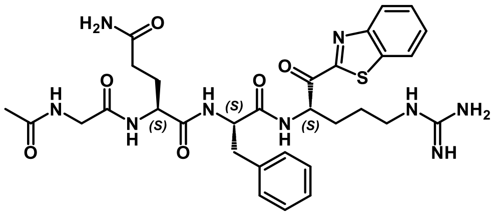

**(S)-2-(2-acetamidoacetamido)-N1-((S)-1-(((S)-1-(benzo[d]thiazol-2-yl)-5-guanidino-1-oxopentan-2-yl)amino)-1-oxo-3-phenylpropan-2-yl)pentanediamide**

Yield: 35 mg (35%). ESI-MS $[M+H]^+$ calcd. $C_{31}H_{40}N_9O_6S$ 666.78, found 666.5.

$^1$H NMR (400 MHz, dmso) δ 8.52 (d, *J* = 6.6 Hz, 1H), 8.27 (t, *J* = 8.5 Hz, 2H), 8.21 – 8.09 (m, 2H), 8.02 (d, *J* = 8.2 Hz, 1H), 7.78 – 7.59 (m, 2H), 7.60 – 7.45 (m, 1H), 7.29 – 7.11 (m, 6H), 6.80 (s, 1H), 5.56 – 5.43 (m, 1H), 4.62 – 4.47 (m, 1H), 4.27 – 4.08 (m, 1H), 3.68 (d, *J* = 5.6 Hz, 2H), 3.22 – 3.09 (m, 2H), 3.03 (d, *J* = 13.8 Hz, 1H), 2.86 – 2.73 (m, 1H), 2.10 – 1.92 (m, 3H), 1.87 (s, 3H), 1.84 – 1.71 (m, 2H), 1.68 – 1.54 (m, 3H).

**MM3123 (3) or Ac-PQFR-kbt**

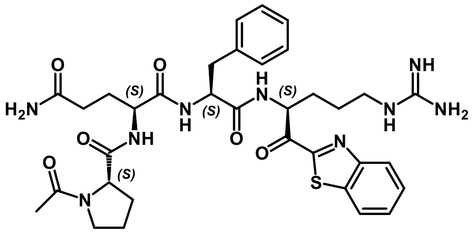

**(S)-2-((S)-1-acetylpyrrolidine-2-carboxamido)-N1-((S)-1-(((S)-1-(benzo[d]thiazol-2-yl)-5-guanidino-1-oxopentan-2-yl)amino)-1-oxo-3-phenylpropan-2-yl)pentanediamide**

Yield: 39 mg (35%). ESI-MS $[M+H]^+$ calcd. $C_{34}H_{44}N_9O_6S$ 706.84, found 706.6.

$^1$H NMR (400 MHz, $CD_3OD$) δ 9.25 (d, *J* = 6.3 Hz, 1H), 8.29 – 8.19 (m, 2H), 8.14 (d, *J* = 7.8 Hz, 1H), 8.04 (d, *J* = 8.2 Hz, 1H), 7.64 (p, *J* = 7.3 Hz, 2H), 7.32 – 7.10 (m, 5H), 5.77 – 5.66 (m, 1H), 4.74 – 4.62 (m, 1H), 4.36 – 4.29 (m, 1H), 4.24 (d, *J* = 5.8 Hz, 1H), 3.89 – 3.78 (m, 1H), 3.69 – 3.56 (m, 1H), 3.40 – 3.19 (m, 7H), 2.97 (dd, *J* = 14.0, 10.2 Hz, 1H), 2.35 – 1.68 (m, 16H).

**MM3131 (4) or Ac-LQFR-kbt**

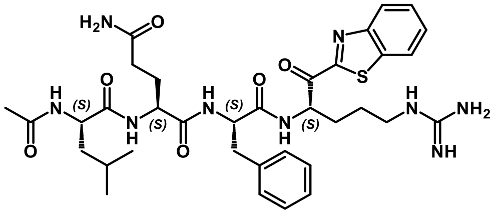

**(S)-2-((S)-2-acetamido-4-methylpentanamido)-N1-((S)-1-(((S)-1-(benzo[d]thiazol-2-yl)-5-guanidino-1-oxopentan-2-yl)amino)-1-oxo-3-phenylpropan-2-yl)pentanediamide**

Yield: 43 mg (31%). ESI-MS $[M+H]^+$ calcd. $C_{35}H_{48}N_9O_6S$ 722.89, found 722.4.

$^1$H NMR (400 MHz, MeOD) δ 8.75 (d, *J* = 6.7 Hz, 1H), 8.37 (d, *J* = 7.8 Hz, 1H), 8.32 (d, *J* = 6.1 Hz, 1H), 8.23 (d, *J* = 7.9 Hz, 1H), 8.17 – 8.08 (m, 2H), 7.72 – 7.58 (m, 2H), 7.25 – 7.07 (m, 5H), 5.76 – 5.65 (m, 1H), 4.65 (q, *J* = 7.6 Hz, 1H), 4.33 – 4.18 (m, 2H), 3.41 – 3.12 (m, 2H),

2.97 (dd, $J$ = 14.0, 9.5 Hz, 1H), 2.32 – 1.47 (m, 15H), 0.97 (dd, $J$ = 6.6, 2.0 Hz, 3H), 0.93 (dd, $J$ = 6.6, 2.0 Hz, 3H).

**MM3130 (5) or Ac-MQFR-kbt**

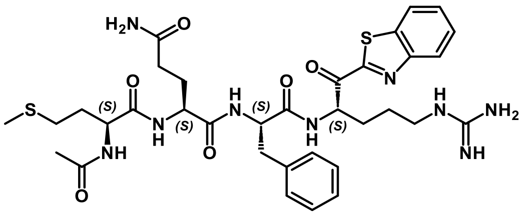

**(S)-2-((S)-2-acetamido-4-(methylthio)butanamido)-N1-((S)-1-(((S)-1-(benzo[d]thiazol-2-yl)-5-guanidino-1-oxopentan-2-yl)amino)-1-oxo-3-phenylpropan-2-yl)pentanediamide**

Yield: 35 mg (25%). ESI-MS $[M+H]^+$ calcd. $C_{34}H_{46}N_9O_6S_2$ 740.92, found 740.3.

$^1H$ NMR (400 MHz, dmso) δ 8.58 (d, $J$ = 7.0 Hz, 1H), 8.28 (t, $J$ = 8.4 Hz, 2H), 8.15 (dd, $J$ = 14.5, 7.4 Hz, 2H), 7.96 (d, $J$ = 8.2 Hz, 1H), 7.74 – 7.63 (m, 2H), 7.53 (d, $J$ = 5.5 Hz, 1H), 7.34 – 7.09 (m, 6H), 6.82 (s, 1H), 5.51 (q, $J$ = 6.0 Hz, 1H), 4.62 – 4.52 (m, 1H), 4.25 (q, $J$ = 6.8 Hz, 1H), 4.15 (q, $J$ = 7.2 Hz, 1H), 3.16 (q, $J$ = 6.7 Hz, 2H), 3.07 – 2.98 (m, 1H), 2.77 (dd, $J$ = 14.3, 10.0 Hz, 1H), 2.46 – 2.38 (m, 2H), 2.09 – 1.57 (m, 16H).

**MM4037 (7) or Ac-IEFR-kbt**

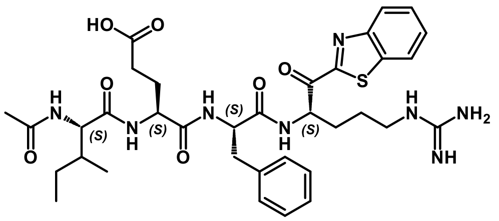

**(4S)-4-((2S)-2-acetamido-3-methylpentanamido)-5-(((S)-1-(((S)-1-(benzo[d]thiazol-2-yl)-5-guanidino-1-oxopentan-2-yl)amino)-1-oxo-3-phenylpropan-2-yl)amino)-5-oxopentanoic acid**

Yield: 24 mg (29 %). ESI-MS $[M+H]^+$ calcd. $C_{35}H_{47}N_8O_7S$ 723.87, found 723.6.

$^1H$ NMR (399 MHz, dmso) δ 12.09 (s, 1H), 8.62 (d, $J$ = 6.6 Hz, 1H), 8.33 – 8.23 (m, 2H), 7.96 (dd, $J$ = 12.7, 8.0 Hz, 2H), 7.89 (d, $J$ = 7.8 Hz, 1H), 7.68 (dt, $J$ = 6.6, 2.7 Hz, 2H), 7.48 (t, $J$ = 5.6 Hz, 1H), 7.23 – 7.11 (m, 5H), 5.49 (q, $J$ = 6.9 Hz, 1H), 4.58 (td, $J$ = 9.0, 4.5 Hz, 1H), 4.18 (q, $J$ = 8.4 Hz, 1H), 4.08 (t, $J$ = 7.8 Hz, 1H), 3.14 (q, $J$ = 7.2 Hz, 2H), 3.00 (dd, $J$ = 14.2, 4.5 Hz, 1H), 2.76 (dd, $J$ = 14.2, 9.5 Hz, 1H), 2.21 – 2.09 (m, 2H), 1.97 (d, $J$ = 6.6 Hz, 1H), 1.85 (s, 3H), 1.78 (s, 2H), 1.71 – 1.53 (m, 4H), 1.46 – 1.31 (m, 1H), 1.14 – 0.98 (m, 1H), 0.81 – 0.72 (m, 6H).

**MM4028 (8) or Ac-ITFR-kbt**

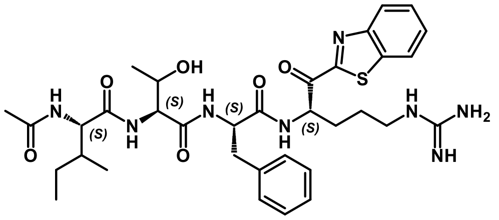

**(2S)-2-acetamido-N-((2S)-1-(((S)-1-(((S)-1-(benzo[d]thiazol-2-yl)-5-guanidino-1-oxopentan-2-yl)amino)-1-oxo-3-phenylpropan-2-yl)amino)-3-hydroxy-1-oxobutan-2-yl)-3-methylpentanamide**

Yield: 29 mg (23 %). ESI-MS $[M+H]^+$ calcd. $C_{34}H_{47}N_8O_6S$ 695.86, found 695.6.

$^1$H NMR (399 MHz, dmso) δ 8.62 (d, *J* = 6.6 Hz, 1H), 8.28 (ddd, *J* = 12.5, 6.4, 3.1 Hz, 2H), 7.97 (d, *J* = 8.6 Hz, 1H), 7.85 (d, *J* = 8.2 Hz, 1H), 7.74 – 7.62 (m, 3H), 7.51 – 7.43 (m, 1H), 7.23 – 7.08 (m, 5H), 5.48 (q, *J* = 5.6 Hz, 1H), 4.89 (d, *J* = 5.4 Hz, 1H), 4.61 (h, *J* = 4.7 Hz, 1H), 4.21 – 4.13 (m, 2H), 3.97 – 3.84 (m, 1H), 3.14 (q, *J* = 6.6 Hz, 2H), 3.02 (dd, *J* = 14.4, 4.7 Hz, 1H), 2.81 (dd, *J* = 14.2, 9.1 Hz, 1H), 1.95 (s, 1H), 1.85 (s, 3H), 1.82 – 1.64 (m, 2H), 1.58 (d, *J* = 16.3 Hz, 2H), 1.39 (s, 1H), 1.07 (dt, *J* = 16.0, 7.6 Hz, 1H), 0.95 (d, *J* = 6.2 Hz, 3H), 0.83 – 0.71 (m, 6H).

**MM4027 (9) or Ac-IMFR-kbt**

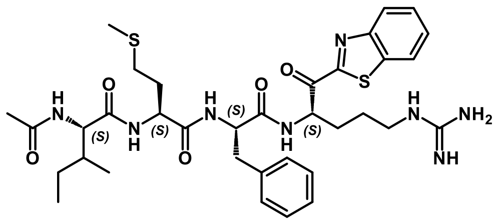

**(2S)-2-acetamido-N-((6S,9S,12S)-1-amino-6-(benzo[d]thiazole-2-carbonyl)-9-benzyl-1-imino-8,11-dioxo-15-thia-2,7,10-triazahexadecan-12-yl)-3-methylpentanamide**

Yield: 17 mg (13 %). ESI-MS $[M+H]^+$ calcd. $C_{35}H_{49}N_8O_5S_2$ 725.95, found 725.6.

$^1$H NMR (399 MHz, dmso) δ 8.61 (d, *J* = 7.0 Hz, 1H), 8.32 – 8.23 (m, 2H), 8.01 (d, *J* = 7.8 Hz, 1H), 7.94 (d, *J* = 8.2 Hz, 1H), 7.88 (d, *J* = 8.6 Hz, 1H), 7.73 – 7.62 (m, 2H), 7.48 (t, *J* = 5.8 Hz, 1H), 7.29 – 7.14 (m, 6H), 5.48 – 5.39 (m, 1H), 4.66 (q, *J* = 8.4 Hz, 1H), 4.28 (td, *J* = 8.6, 5.1 Hz, 1H), 4.08 (t, *J* = 7.8 Hz, 1H), 3.07 (q, *J* = 6.6 Hz, 2H), 2.94 (dd, *J* = 13.8, 6.0 Hz, 1H), 2.81 (dd, *J* = 13.6, 8.6 Hz, 1H), 2.32 (hept, *J* = 7.0 Hz, 2H), 2.03 – 1.56 (m, 12H), 1.54 – 1.31 (m, 3H), 1.04 (dt, *J* = 16.0, 8.0 Hz, 1H), 0.74 (t, *J* = 7.2 Hz, 6H).

**MM4009 (10) or Ac-ISFR-kbt**

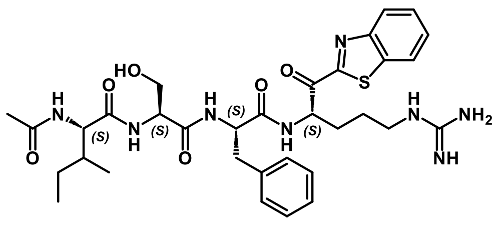

**(2S)-2-acetamido-N-((S)-1-(((S)-1-(((S)-1-(benzo[d]thiazol-2-yl)-5-guanidino-1-oxopentan-2-yl)amino)-1-oxo-3-phenylpropan-2-yl)amino)-3-hydroxy-1-oxopropan-2-yl)-3-methylpentanamide**

Yield: 24 mg ( 36 %). ESI-MS $[M+H]^+$ calcd. $C_{33}H_{45}N_8O_6S$ 681.83, found 681.5.

$^1$H NMR (399 MHz, dmso) δ 8.55 (d, *J* = 7.0 Hz, 1H), 8.33 – 8.23 (m, 2H), 7.98 – 7.88 (m, 3H), 7.69 (td, *J* = 7.2, 3.9 Hz, 2H), 7.48 (t, *J* = 5.8 Hz, 1H), 7.21 – 7.10 (m, 5H), 5.54 – 5.44 (m, 1H), 4.95 (t, *J* = 5.6 Hz, 1H), 4.57 (dq, *J* = 9.0, 4.3 Hz, 1H), 4.26 (q, *J* = 6.0 Hz, 1H), 4.15 (t, *J* = 7.8 Hz, 1H), 3.50 (q, *J* = 5.1 Hz, 2H), 3.14 (q, *J* = 7.2 Hz, 2H), 3.03 (dd, *J* = 14.0, 4.7 Hz, 1H), 2.79 (dd,

$J$ = 14.4, 9.3 Hz, 1H), 1.96 (d, $J$ = 6.2 Hz, 1H), 1.85 (s, 3H), 1.82 – 1.51 (m, 4H), 1.38 (ddd, $J$ = 10.9, 7.2, 3.7 Hz, 1H), 1.13 – 0.98 (m, 1H), 0.78 (t, $J$ = 7.4 Hz, 6H).

**MM4038 (11) or Ac-IdWFR-kbt**

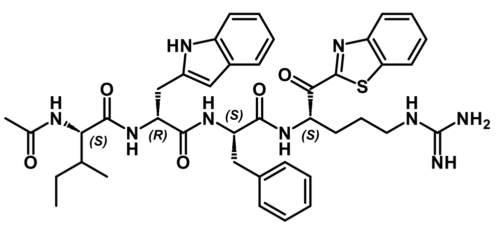

**(2S)-2-acetamido-N-((R)-1-(((S)-1-(((S)-1-(benzo[d]thiazol-2-yl)-5-guanidino-1-oxopentan-2-yl)amino)-1-oxo-3-phenylpropan-2-yl)amino)-3-(1H-indol-2-yl)-1-oxopropan-2-yl)-3-methylpentanamide**

Yield: 27 mg (32 %). ESI-MS $[M+H]^+$ calcd. $C_{41}H_{50}N_9O_5S$ 780.97, found 780.6.

$^1$H NMR (399 MHz, dmso) δ 10.72 (s, 1H), 8.70 (dd, $J$ = 18.5, 6.8 Hz, 1H), 8.42 (t, $J$ = 9.5 Hz, 1H), 8.33 – 8.24 (m, 2H), 8.12 – 8.04 (m, 1H), 7.78 (d, $J$ = 8.6 Hz, 1H), 7.74 – 7.63 (m, 2H), 7.59 (d, $J$ = 8.2 Hz, 1H), 7.53 – 7.43 (m, 1H), 7.31 – 7.09 (m, 7H), 7.06 – 6.98 (m, 2H), 6.95 (t, $J$ = 7.0 Hz, 1H), 5.54 (t, $J$ = 10.3 Hz, 1H), 4.77 – 4.64 (m, 1H), 4.52 – 4.41 (m, 1H), 4.10 (t, $J$ = 8.0 Hz, 1H), 3.20 – 3.14 (m, 1H), 3.14 – 3.07 (m, 1H), 3.07 – 2.95 (m, 1H), 2.82 – 2.68 (m, 2H), 2.65 – 2.53 (m, 1H), 2.08 – 1.81 (m, 2H), 1.78 (d, $J$ = 7.0 Hz, 3H), 1.70 – 1.62 (m, 1H), 1.58 – 1.45 (m, 1H), 1.46 – 1.35 (m, 1H), 1.15 – 1.00 (m, 1H), 0.82 – 0.75 (m, 1H), 0.62 – 0.54 (m, 3H), 0.40 (dd, $J$ = 7.0, 3.5 Hz, 3H).

**MM3187 (12) or Ac-IQWR-kbt**

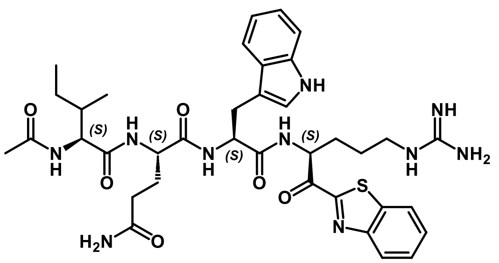

**(2S)-2-((2S)-2-acetamido-3-methylpentanamido)-N1-((S)-1-(((S)-1-(benzo[d]thiazol-2-yl)-5-guanidino-1-oxopentan-2-yl)amino)-3-(1H-indol-3-yl)-1-oxopropan-2-yl)pentanediamide**

Yield: 18 mg (26 %). ESI-MS $[M+H]^+$ calcd. $C_{37}H_{49}N_{10}O_6S$ 761.92, found 761.6.

$^1$H NMR (399 MHz, dmso) δ 10.77 (s, 1H), 8.59 – 8.47 (m, 1H), 8.32 – 8.23 (m, 2H), 7.99 – 7.80 (m, 2H), 7.73 – 7.62 (m, 2H), 7.55 – 7.39 (m, 2H), 7.35 – 7.18 (m, 2H), 7.11 (dd, $J$ = 10.5, 2.3 Hz, 1H), 7.08 – 6.86 (m, 2H), 6.84 – 6.71 (m, 1H), 5.56 – 5.37 (m, 1H), 4.59 (p, $J$ = 7.8 Hz, 1H), 4.24 – 4.13 (m, 1H), 4.13 – 4.04 (m, 1H), 3.18 – 3.02 (m, 3H), 2.94 (td, $J$ = 15.2, 8.0 Hz, 1H), 1.14 – 1.00 (m, 1H), 0.82 – 0.71 (m, 6H), 8.23 – 8.08 (m, 1H), 2.14 – 2.01 (m, 2H), 1.95 (s, 1H), 1.86 (s, 3H), 1.86 – 1.53 (m, 6H), 1.52 – 1.32 (m, 2H).

**MM3186 (13) or Ac-IQTR-kbt**

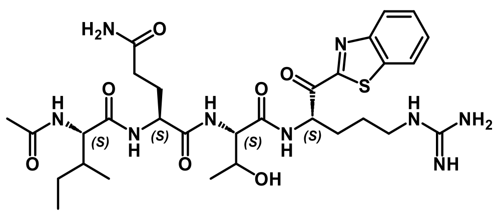


**(2S)-2-((2S)-2-acetamido-3-methylpentanamido)-N1-((2S)-1-(((S)-1-(benzo[d]thiazol-2-yl)-5-guanidino-1-oxopentan-2-yl)amino)-3-hydroxy-1-oxobutan-2-yl)pentanediamide**

Yield: 23 mg (22 %). ESI-MS $[M+H]^+$ calcd. $C_{30}H_{46}N_9O_7S$ 676.81, found 676.6.

$^1$H NMR (399 MHz, dmso) δ 8.43 – 8.33 (m, 1H), 8.32 – 8.23 (m, 2H), 8.18 (t, *J* = 8.8 Hz, 1H), 7.98 – 7.88 (m, 1H), 7.72 – 7.56 (m, 3H), 7.47 – 7.37 (m, 1H), 7.30 – 7.19 (m, 1H), 6.86 – 6.73 (m, 1H), 5.61 – 5.46 (m, 1H), 4.93 – 4.77 (m, 1H), 4.37 – 4.19 (m, 2H), 4.13 (t, *J* = 7.8 Hz, 1H), 4.02 – 3.87 (m, 1H), 3.17 – 3.11 (m, 2H), 2.16 – 2.07 (m, 2H), 2.04 – 1.88 (m, 2H), 1.86 (s, 3H), 1.78 – 1.64 (m, 3H), 1.64 – 1.52 (m, 2H), 1.51 – 1.34 (m, 1H), 1.16 – 0.95 (m, 4H), 0.85 – 0.73 (m, 6H).

**MM3180 (14) or Ac-IQVR-kbt**

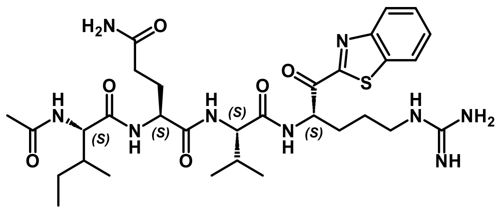

**(2S)-2-((2S)-2-acetamido-3-methylpentanamido)-N1-((S)-1-(((S)-1-(benzo[d]thiazol-2-yl)-5-guanidino-1-oxopentan-2-yl)amino)-3-methyl-1-oxobutan-2-yl)pentanediamide**

Yield: 16 mg (14 %). ESI-MS $[M+H]^+$ calcd. $C_{31}H_{48}N_9O_6S$ 674.84, found 674.6.

$^1$H NMR (399 MHz, dmso) δ 8.61 (d, *J* = 6.2 Hz, 1H), 8.26 (ddd, *J* = 12.5, 6.6, 3.1 Hz, 2H), 8.11 (d, *J* = 7.8 Hz, 1H), 7.90 (d, *J* = 8.6 Hz, 1H), 7.74 – 7.60 (m, 3H), 7.48 (t, *J* = 5.8 Hz, 1H), 7.26 (s, 1H), 6.79 (s, 1H), 5.52 – 5.43 (m, 1H), 4.29 – 4.19 (m, 2H), 4.14 (t, *J* = 8.0 Hz, 1H), 3.14 (q, *J* = 6.6 Hz, 2H), 2.18 – 2.03 (m, 2H), 2.00 – 1.89 (m, 2H), 1.85 (s, 3H), 1.83 – 1.53 (m, 6H), 1.46 – 1.35 (m, 1H), 1.13 – 1.01 (m, 1H), 0.90 – 0.74 (m, 13H).

**MM3178 (15) or Ac-IQAR-kbt**

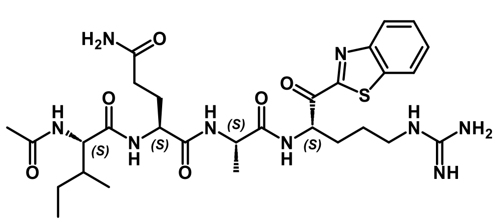

**(2S)-2-((2S)-2-acetamido-3-methylpentanamido)-N1-((S)-1-(((S)-1-(benzo[d]thiazol-2-yl)-5-guanidino-1-oxopentan-2-yl)amino)-1-oxopropan-2-yl)pentanediamide**

Yield: 40 mg (52 %). ESI-MS $[M+H]^+$ calcd. $C_{29}H_{44}N_9O_6S$ 646.79, found 646.5.

$^1$H NMR (399 MHz, dmso) δ 8.55 – 8.46 (m, 1H), 8.32 – 8.22 (m, 2H), 8.16 – 8.04 (m, 1H), 7.98 – 7.77 (m, 2H), 7.73 – 7.62 (m, 2H), 7.49 (q, *J* = 6.2 Hz, 1H), 7.29 – 7.23 (m, 1H), 6.78 (s, 1H), 5.51 – 5.42 (m, 1H), 4.39 – 4.29 (m, 1H), 4.19 (q, *J* = 8.0 Hz, 1H), 4.11 (q, *J* = 7.6 Hz, 1H), 3.14

(p, $J$ = 6.6 Hz, 2H), 2.16 – 2.05 (m, 2H), 2.03 – 1.91 (m, 1H), 1.86 (s, 4H), 1.80 – 1.53 (m, 5H), 1.48 – 1.34 (m, 1H), 1.26 – 1.15 (m, 3H), 1.15 – 1.03 (m, 1H), 0.87 – 0.73 (m, 6H).

**MM3177 (16) or Ac-IQSR-kbt**

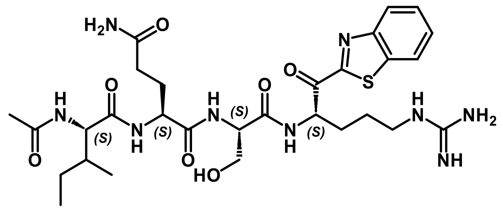

**(2S)-2-((2S)-2-acetamido-3-methylpentanamido)-N1-((S)-1-(((S)-1-(benzo[d]thiazol-2-yl)-5-guanidino-1-oxopentan-2-yl)amino)-3-hydroxy-1-oxopropan-2-yl)pentanediamide**

Yield: 28 mg (31 %). ESI-MS $[M+H]^+$ calcd. $C_{29}H_{44}N_9O_7S$ 662.79, found 662.5.

$^1$H NMR (399 MHz, dmso) δ 8.51 – 8.43 (m, 1H), 8.32 – 8.23 (m, 2H), 8.10 (d, $J$ = 7.8 Hz, 1H), 7.99 – 7.86 (m, 1H), 7.81 (d, $J$ = 7.4 Hz, 1H), 7.74 – 7.63 (m, 2H), 7.46 (t, $J$ = 5.8 Hz, 1H), 7.33 – 7.20 (m, 1H), 6.80 (s, 1H), 5.52 (dd, $J$ = 8.2, 3.9 Hz, 1H), 4.86 (s, 1H), 4.35 (p, $J$ = 5.6 Hz, 1H), 4.25 (td, $J$ = 8.4, 5.3 Hz, 1H), 4.11 (t, $J$ = 7.8 Hz, 1H), 3.62 – 3.49 (m, 2H), 3.14 (q, $J$ = 6.8 Hz, 2H), 2.15 – 2.05 (m, 2H), 2.00 – 1.81 (m, 5H), 1.79 – 1.64 (m, 3H), 1.60 (d, $J$ = 6.2 Hz, 2H), 1.47 – 1.37 (m, 1H), 1.19 – 1.03 (m, 1H), 0.85 – 0.73 (m, 6H).

**MM3194A (17) or Ac-PSKR-kbt**

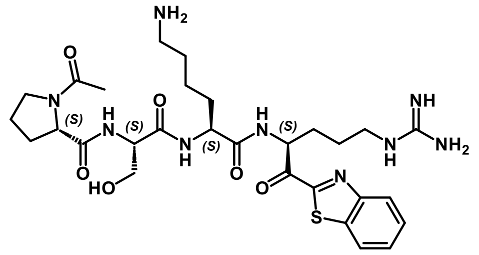

**(S)-1-acetyl-N-((S)-1-(((S)-6-amino-1-(((S)-1-(benzo[d]thiazol-2-yl)-5-guanidino-1-oxopentan-2-yl)amino)-1-oxohexan-2-yl)amino)-3-hydroxy-1-oxopropan-2-yl)pyrrolidine-2-carboxamide**

Yield: 10 mg (12 %). ESI-MS $[M+H]^+$ calcd. $C_{29}H_{44}N_9O_6S$ 646.79, found 646.5.

$^1$H NMR (600 MHz, DMSO) δ 8.37 (d, $J$ = 6.5 Hz, 1H), 8.27 (m, 2H), 8.03 – 7.90 (m, 1H), 7.80 (d, $J$ = 8.1 Hz, 1H), 7.74 – 7.62 (m, 5H), 7.58 – 7.48 (m, 1H), 5.51 – 5.43 (m, 1H), 5.25 – 4.91 (m, 1H), 4.48 – 4.15 (m, 3H), 3.72 – 3.43 (m, 4H), 3.21 – 3.04 (m, 2H), 2.78 – 2.68 (m, 2H), 2.25 – 1.80 (m, 8H), 1.80 – 1.42 (m, 8H), 1.40 – 1.15 (m, 3H).

**MM3144 (18) or Ac-QFR-kbt**

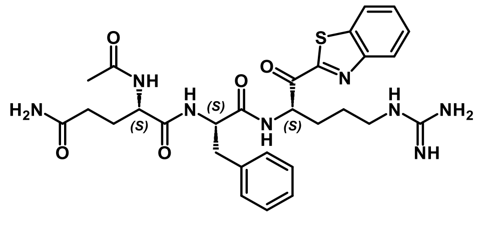

**N-[(S)-1-{N-(S)-1-[(1,3-benzothiazol-2-yl)carbonyl]-4-guanidinobutylcarbamoyl}.-2-phenylethyl](S)-2-acetylaminoglutaramide** or **Ac-QFR-kbt (17)**

Yield: 29 mg (38 %). ESI-MS $[M+H]^+$ calcd. $C_{29}H_{37}N_8O_5S$ 609.725, found 609.5.

$^1$H NMR (400 MHz, MeOD) δ 8.54 (d, *J* = 8.2 Hz, 1H), 8.26 – 8.18 (m, 2H), 8.14 (d, *J* = 7.4 Hz, 1H), 7.71 – 7.59 (m, 2H), 7.22 – 7.07 (m, 5H), 5.79 – 5.65 (m, 1H), 4.65 (q, *J* = 7.4 Hz, 1H), 4.31 – 4.23 (m, 1H), 3.29 – 3.19 (m, 1H), 3.11 (dd, *J* = 13.9, 6.5 Hz, 1H), 2.96 (dd, *J* = 14.1, 8.6 Hz, 1H), 2.28 – 2.19 (m, 2H), 2.20 – 2.08 (m, 1H), 2.02 – 1.92 (m, 4H), 1.91 – 1.79 (m, 2H), 1.79 – 1.70 (m, 2H).

### Synthesis of capped tripeptidyl ketobenzothiazoles, 19-25

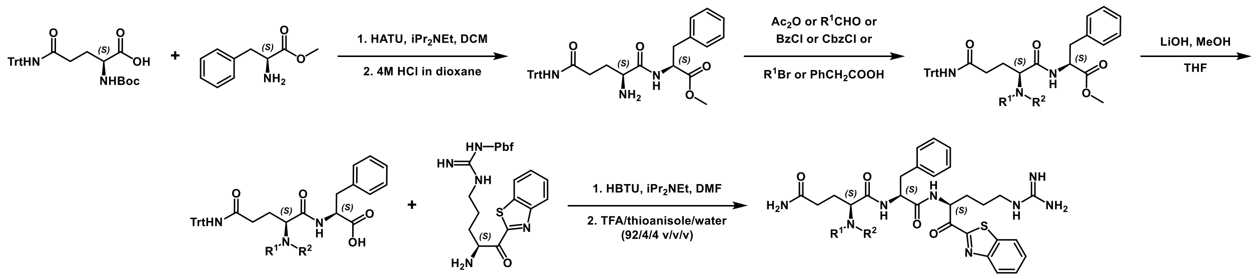


*Scheme S2. Synthesis of capped tripeptidyl ketobenzothiazoles, 19-25.*

As shown in Scheme S2Scheme S1. Synthesis of tetrapeptide ketobenzothiazoles (kbts) 1-18., we first construct the H2N-Q(Trt)-F-COOMe peptide using standard coupling protocols including HATU for amide bond coupling and 4M HCl in dioxane solution for Boc deprotection. The peptide was then capped with the corresponding acyl or alkyl group. Next, deprotection of methyl ester was performed by lithium hydroxide followed by H-Arg(Pbf)-kbt introduced with HBTU in DMF. Final deprotection of the amino acid sidechains using TFA:water:thioanisole (95:2.5:2.5 %v/v) generates the target compounds which were purified by reverse-phase prep HPLC.

**CA1041 (19) or Dimethyl-QFR-kbt**

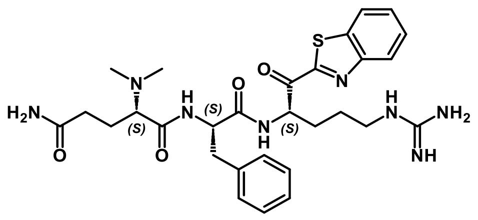

**N-[(S)-1-{N-(S)-1-[(1,3-benzothiazol-2-yl)carbonyl]-4-guanidinobutylcarbamoyl}-2-phenylethyl](S)-2-(dimethylamino)glutaramide**

Into a round bottom flask containing H2N-Q(Trt)-F-COOMe (300 mg, 0.54 mmol) and formaldehyde (0.5 ml) dissolved in acetonitrile (4 ml) was added sodium cyanoborohydride (303 mg, 4.8 mmol) and stirred for 2 h. The solvent was removed and dried in a vacuum. EtOAc and water were added to the residue, the organic layer was washed with brine, dried with $MgSO_4$, filtered, and concentrated. To the residue, 2M LiOH solution in water (0.312 ml, 0.63 mmol), 1.5 ml of MeOH, and 1.5 ml of THF were added. The flask was stirred for 3 h under argon and the solvents were removed under reduced pressure. The crude peptide acid residue (270 mg, 0.48 mmol) was dissolved in DMF (8 ml). To the solution under argon atmosphere at 0°C was added HBTU (200 mg, 0.52 mmol) and stirred for 5 min. Next, Arg(Pbf)-kbt×HCl (260 mg, 0.48 mmol) and iPr$_2$NEt (0.17 ml, 1.0 mmol) were added to the reaction and stirred overnight. The solvent was removed in a vacuum. To the residue, 10 ml of TFA/thioanisole/water mixture (92/4/4 v/v/v) was added and stirred for 3 h. The solvent was evaporated, and the residue was dried in a vacuum. The crude product was purified by HPLC to give the final compound.

Yield: 13 mg (4%), ESI-MS $[M+H]^+$ calcd. $C_{29}H_{39}N_8O_4S$ 595.742, found 595.91

$^1$H NMR (399 MHz, cd$_3$od) δ 8.27 – 8.19 (m, 1H), 8.19 – 8.12 (m, 1H), 7.75 – 7.59 (m, 2H), 7.35 – 7.11 (m, 6H), 5.76 – 5.66 (m, 1H), 4.96 (dd, *J* = 10.9, 4.7 Hz, 1H), 3.62 (dd, *J* = 9.9, 3.7 Hz, 1H), 3.30 – 3.24 (m, 3H), 3.19 (s, 1H), 2.92 – 2.80 (m, 2H), 2.79 – 2.35 (m, 3H), 2.34 – 2.14 (m, 4H), 2.06 – 1.72 (m, 5H).

**CA1033 (20) or PhCH$_2$CH$_2$-QFR-kbt**

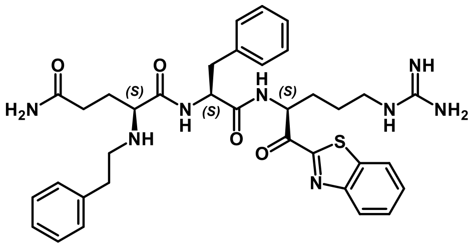


**N-[(S)-1-{N-(S)-1-[(1,3-benzothiazol-2-yl)carbonyl]-4-guanidinobutylcarbamoyl}-2-phenylethyl](S)-2-(phenethylamino)glutaramide**

Phenylacetaldehyde (180 mg, 1.5 mmol) and sodium triacetoxyborohydride (975 mg, 4.6 mmol) were added to the solution of H$_2$N-Q(Trt)-F-COOMe peptide (900 mg, 1.5 mmol) dissolved in DCE (10 ml) and stirred overnight. The intermediate was purified by silica gel chromatography using DCM/MeOH gradient as eluent. The residue (166 mg, 0.26 mmol) was redissolved in anhydrous THF (5 ml) under an inert atmosphere, Boc anhydride (0.18 ml, 0.78 mmol) was added to the solution followed by the Et$_3$N (0.20 ml, 1.4 mmol). The mixture was stirred at 60°C overnight and then cooled to rt. LiOH (84 mg, 2 mmol) was added to the mixture, followed by the addition of $H_2O$ (1.5 ml) and MeOH (3 ml). The mixture was stirred at rt and monitored by LCMS until completion. The mixture was concentrated, followed by adding 0.5 M HCl aqueous solution at 0°C to pH 5. The product was extracted with ethyl acetate, the organic layer was washed with water, dried over sodium sulfate and the solvent was removed in a vacuum. To the residue anhydrous DMF (5 ml) was added, followed by HATU (116 mg, 0.31 mmol) at 0°C. The mixture was stirred for 15 min. Then Arg(Pbf)-kbt×HCl (0.177 g, 0.31 mmol) was added followed by iPr$_2$NEt (0.22 ml, 1.27 mmol). The mixture was allowed to warm up to rt for 2 h. The solvent was removed by evaporation, water (20 ml) was added, and the mixture was sonicated. The precipitate was filtered, washed with water, and then dried in a vacuum. 10 ml of TFA/thioanisole/water mixture (92/4/4 v/v/v) was added to the residue and stirred for 3 h. The solvent was evaporated, and the residue was dried in a vacuum. The crude product was purified by HPLC to give the final compound.

Yield: 29 mg (25 %). ESI-MS $[M+H]^+$ calcd. $C_{35}H_{42}N_8O_4S$ 670.83, found 671.3

$^1$H NMR (399 MHz, cd$_3$od) δ 8.23 – 8.20 (m, 1H), 8.14 – 8.11 (m, 1H), 7.68 – 7.60 (m, 2H), 7.38 – 7.33 (m, 2H), 7.33 – 7.21 (m, 6H), 7.20 – 7.16 (m, 2H), 7.13 – 7.08 (m, 2H), 6.90 (t, *J* = 7.2 Hz, 1H), 5.63 (dd, *J* = 9.6, 4.1 Hz, 1H), 3.78 – 3.72 (m, 1H), 3.25 – 3.14 (m, 4H), 2.91 – 2.80 (m, 4H),

2.63 – 2.56 (m, 1H), 2.42 – 2.33 (m, 2H), 2.20 – 2.11 (m, 1H), 2.08 – 1.97 (m, 3H), 1.86 – 1.73 (m, 2H), 1.66 – 1.57 (m, 2H).

**CA1022 (21) or Bn-QFR-kbt**

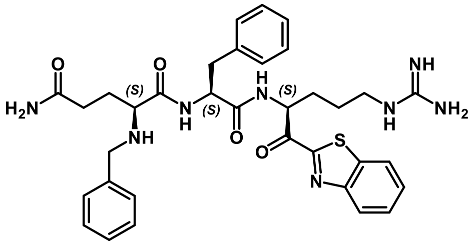

**N-[(S)-1-{N-(S)-1-[(1,3-benzothiazol-2-yl)carbonyl]-4-guanidinobutylcarbamoyl}-2-phenylethyl](S)-2-benzylaminoglutaramide**

Into a round bottom flask containing $H_2N$-Q(Trt)-F-COOMe (200 mg, 0.34 mmol) dissolved in DMF (5 ml) was added potassium carbonate (94 mg, 0.68 mmol) and stirred overnight at 70°C. The solvent was removed by evaporation and the residue was dried in a vacuum. To the intermediate 2M LiOH water solution (0.312 ml, 0.63 mmol) was added followed by 3 ml of MeOH/THF 1/1 mixture. The reaction mixture was stirred for 3 h under an inert atmosphere and then concentrated in a vacuum. The residue was dissolved in 40 ml of water and 1M HCl was added dropwise at 0ºC to pH 3. The reaction mixture was stirred for 30 min, precipitate was collected by filtration and dried in a vacuum. The crude peptide acid residue (150 mg, 0.24 mmol) was dissolved in DMF (4 ml). To the solution under argon atmosphere at 0°C was added HBTU (100 mg, 0.26 mmol) and stirred for 5 min. Next, Arg(Pbf)-kbt×HCl (130 mg, 0.24 mmol) and $iPr_2NEt$ (0.08 ml, 0.5 mmol) were added to the reaction and stirred overnight. The solvent was removed in a vacuum. To the residue, 10 ml of TFA/thioanisole/water mixture (92/4/4 v/v/v) was added and stirred for 3 h. The solvent was evaporated, and the residue was dried in a vacuum. The crude product was purified by HPLC to give the final compound.

Yield: 110 mg ( 49%). ESI-MS $[M+H]^+$ calcd. $C_{34}H_{41}N_8O_4S$ 657.81, found 657.3

1H NMR (399 MHz, dmso) δ 9.03 – 8.94 (m, 2H), 8.33 – 8.23 (m, 2H), 7.74 – 7.61 (m, 3H), 7.54 (s, 1H), 7.41 – 7.15 (m, 12H), 7.14 – 7.06 (m, 3H), 5.57 – 5.45 (m, 1H), 4.96 – 4.85 (m, 1H), 3.80 – 3.64 (m, 1H), 3.60 – 3.55 (m, 1H), 3.44 – 3.28 (m, 2H), 2.69 (t, $J$ = 12.7 Hz, 1H), 2.19 (t, $J$ = 7.2 Hz, 2H), 2.10 – 1.89 (m, 1H), 1.90 – 1.75 (m, 2H), 1.66 – 1.61 (m, 1H).

**CA1018 (22) or PhAc-QFR-kbt**

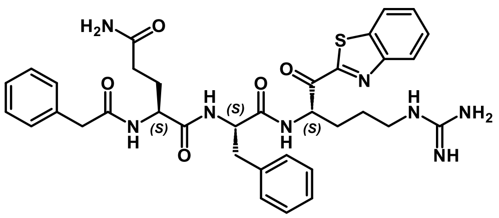

**N-[(S)-1-{N-(S)-1-[(1,3-benzothiazol-2-yl)carbonyl]-4-guanidinobutylcarbamoyl}-2-phenylethyl](S)-2-(benzylcarbonylamino)glutaramide**

Into a round bottom flask containing $H_2N$-Q(Trt)-F-COOMe (300 mg, 0.52 mmol) and HBTU (213 mg, 0.56 mmol) in DMF (2.6 ml) phenylacetic acid (70 mg, 0.52 mmol) and $iPr_2NEt$

(0.186 ml, 1.1 mmol) at 0°C were added, and stirred overnight under argon atmosphere. The mixture was concentrated under a vacuum. To the residue EtOAc and water were added, water was washed with EtOAc, and combined organic layers were washed with brine, dried with $MgSO_4$, and concentrated under vacuum. The intermediate was purified by silica gel chromatography (eluent: DCM/MeOH). To the intermediate 2M LiOH water solution (0.36 ml, 0.72 mmol) was added followed by 3 ml of MeOH/THF 1/1 mixture. The reaction mixture was stirred for 3 h under an inert atmosphere and then concentrated in a vacuum. The residue was dissolved in 40 ml of water and 1M HCl was added dropwise at 0°C to pH 3. The reaction mixture was stirred for 30 min, precipitate was collected by filtration and dried in a vacuum. The crude peptide acid residue (157 mg, 0.24 mmol) was dissolved in DMF (4 ml). To the solution under argon atmosphere at 0°C was added HBTU (100 mg, 0.26 mmol) and stirred for 5 min. Next, Arg(Pbf)-kbt×HCl (130 mg, 0.24 mmol) and $iPr_2NEt$ (0.08 ml, 0.5 mmol) were added to the reaction and stirred overnight. The solvent was removed in a vacuum. To the residue, 10 ml of TFA/thioanisole/water mixture (92/4/4 v/v/v) was added and stirred for 3 h. The solvent was evaporated, and the residue was dried in a vacuum. The crude product was purified by HPLC to give the final compound.

Yield: 60 mg (17%). ESI-MS $[M+H]^+$ calcd. $C_{35}H_{41}N_8O_5S$ 685.823, found 685.78

$^1H$ NMR (399 MHz, dmso) δ 8.60 (dd, $J$ = 18.4, 6.7 Hz, 1H), 8.25 – 8.21 (m, 2H), 8.22 – 8.14 (m, 1H), 8.00 (t, $J$ = 8.4 Hz, 1H), 7.68 – 7.61 (m, 2H), 7.47 – 7.36 (m, 1H), 7.27 – 7.05 (m, 13H), 6.78 – 6.71 (m, 1H), 5.51 – 5.37 (m, 1H), 4.63 – 4.51 (m, 1H), 4.18 – 4.08 (m, 1H), 3.13 – 3.00 (m, 2H), 2.99 – 2.87 (m, 1H), 2.79 – 2.62 (m, 1H), 2.02 – 1.93 (m, 2H), 1.79 – 1.67 (m, 2H), 1.63 – 1.50 (m, 3H), 1.45 – 1.36 (m, 1H).

**CA1043 (23) or Bz-QFR-kbt**

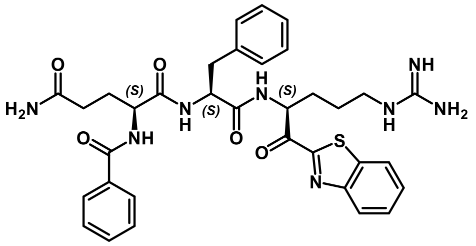


**N-[(S)-1-{N-(S)-1-[(1,3-benzothiazol-2-yl)carbonyl]-4-guanidinobutylcarbamoyl}-2-phenylethyl](S)-2-benzylaminoglutaramide**

Into a round bottom flask containing $H_2N$-Q(Trt)-F-COOMe (300 mg, 0.52 mmol) and HBTU (213 mg, 0.56 mmol) in DMF (2.6 ml) benzoic acid (63 mg, 0.52 mmol) and $iPr_2NEt$ (0.186 ml, 1.1 mmol) at 0°C were added, and stirred overnight under argon atmosphere. The mixture was concentrated under a vacuum. To the residue EtOAc and water were added, water was washed with EtOAc, and combined organic layers were washed with brine, dried with $MgSO_4$, and concentrated under vacuum. To the intermediate 2M LiOH water solution (0.36 ml, 0.72 mmol) was added followed by 3 ml of MeOH/THF 1/1 mixture. The reaction mixture was stirred for 3 h under an inert atmosphere and then concentrated in a vacuum. The residue was dissolved in 40 ml of water and 1M HCl was added dropwise at 0°C to pH 3. The reaction mixture was stirred for 30 min, precipitate was collected by filtration and dried in a vacuum. The crude peptide acid residue (350 mg, 0.54 mmol) was dissolved in DMF (10 ml). To the solution under argon atmosphere at 0°C was added HBTU (228 mg, 0.6 mmol) and stirred for 5 min. Next, Arg(Pbf)-kbt×HCl (297 mg, 0.54 mmol) and $iPr_2NEt$ (0.2 ml, 1.1 mmol) were added to the reaction and stirred overnight. The solvent was removed in a vacuum. To the residue, 10 ml of TFA/thioanisole/water mixture (92/4/4 v/v/v) was added and stirred for 3 h. The solvent was evaporated, and the residue was dried in a vacuum. The crude product was purified by HPLC to give the final compound.

Yield: 48 mg (40%), ESI-MS $[M+H]^+$ calcd. for $C_{34}H_{39}N_8O_5S$ 671.796, found 671.931

$^1$H NMR (399 MHz, dmso) δ 8.68 – 8.52 (m, 2H), 8.28 – 8.20 (m, 2H), 8.00 (t, *J* = 8.7 Hz, 1H), 7.84 (t, *J* = 7.6 Hz, 2H), 7.70 – 7.60 (m, 2H), 7.50 (d, *J* = 7.3 Hz, 1H), 7.43 (t, *J* = 7.3 Hz, 3H), 7.29 (s, 1H), 7.18 – 7.13 (m, 3H), 7.11 – 7.07 (m, 2H), 6.83 (s, 1H), 5.49 – 5.43 (m, 1H), 4.62 – 4.54 (m, 1H), 4.32 – 4.26 (m, 1H), 3.14 – 3.03 (m, 2H), 3.01 – 2.90 (m, 1H), 2.82 – 2.72 (m, 1H), 2.15 – 1.99 (m, 2H), 1.99 – 1.82 (m, 2H), 1.81 – 1.69 (m, 2H), 1.65 – 1.42 (m, 2H).

**CA1046-2 (24) or Pip-QFR-kbt**

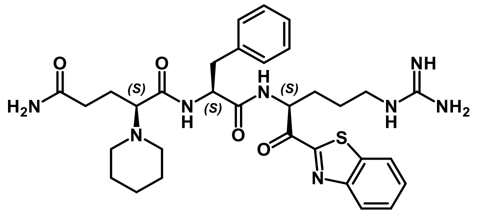

**N-[(S)-1-{N-(S)-1-[(1,3-benzothiazol-2-yl)carbonyl]-4-guanidinobutylcarbamoyl}-2-phenylethyl](S)-2-piperidinoglutaramide**

Into a round bottom flask containing $H_2N$-Q(Trt)-F-COOMe (260 mg, 0.37 mmol) and 1,5-dibromopentane (340 mg, 0.74 mmol) dissolved in ethanol (100 ml) potassium carbonate (420 mg, 1.5 mmol) was added and the reaction mixture was stirred under reflux for 24 h. Potassium carbonate was filtered off, and the solvent was removed by evaporation and the residue was dried in a vacuum. The intermediate was purified by silica gel chromatography (eluent: EtOAc/hexane). To the intermediate 2M LiOH water solution (0.16 ml, 0.31 mmol) was added followed by 3 ml of MeOH/THF 1/1 mixture. The reaction mixture was stirred for 3 h under an inert atmosphere and then concentrated in a vacuum. The crude peptide acid residue (160 mg, 0.27 mmol) was dissolved in DMF (4.5 ml). To the solution under argon atmosphere at 0°C was added HBTU (111 mg, 0.29 mmol) and stirred for 5 min. Next, Arg(Pbf)-kbt×HCl (144 mg, 0.27 mmol) and $iPr_2NEt$ (0.1 ml, 0.56 mmol) were added to the reaction and stirred overnight. The solvent was removed in a vacuum. To the residue, 10 ml of TFA/thioanisole/water mixture (92/4/4 v/v/v) was added and stirred for 3 h. The solvent was evaporated, and the residue was dried in a vacuum. The crude product was purified by HPLC to give the final compound.

Yield: 10 mg (15 %). ESI-MS $[M+H]^+$ calcd. $C_{32}H_{43}N_8O_4S$ 635.807, found 635.3

$^1$H NMR (399 MHz, $cd_3od$) δ 8.27 – 8.19 (m, 1H), 8.19 – 8.11 (m, 1H), 7.71 – 7.59 (m, 2H), 7.38 – 7.23 (m, 5H), 7.20 (m, 1H), 5.71 (td, *J* = 9.7, 4.3 Hz, 1H), 5.02 (dd, *J* = 11.3, 4.7 Hz, 0.6H), 4.79 (dd, *J* = 10.9, 5.1 Hz, 0.4H), 3.84 (dd, *J* = 9.3, 4.3 Hz, 0.4H), 3.71 – 3.35 (m, 3H), 3.28 (m, 2H), 3.11 – 2.77 (m, 2H), 2.77 – 2.39 (m, 1H), 2.39 – 2.12 (m, 3H), 2.12 – 1.25 (m, 14H).

**ZFH9141 (25) or Cbz-QFR-kbt**

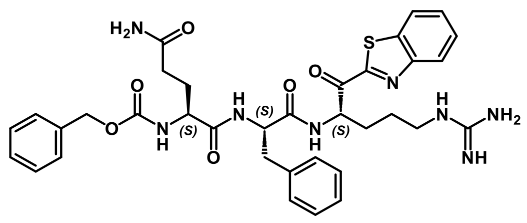

**(S)-1,3-bis[N-(S)-1-{N-(S)-1-[(1,3-benzothiazol-2-yl)carbonyl]-4-guanidinobutylcarbamoyl}-2-phenylethylcarbamoyl]propyl phenylmethanecarbamate**

Into a round bottom flask containing H$_2$N-Q(Trt)-F-COOMe (200 mg, 0.34 mmol) and Et$_3$N (94 µl, 0.68 mmol) dissolved in THF (5 ml), Cbz-Cl (53 µl, 0.37 mmol) was added at 0°C. Conversion was monitored by LCMS. The reaction mixture was stirred for 1h. Then. 2 ml of water was added to quench the reaction, and the mixture was stirred for 30 min. LiOH (24 mg, 0.1 mmol) and MeOH (2 ml) were added to the reaction mixture and stirred at rt for 1h. After that, the second portion of LiOH (12 mg, 0.05 mmol), water (1 ml), and methanol (2 ml) were added. After 1h 1M HCl solution was added dropwise at 0°C to pH 5. Precipitate was formed. THF and MeOH were removed under reduced pressure, the precipitate was filtered, washed with water, and dried in a high vacuum. The crude peptide acid residue (187 mg, 0.28 mmol) was dissolved in DMF (5 ml). To the solution under argon atmosphere at 0°C was added HATU (127 mg, 0.34 mmol) and stirred for 5 min. Next, Arg(Pbf)-kbt×HCl (194 mg, 0.34 mmol) and iPr$_2$NEt (0.24 ml, 1.4 mmol) were added to the reaction and stirred overnight. The solvent was removed in a vacuum. To the residue, 10 ml of TFA/thioanisole/water mixture (92/4/4 v/v/v) was added and stirred for 30 min. The solvent was evaporated, and the residue was dried in a vacuum. The crude product was purified by HPLC to give the final compound.

Yield: 88 mg (32%). ESI-MS $[M+H]^+$ calcd. for $C_{35}H_{41}N_8O_6S$ 701.822, found 701.3.

$^1$H NMR (399 MHz, cd$_3$od) δ 8.22 (d, $J$ = 7.5 Hz, 1H), 8.16 – 8.10 (m, 1H), 7.73 – 7.59 (m, 2H), 7.41 – 7.02 (m, 10H), 5.74 – 5.66 (m, 1H), 5.12 – 5.06 (m, 2H), 4.77 – 4.58 (m, 1H), 4.10 – 3.95 (m, 1H), 3.26 – 3.17 (m, 2H), 3.09 (dd, $J$ = 13.9, 6.7 Hz, 1H), 3.00 – 2.89 (m, 1H), 2.23 (q, $J$ = 6.9 Hz, 1H), 2.18 – 2.05 (m, 2H), 2.01 – 1.89 (m, 1H), 1.89 – 1.59 (m, 2H).

### Synthesis of sulfonyl-capped QFR-kbt peptides, 27-29

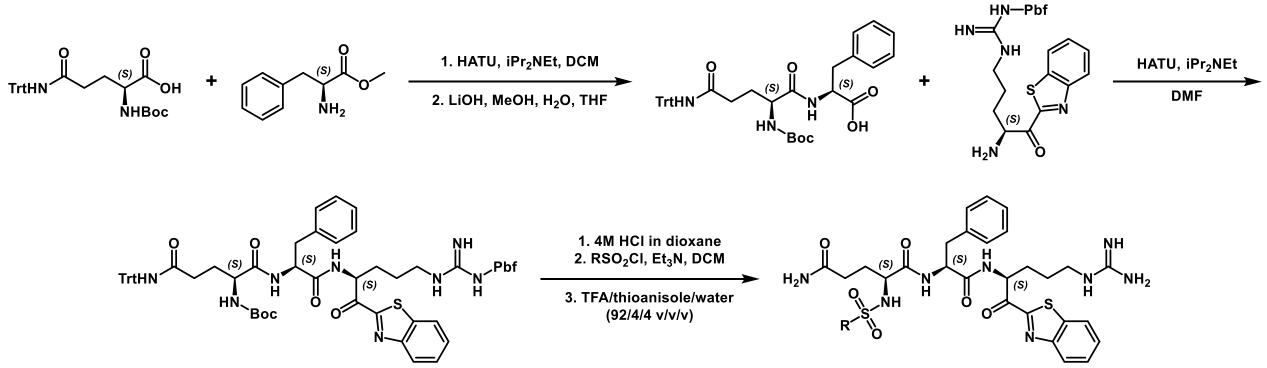


*Scheme S3. Synthesis of sulfonyl-capped QFR-kbt peptides, 27-29*

As shown in Scheme S3Scheme S1. Synthesis of tetrapeptide ketobenzothiazoles (kbts) 1-18., we first construct the Boc-H$_2$N-Q(Trt)-F-Arg(pbf)-kbt peptide using standard coupling protocols including HATU for amide bond coupling, LiOH in water/methanol/THF mixture to ester hydrolysis, and

4M HCl in dioxane solution for Boc deprotection. The peptide was then capped with the corresponding sulfonyl chloride, followed by final deprotection of the amino acid sidechains using TFA:water:thioanisole (95:2.5:2.5 %v/v) to give the target compounds which are purified by reverse-phase prep HPLC.

**MF1101 (27) or PhSO2-QFR-kbt**

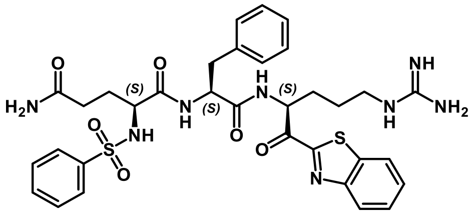

**N-[(S)-1-{N-(S)-1-[(1,3-benzothiazol-2-yl) carbonyl]-4-guanidinobutylcarbamoyl}-2-phenylethyl](S)-2-(phenylsulfonylamino)glutaramide**

Yield: 9 mg (12%). ESI-MS $[M+H]^+$ calcd. for $C_{33}H_{39}N_8O_6S_2$ 707.851, found 708.33.

$^1$H NMR (399 MHz, cd$_3$od) δ 8.38 (dd, *J* = 11.2, 7.8 Hz, 1H), 8.23 – 8.19 (m, 1H), 8.15 – 8.11 (m, 1H), 7.82 – 7.78 (m, 2H), 7.67 – 7.58 (m, 3H), 7.58 – 7.46 (m, 3H), 7.30 – 7.07 (m, 5H), 5.71 – 5.55 (m, 1H), 4.63 – 4.42 (m, 1H), 3.69 – 3.59 (m, 1H), 3.27 – 3.21 (m, 1H), 3.20 – 3.14 (m, 1H), 3.12 – 3.03 (m, 1H), 2.86 – 2.78 (m, 1H), 2.12 – 2.05 (m, 2H), 1.98 (t, *J* = 7.4 Hz, 1H), 1.83 – 1.64 (m, 5H), 1.61 – 1.50 (m, 1H).

**MF1105 (28) or BnSO2-QFR-kbt**

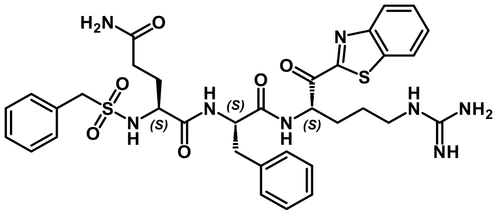

**N-[(S)-1-{N-(S)-1-[(1,3-benzothiazol-2-yl)carbonyl]-4-guanidinobutylcarbamoyl}-2-phenylethyl](S)-2-(benzylsulfonylamino)glutaramide**

Yield: 22 mg (29%). ESI-MS $[M+H]^+$ calcd. for $C_{34}H_{41}N_8O_6S_2$ 721.878, found 722.31.

$^1$H NMR (399 MHz, cd$_3$od) δ 8.48 – 8.34 (m, 1H), 8.24 – 8.20 (m, 1H), 8.13 (td, *J* = 6.9, 1.8 Hz, 1H), 7.67 – 7.61 (m, 2H), 7.35 – 7.31 (m, 6H), 7.26 – 7.21 (m, 3H), 7.14 (t, *J* = 7.4 Hz, 2H), 7.04 (t, *J* = 7.4 Hz, 1H), 5.73 – 5.56 (m, 1H), 4.78 – 4.72 (m, 1H), 4.17 (s, 1H), 4.09 (s, 1H), 3.85 – 3.78 (m, 1H), 3.26 – 3.13 (m, 4H), 2.97 – 2.88 (m, 1H), 2.25 – 2.07 (m, 4H), 1.86 – 1.73 (m, 5H), 1.68 – 1.54 (m, 1H).

**MF1104 (29) or CypSO2-QFR-kbt**

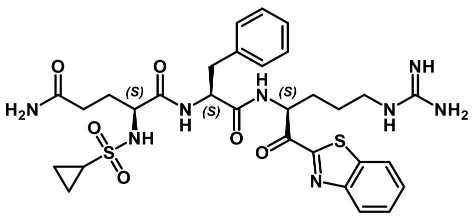

**N-[(S)-1-{N-(S)-1-[(1,3-benzothiazol-2-yl)carbonyl]-4-guanidinobutylcarbamoyl}-2-phenylethyl](S)-2-(cyclopropylsulfonylamino)glutaramide**

Yield: 15 mg (21%). ESI-MS $[M+H]^+$ calcd. for $C_{30}H_{39}N_8O_6S_2$ 671.818, found 672.24.

$^1$H NMR (399 MHz, cd$_3$od) δ 8.24 – 8.20 (m, 1H), 8.16 – 8.11 (m, 1H), 7.68 – 7.60 (m, 2H), 7.32 – 7.05 (m, 6H), 5.77 – 5.57 (m, 1H), 4.81 – 4.70 (m, 1H), 3.93 – 3.84 (m, 1H), 3.23 – 3.14 (m, 2H), 2.98 – 2.88 (m, 1H), 2.37 – 2.10 (m, 5H), 2.00 – 1.69 (m, 5H), 1.61 (p, $J$ = 7.4 Hz, 1H), 0.98 – 0.68 (m, 5H).

### NMR data:

MM3116 (1)

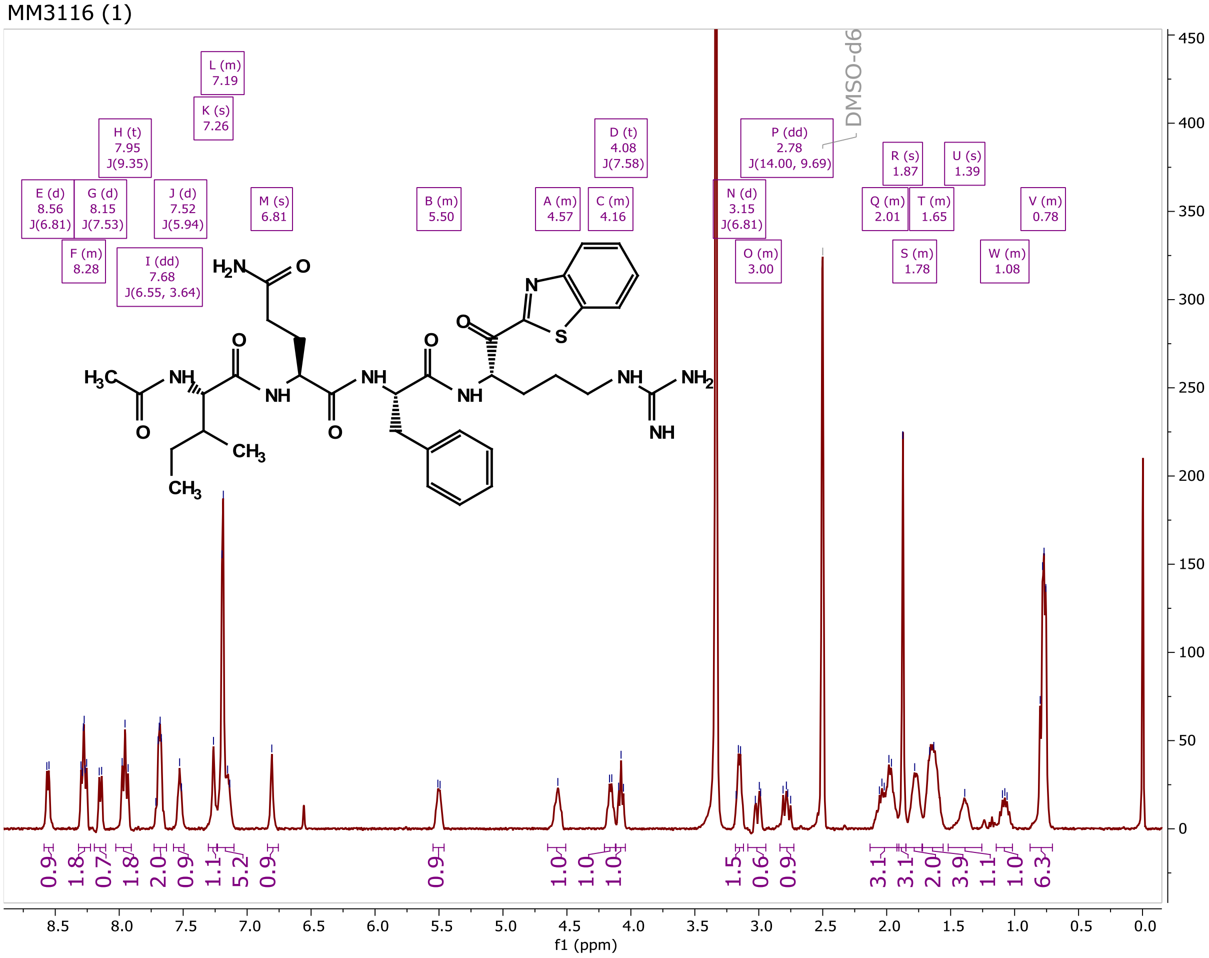

MM3122 (2)

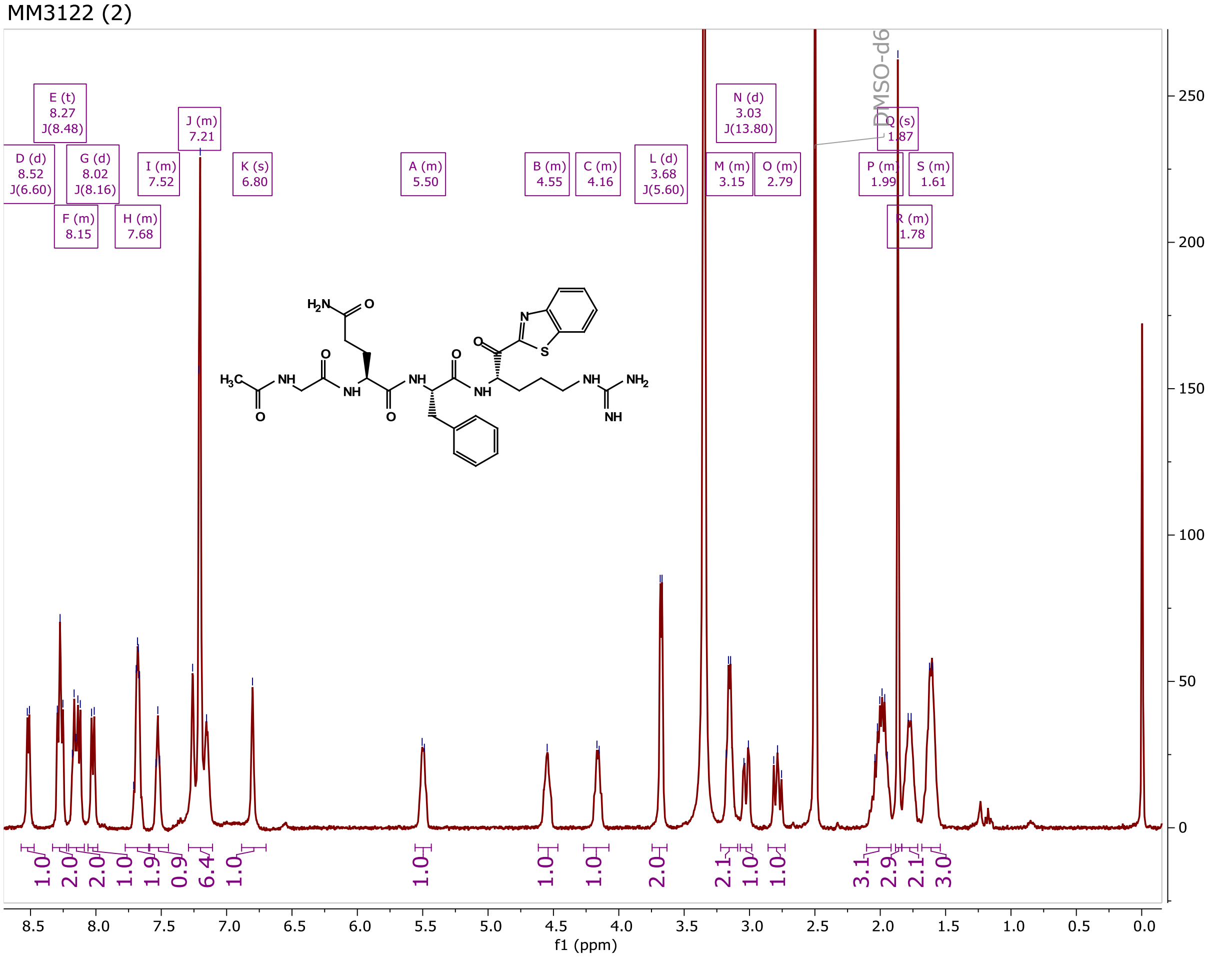


MM3123 (3)

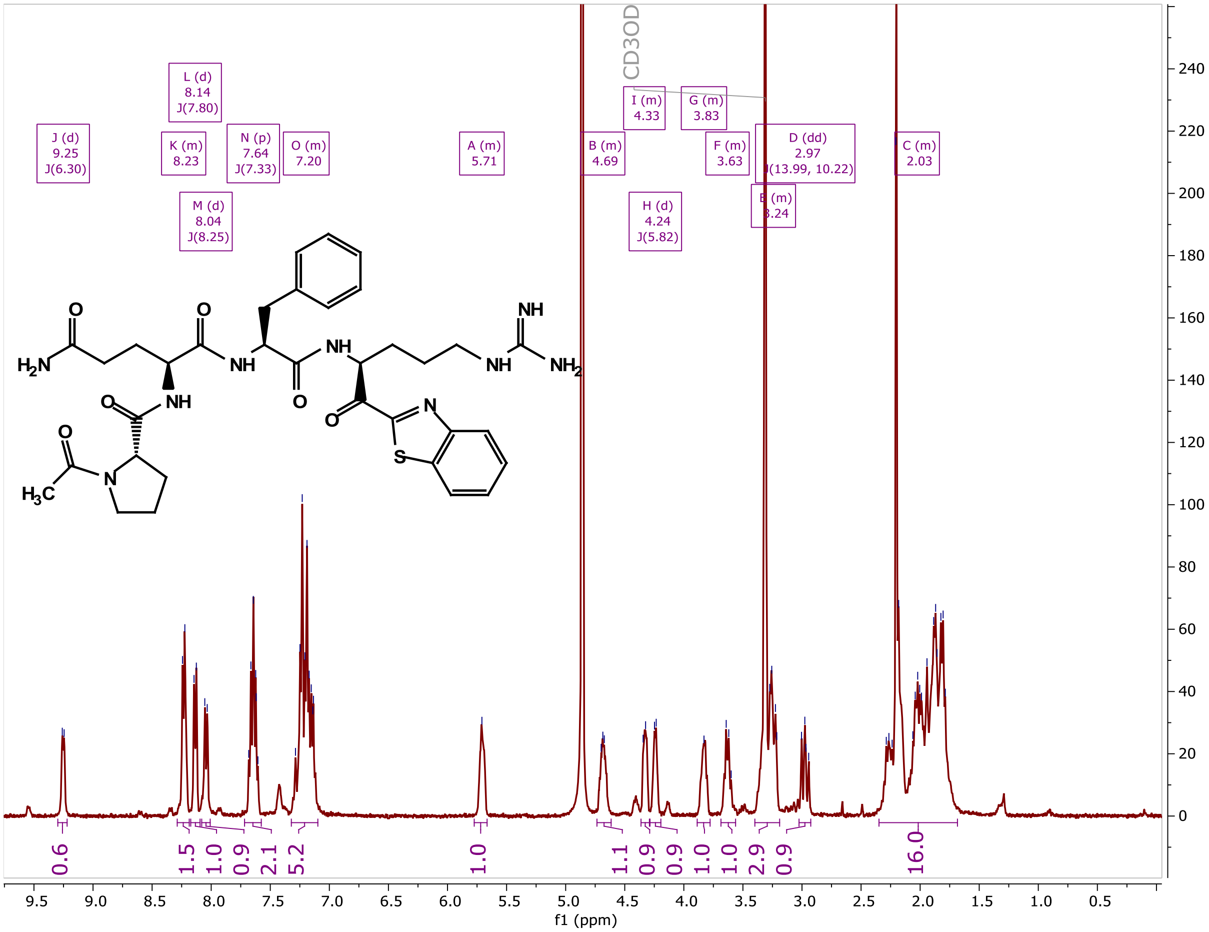

MM3131 (4)

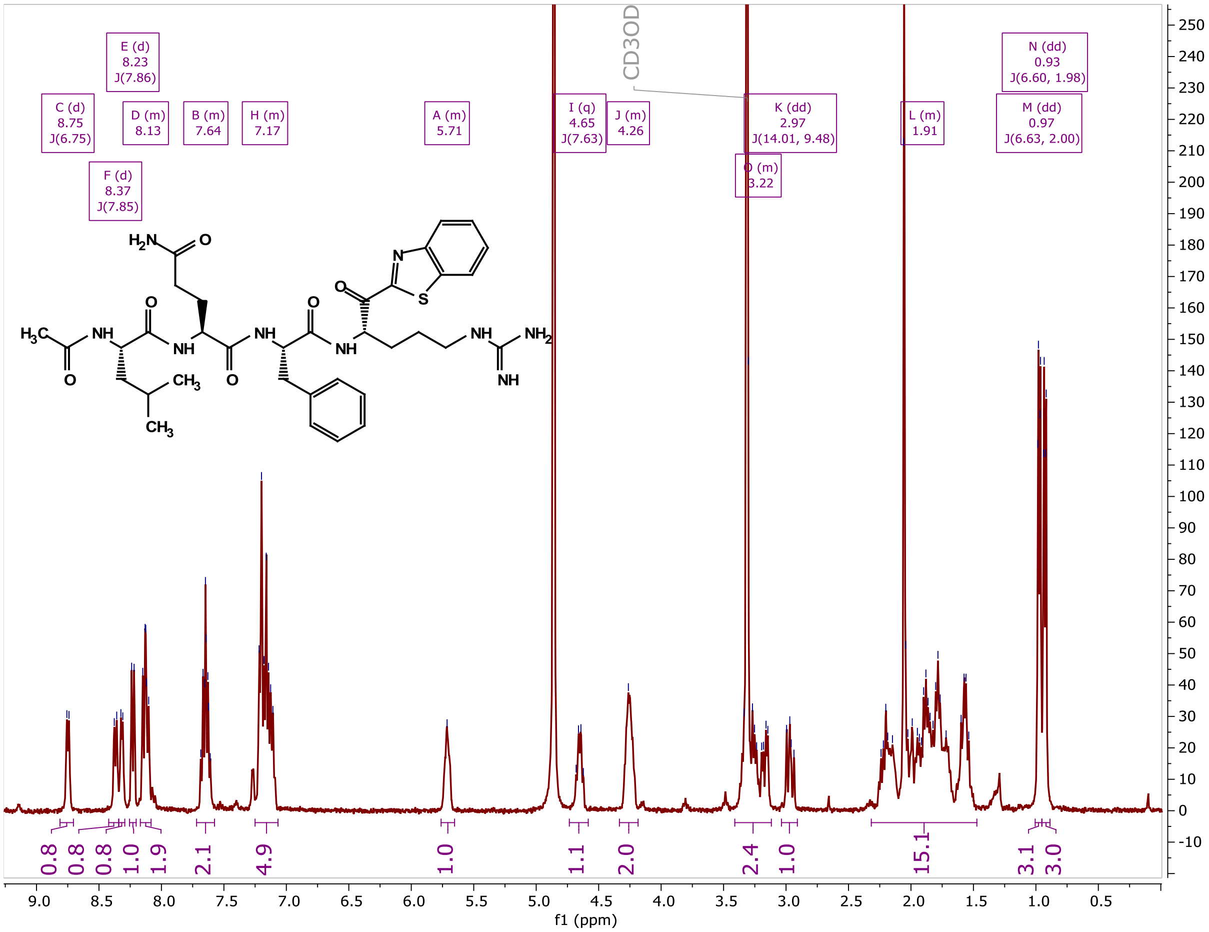


MM3130 (5)

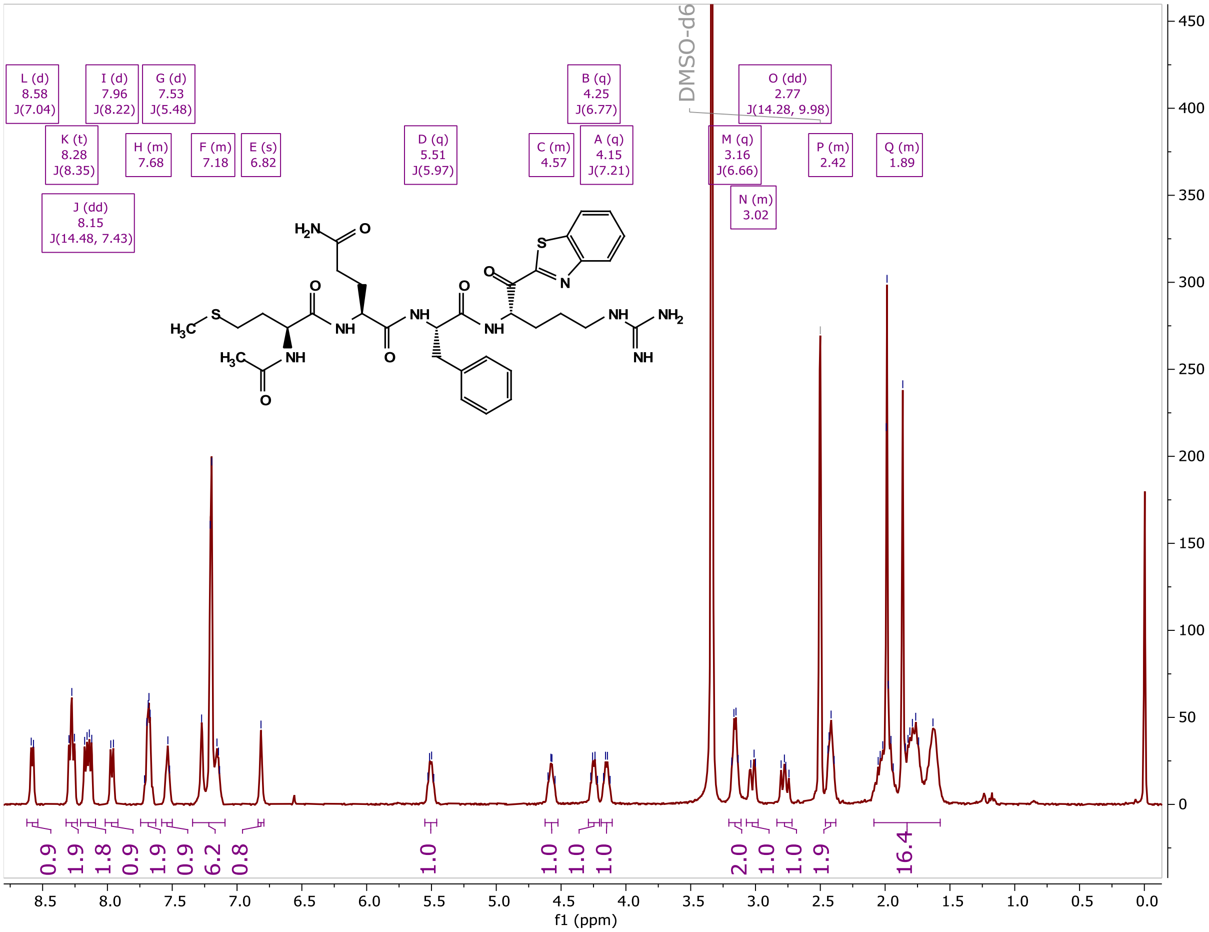

MM4037 (7)

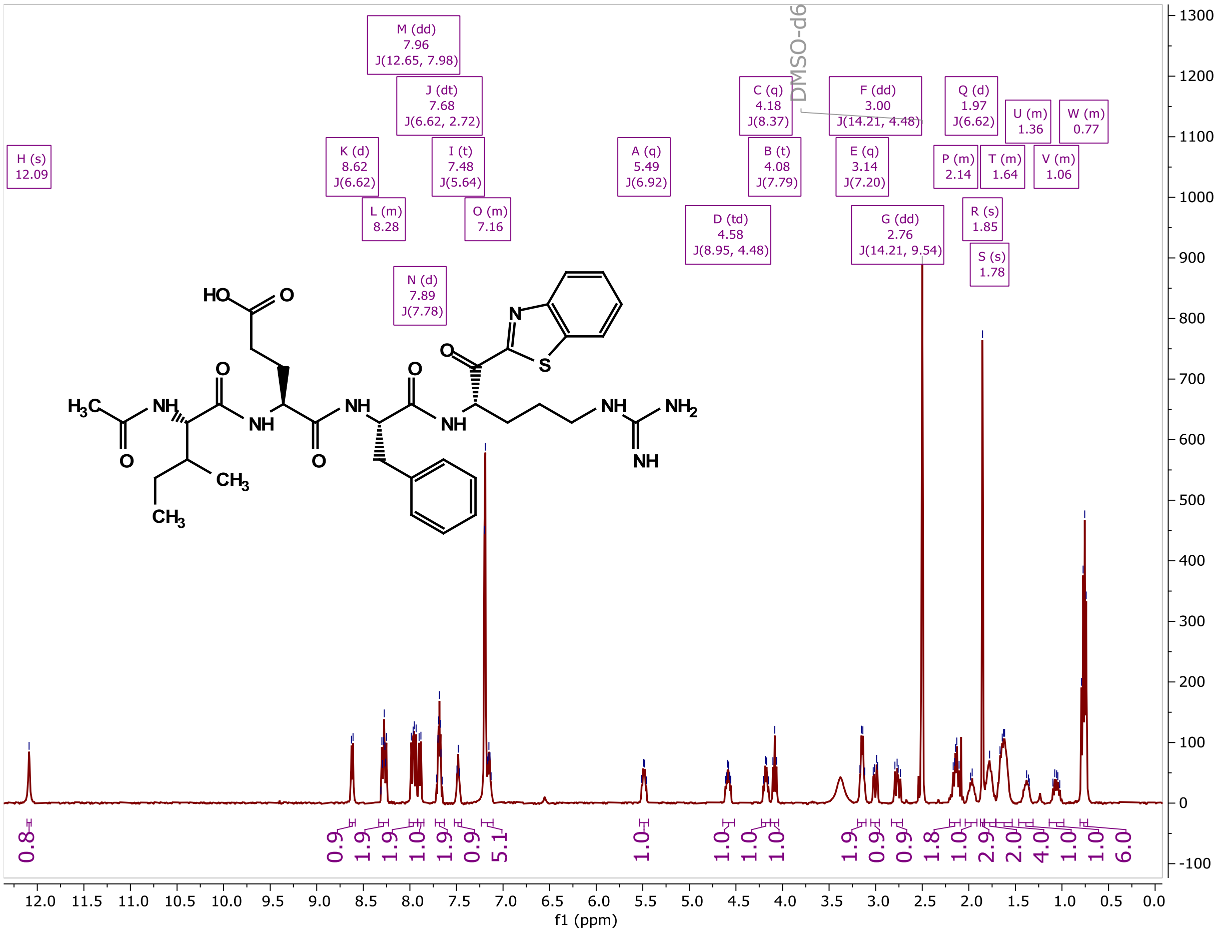


MM4028 (8)

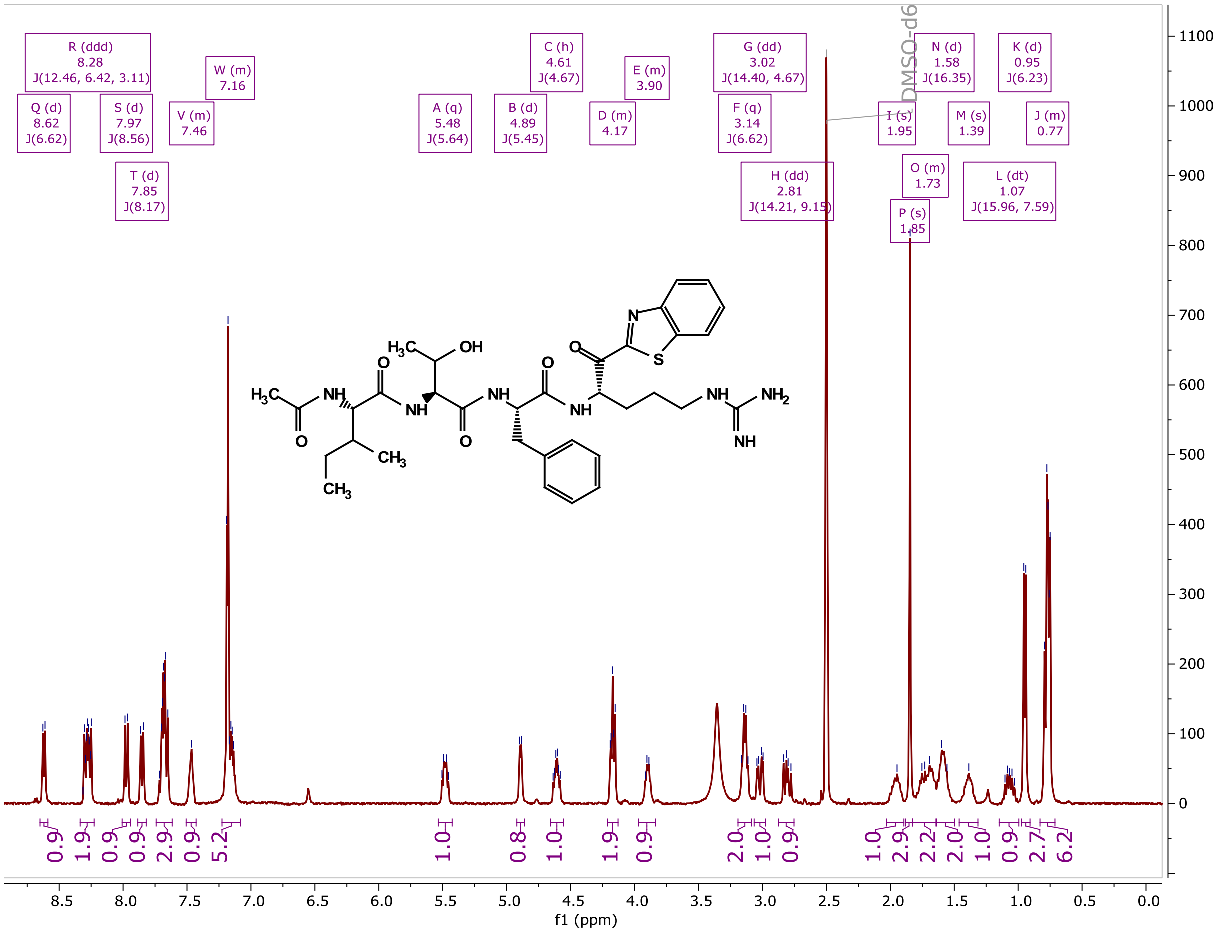

MM4027 (9)

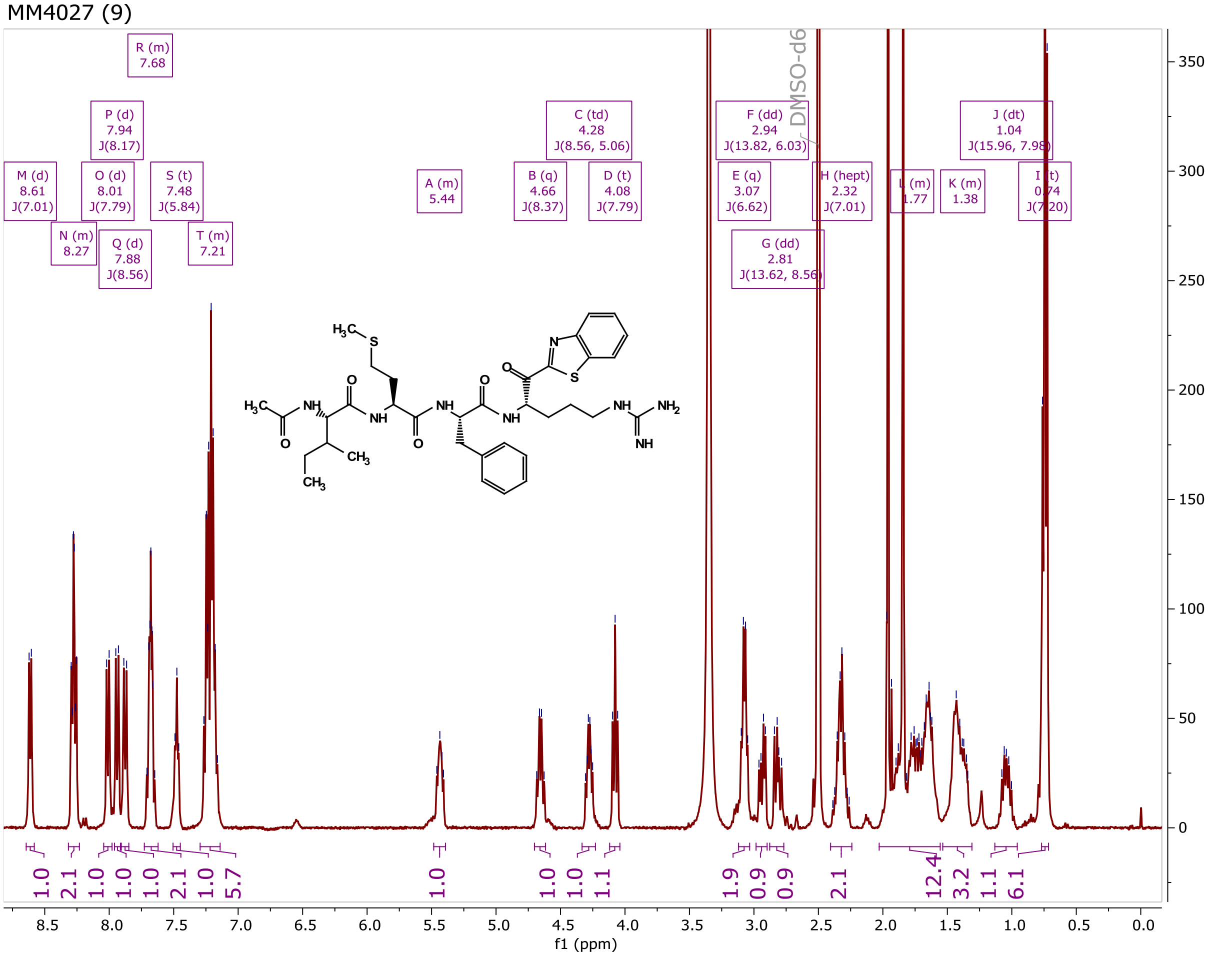


MM4009 (10)

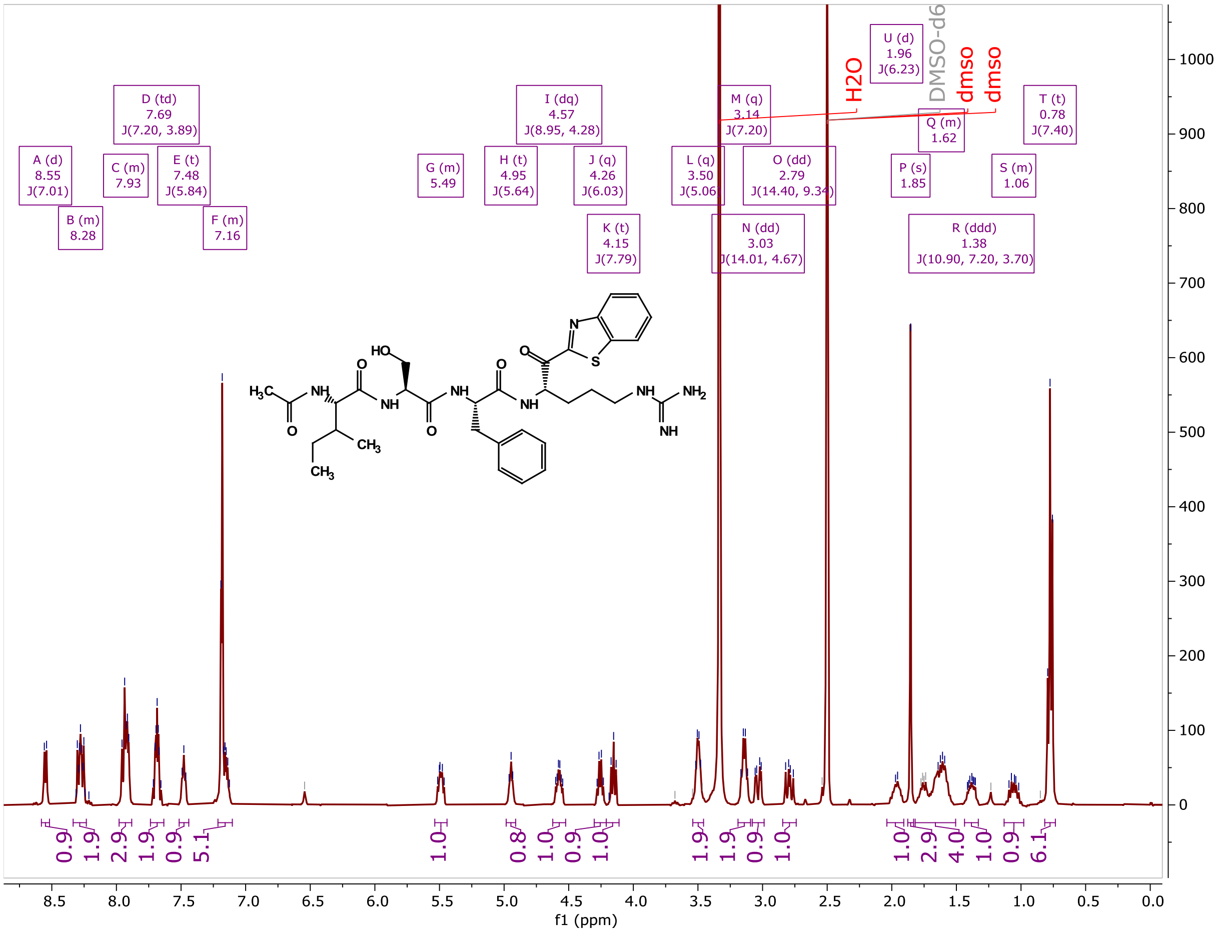

MM4038 (11)

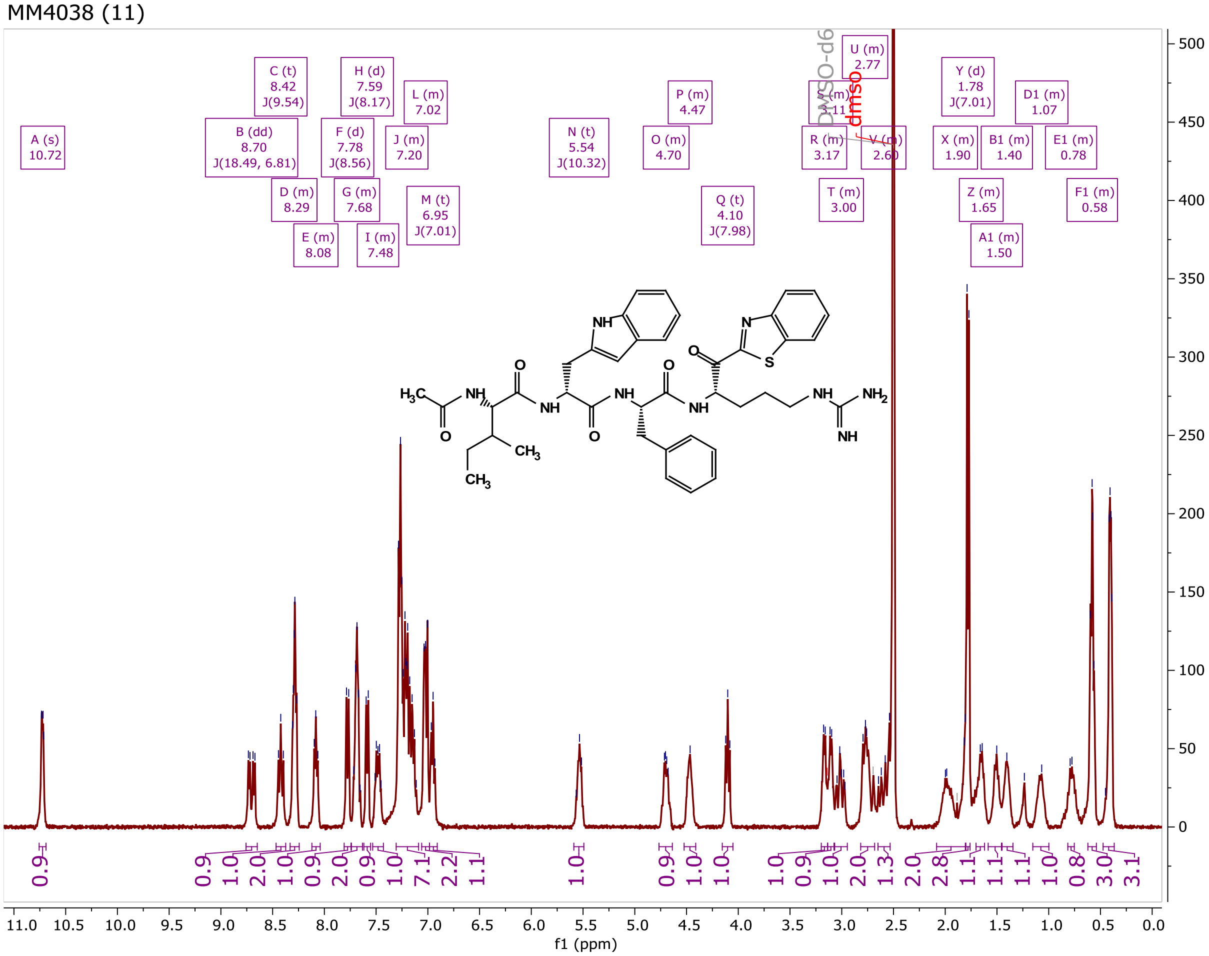


MM3187 (12)

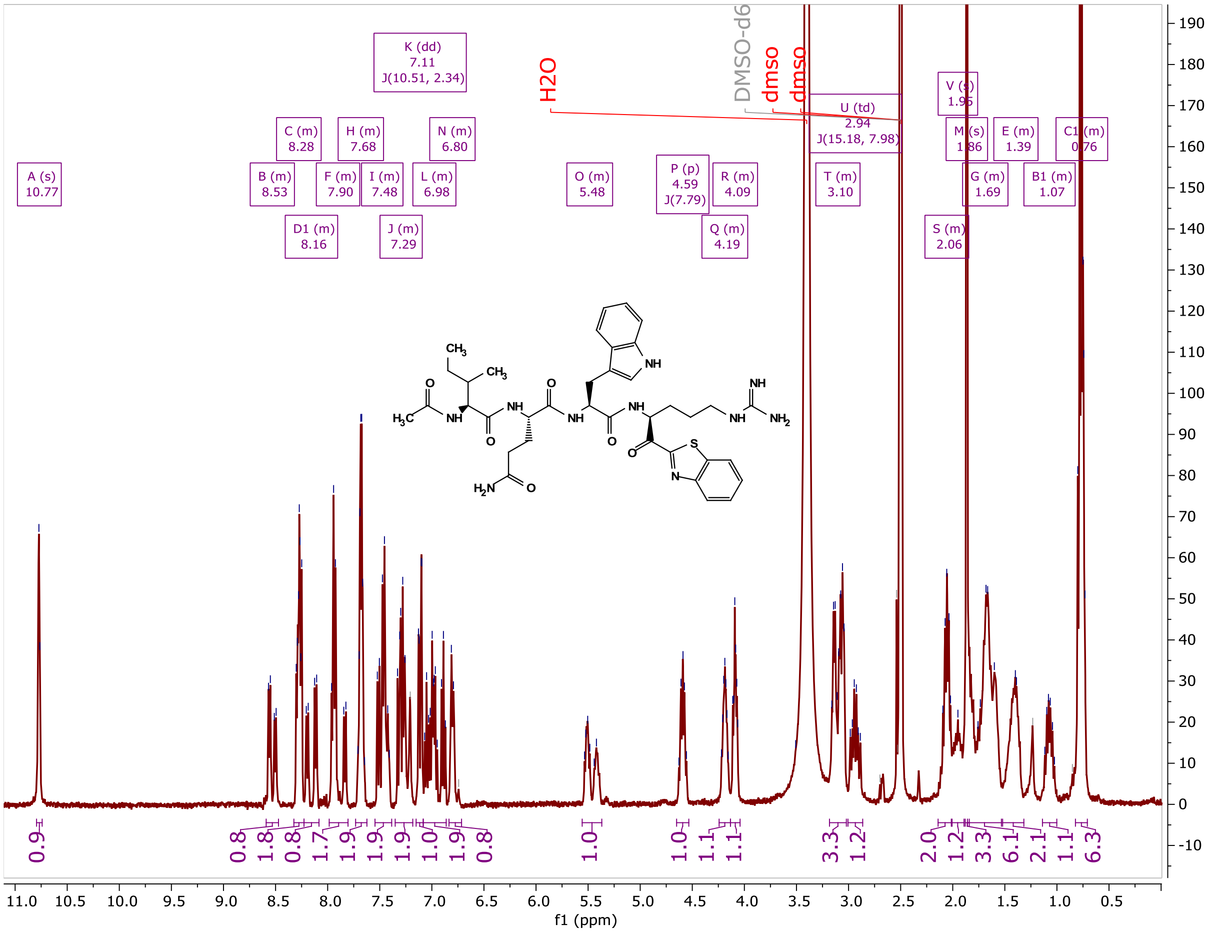

MM3186 (13)

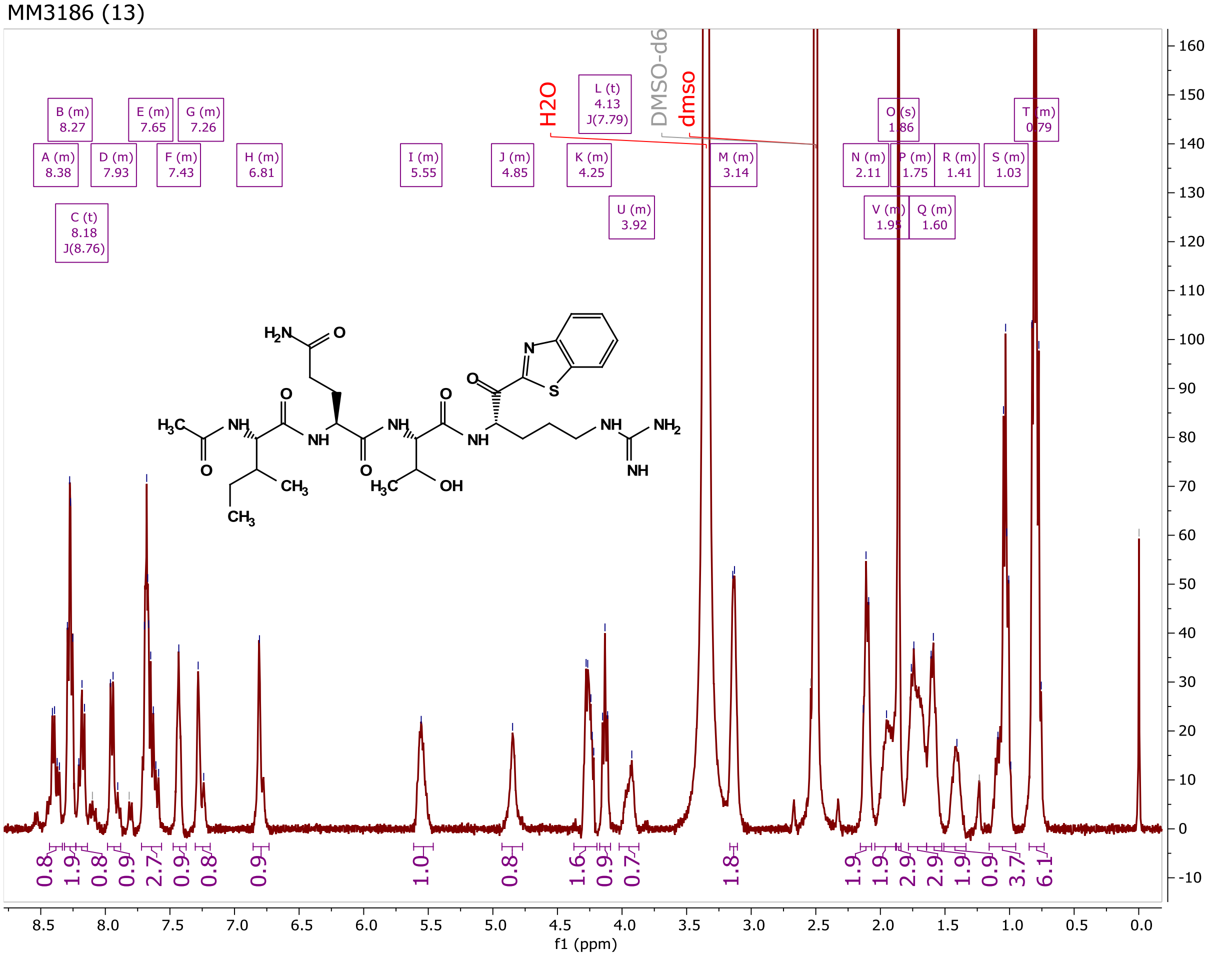


MM3180 (14)

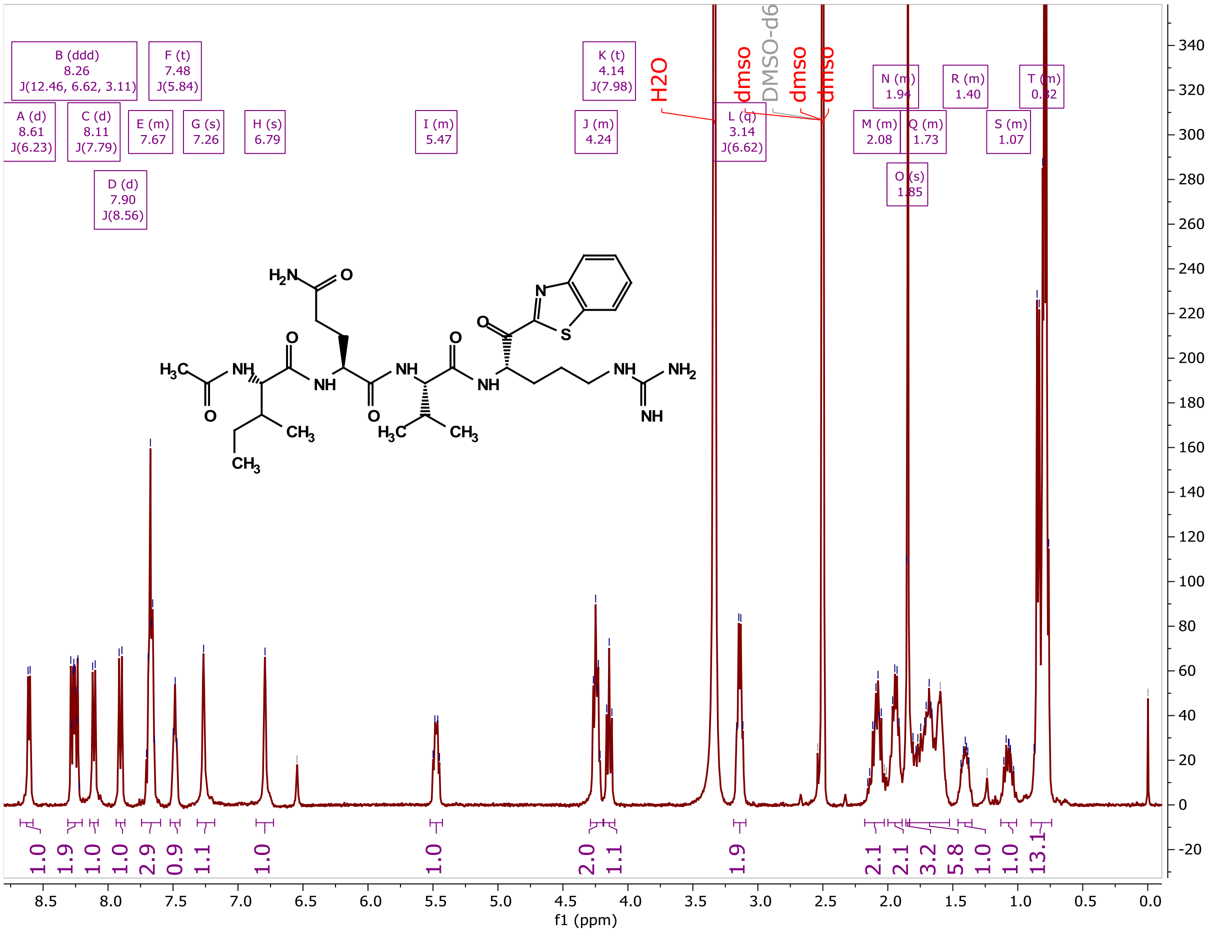

MM3178 (15)

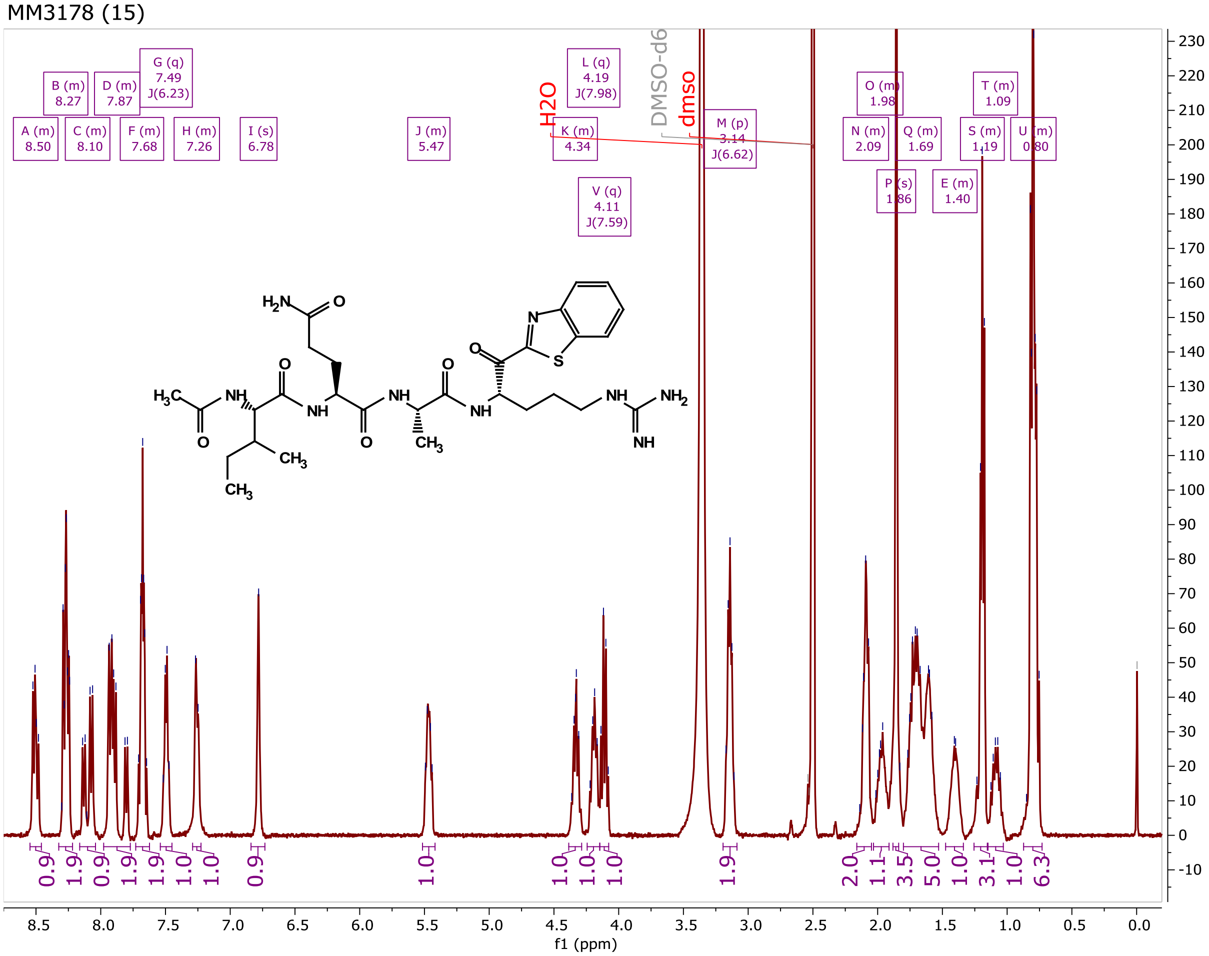


MM3177 (16)

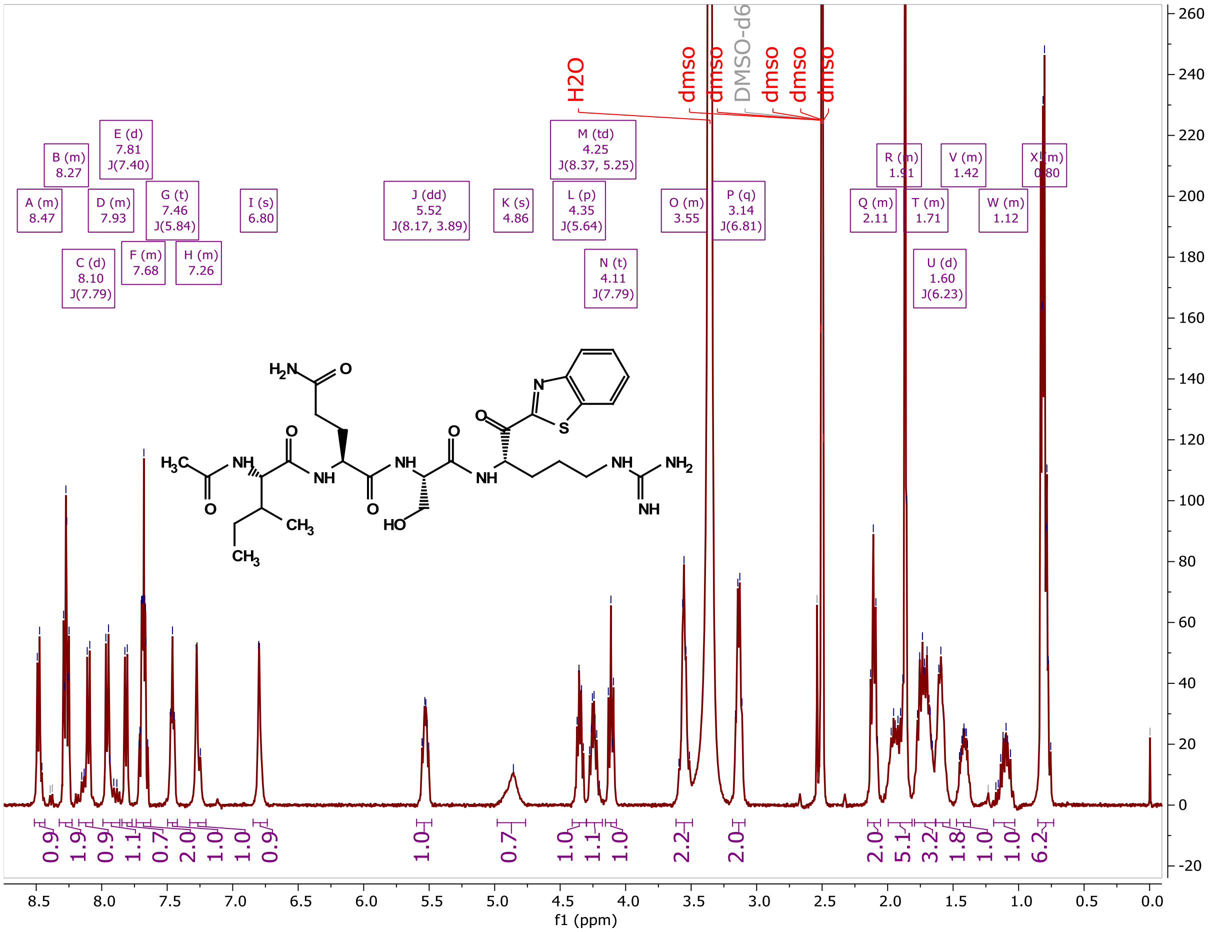

MM3194 (17)

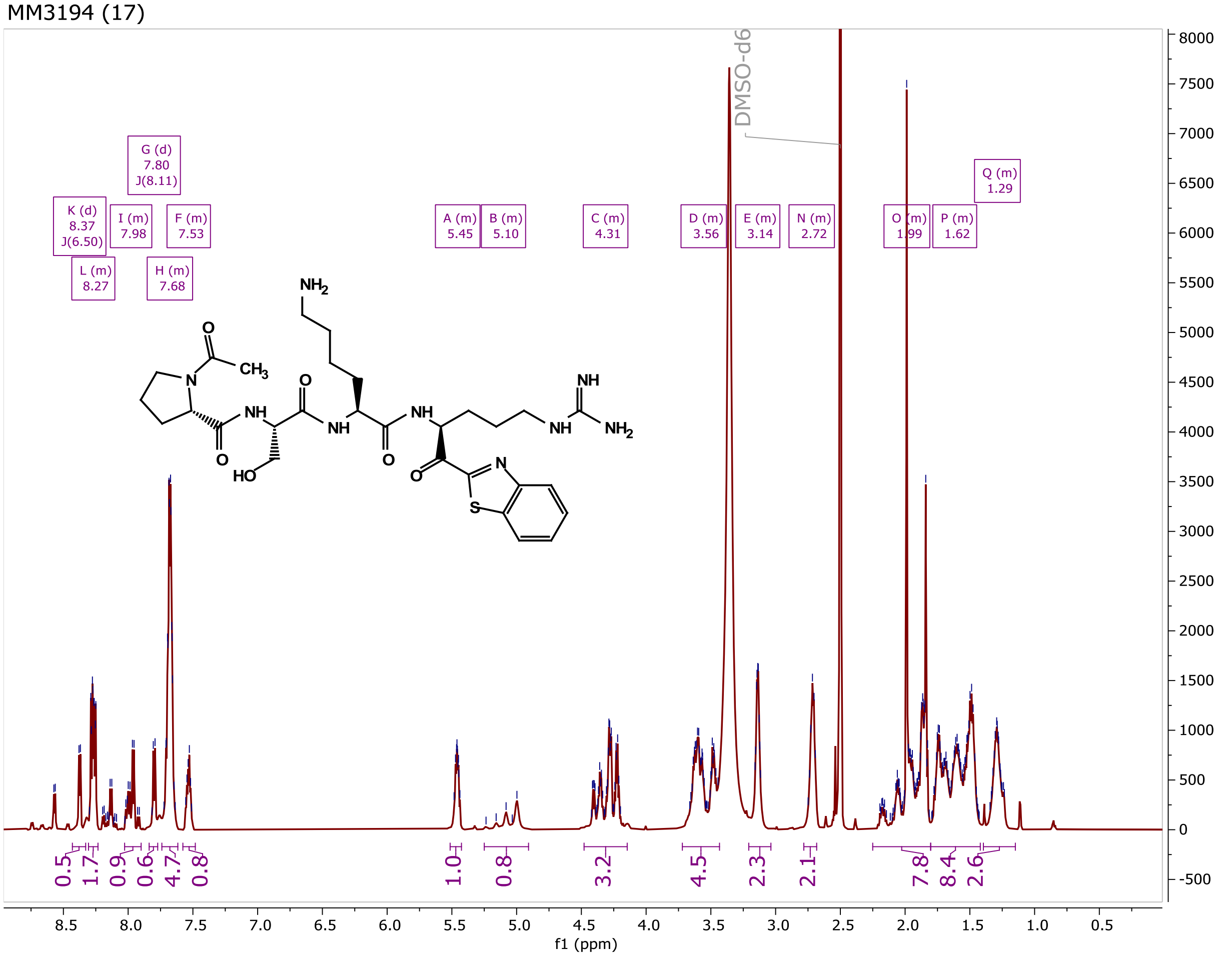


MM3144 (18)

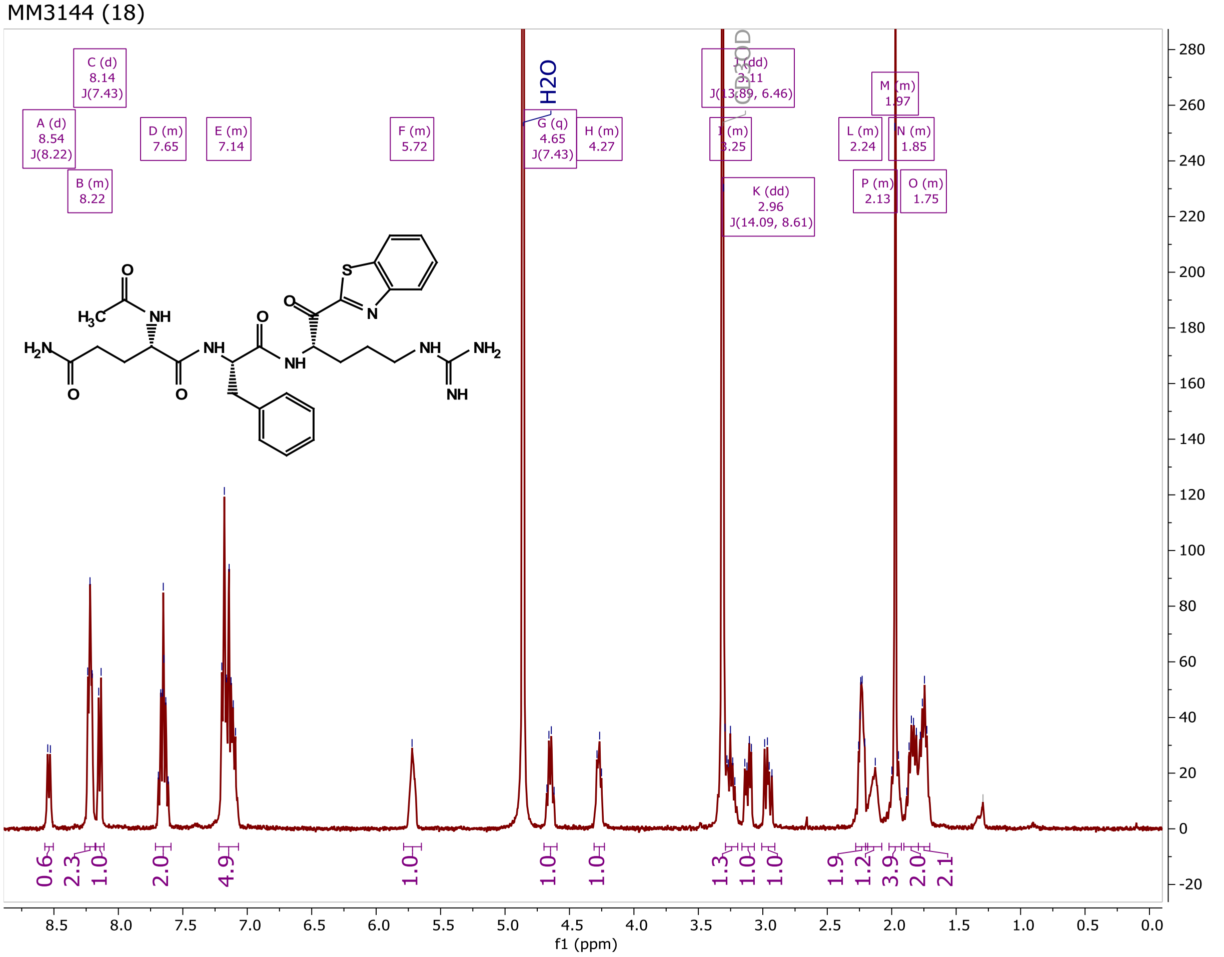

CA1041 (19)

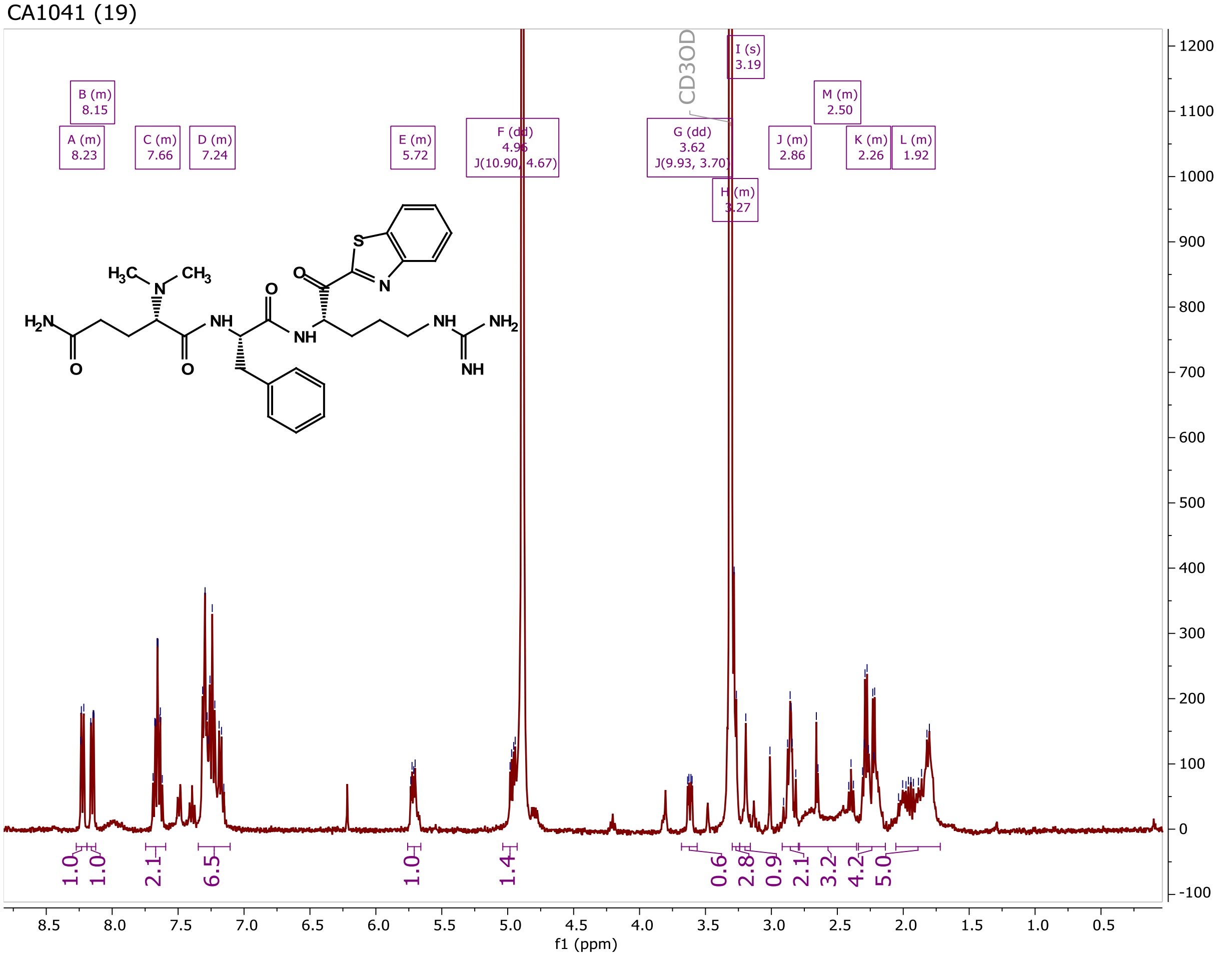


CA1033 (20)

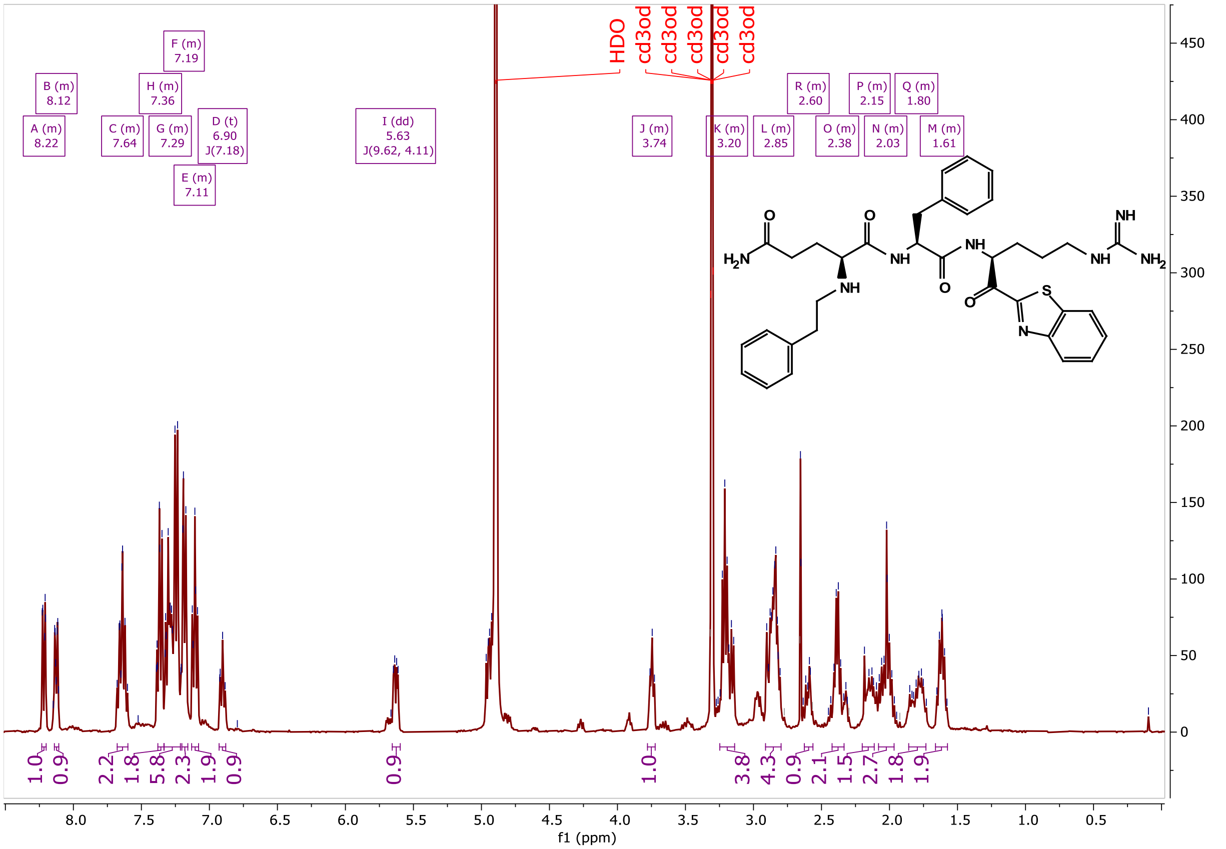

CA1022 (21)

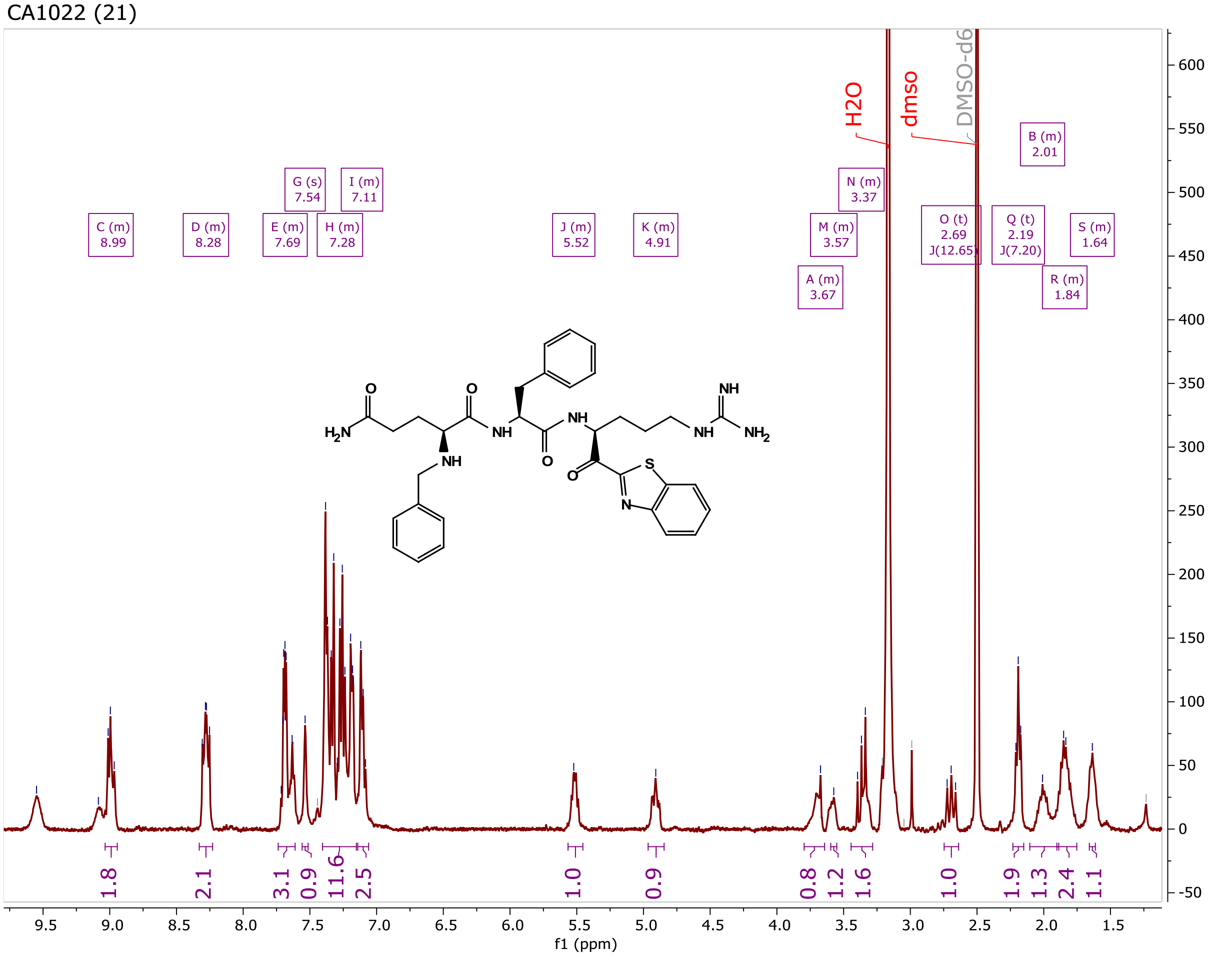


CA1018 (22)

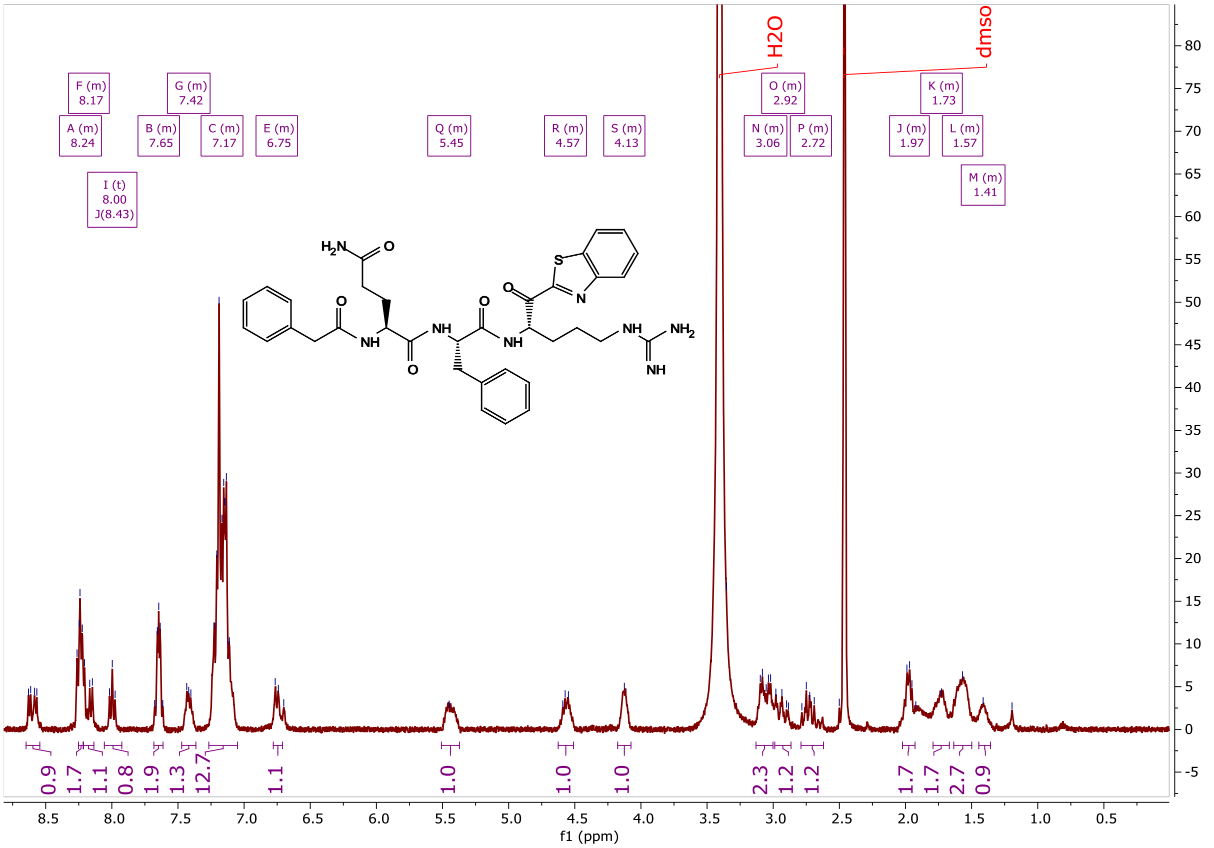

CA1043 (23)

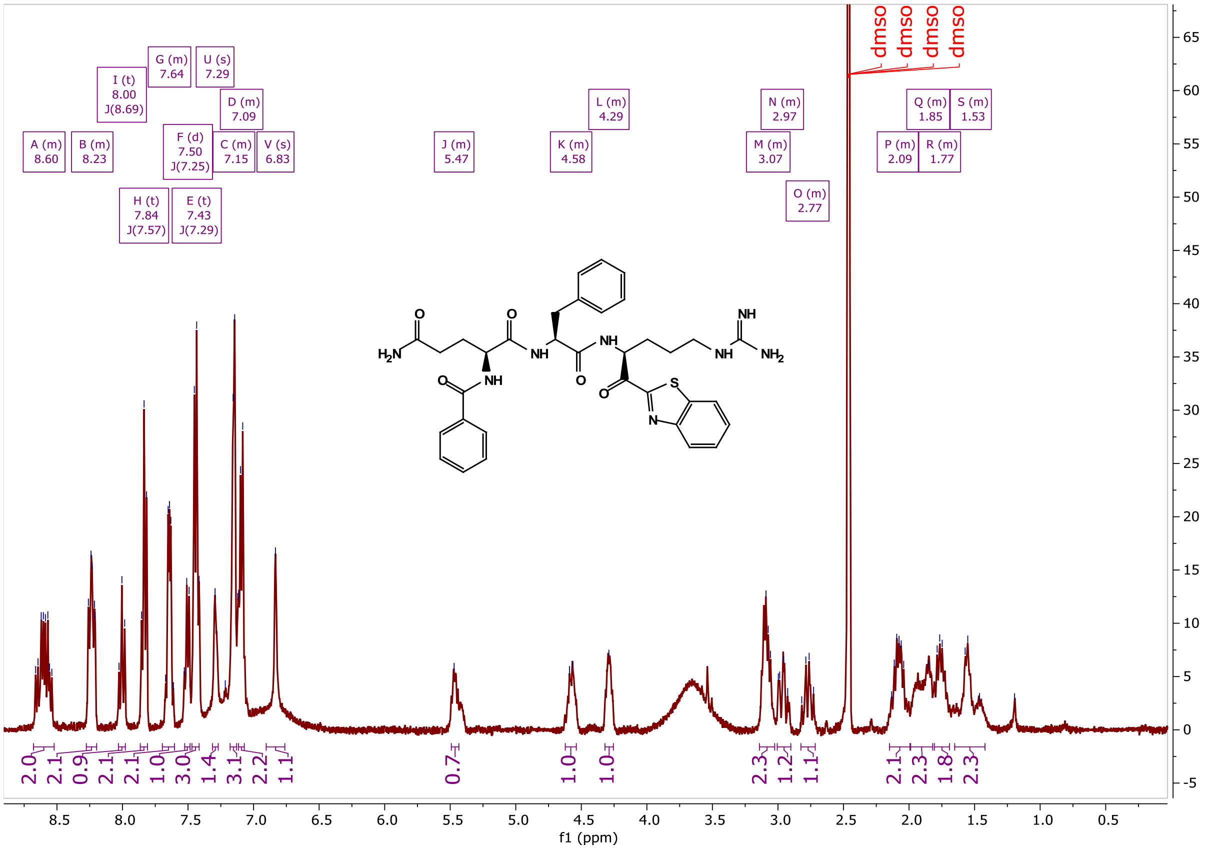


CA1046 (24)

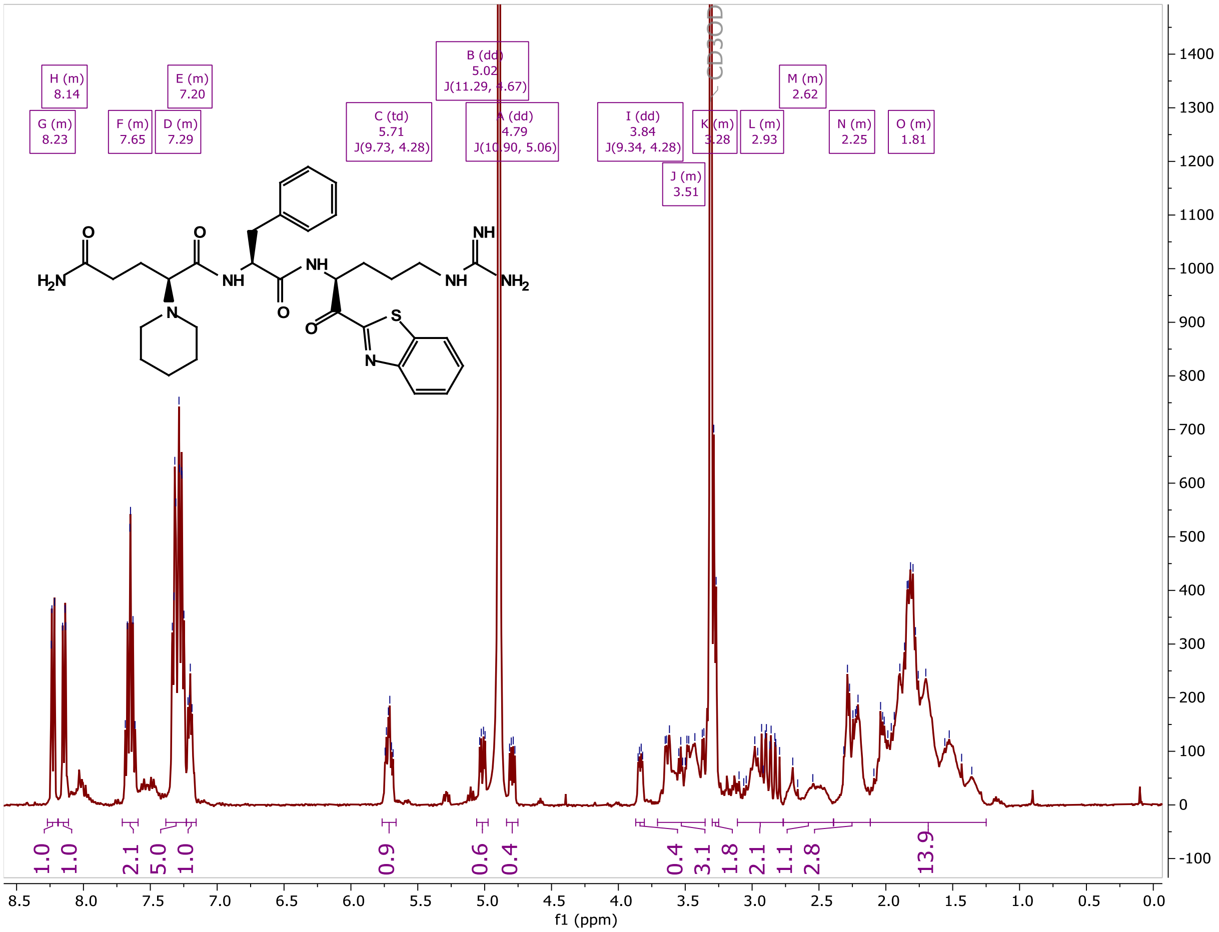

ZFH9141 (25)

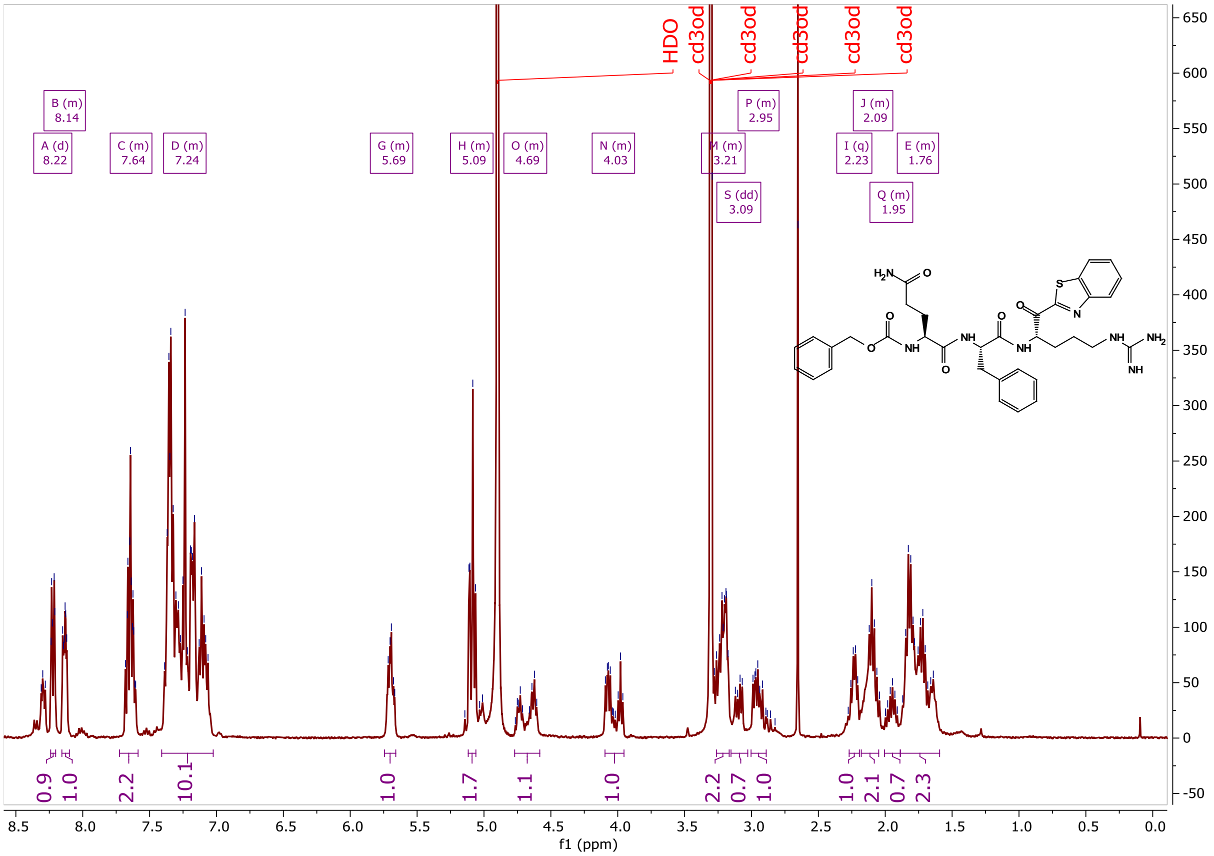

MF1101 (27)

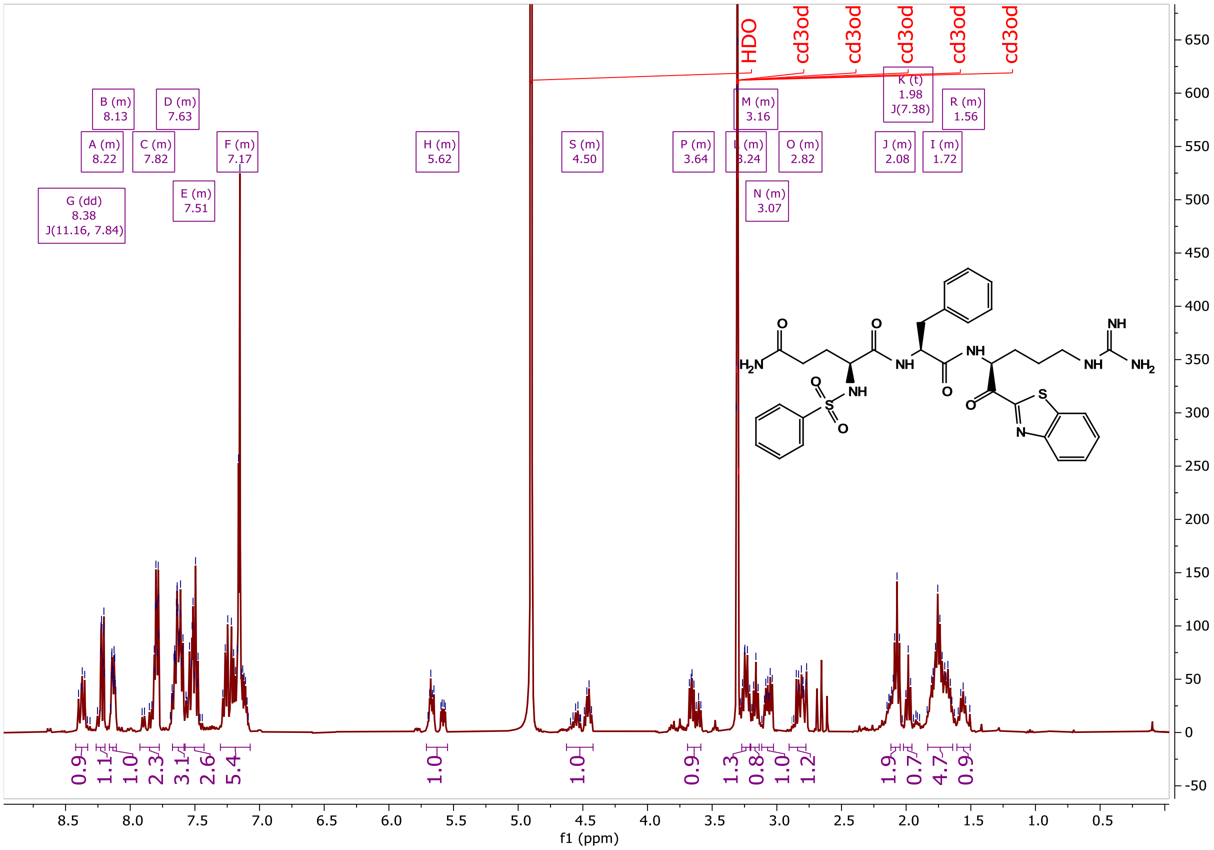

MF1105 (28)

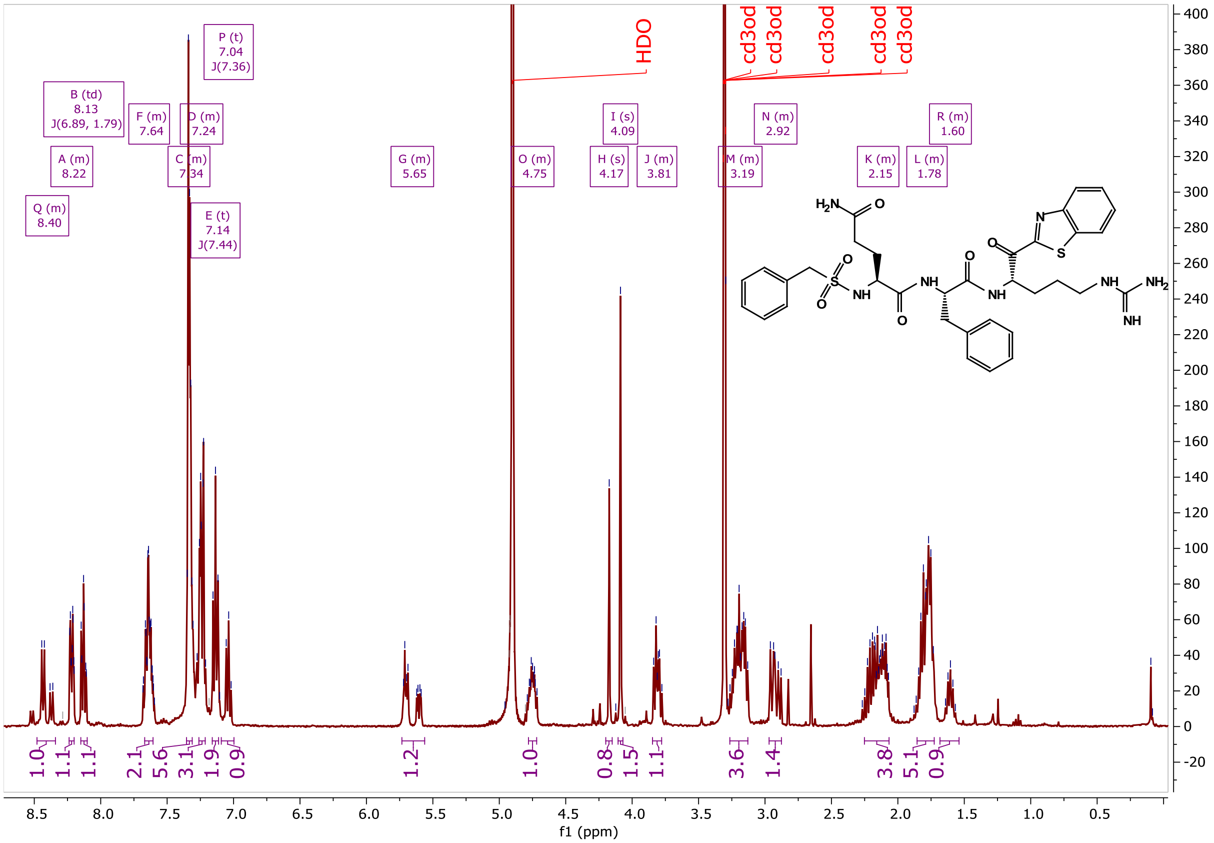


MF1104 (29)

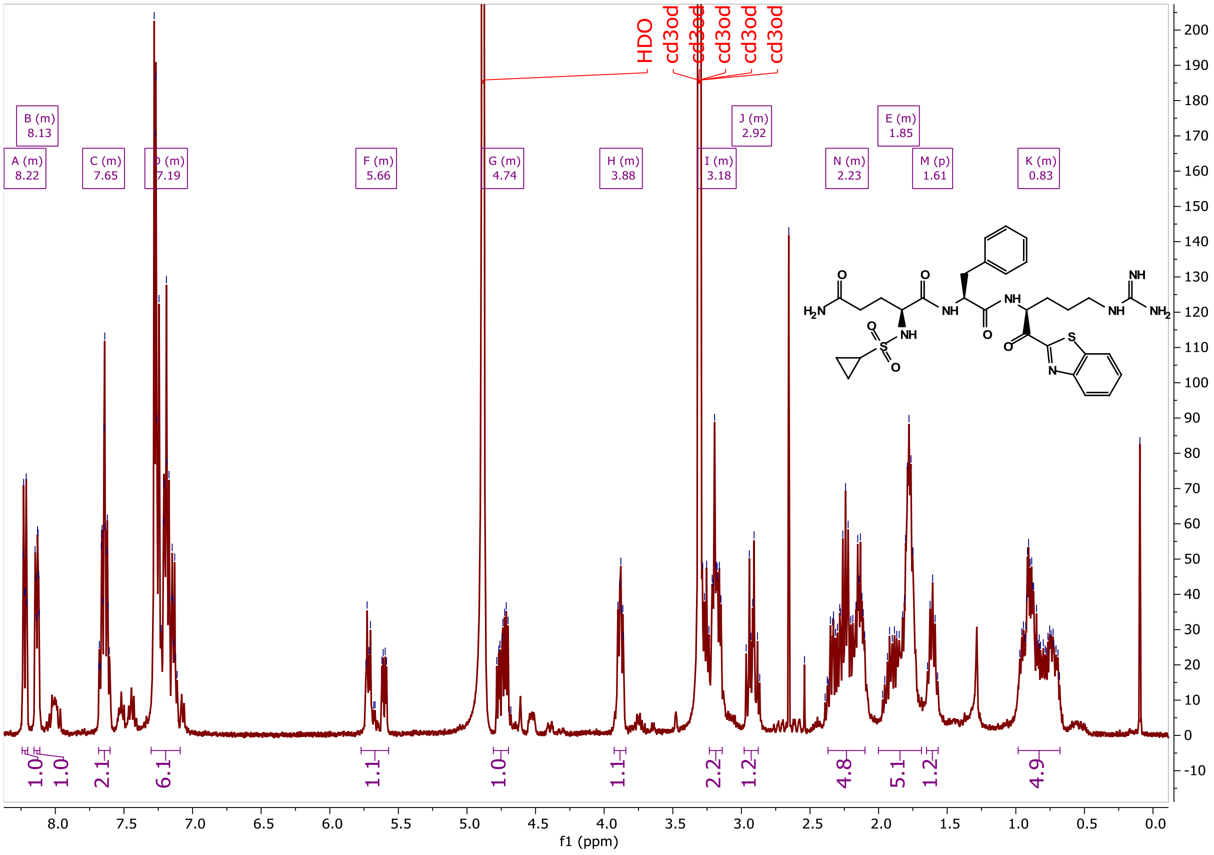

### LCMS Data:

**MM3116 (1)**

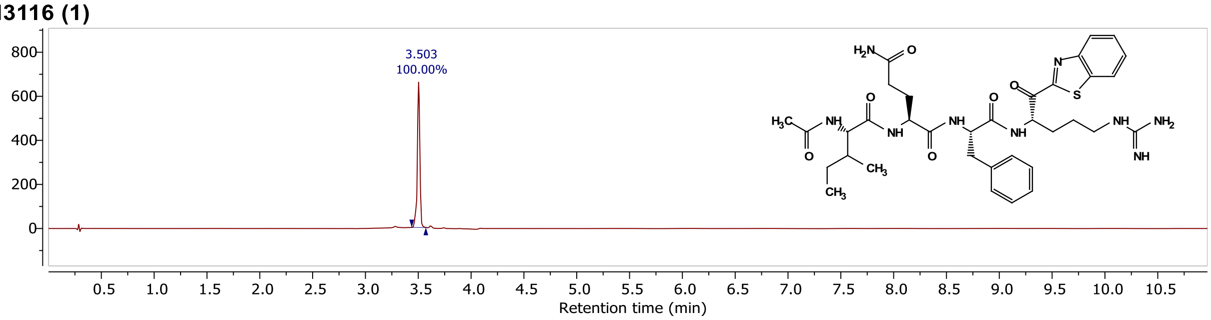


\\quatro-micro...2020\MM3116F2.D Injection 1 Function 1 (MM3116F2) TIC

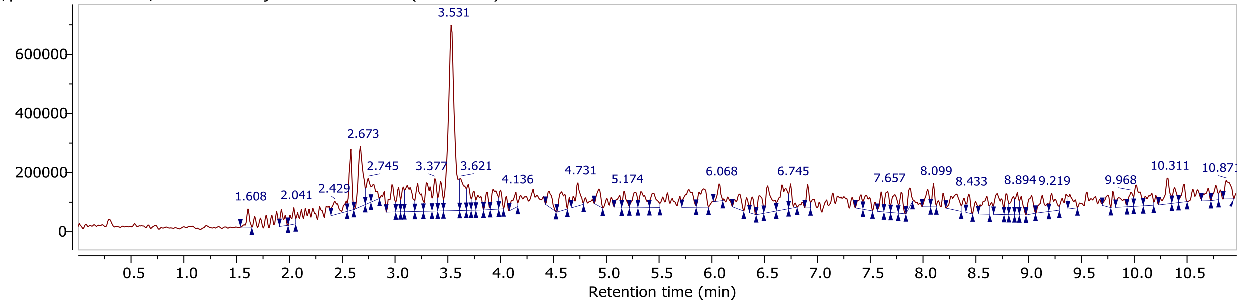


\\quatro-micro...2020\MM3116F2.D Injection 1 Function 1 (MM3116F2) MS + spectrum 3.53

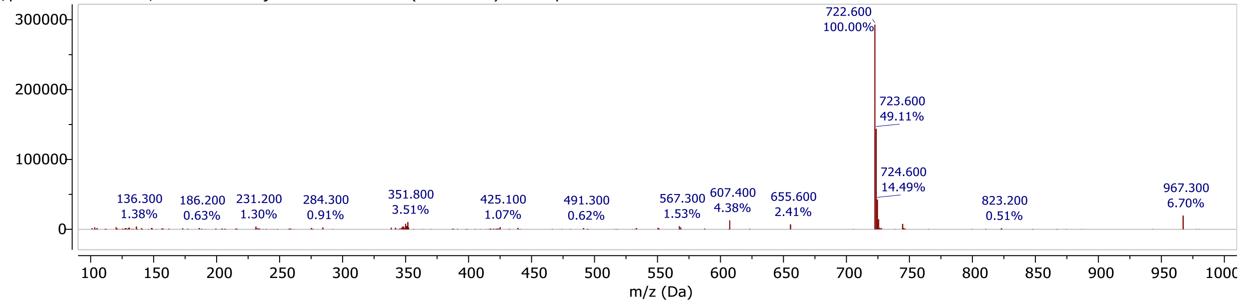

## MM3122 (2)

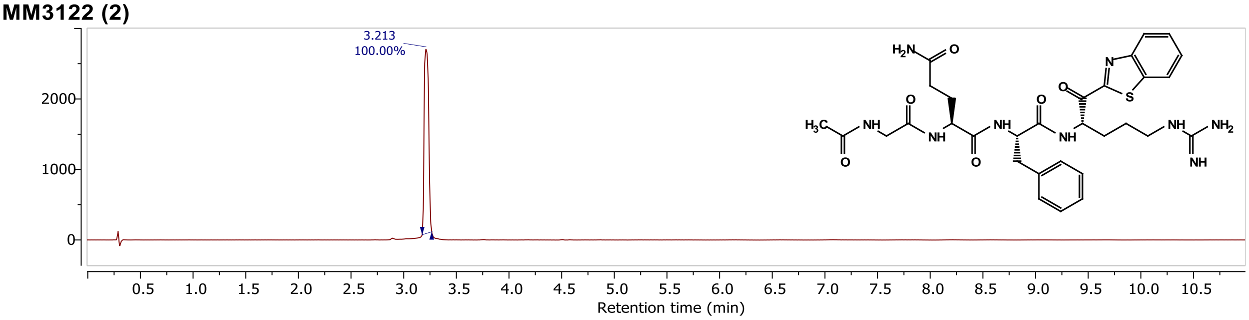


D:\Matt's comp...LCMS\MM3122S.D\ Injection 1 Function 1 (MM3122S) TIC

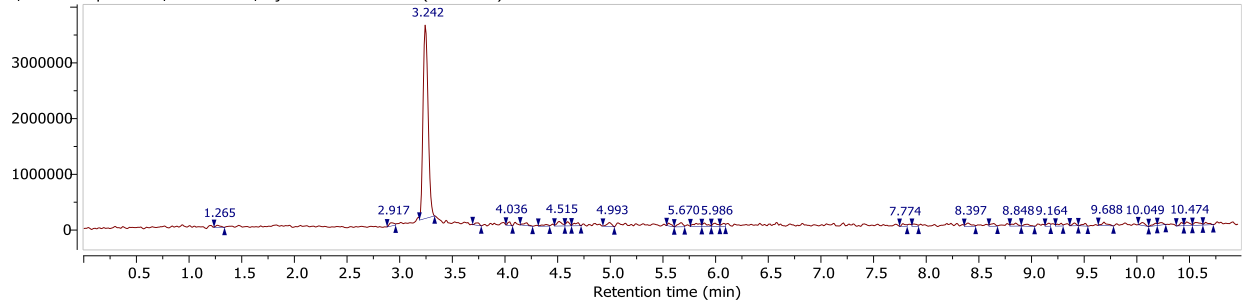


D:\Matt's comp...LCMS\MM3122S.D\ Injection 1 Function 1 (MM3122S) MS + spectrum 3.24

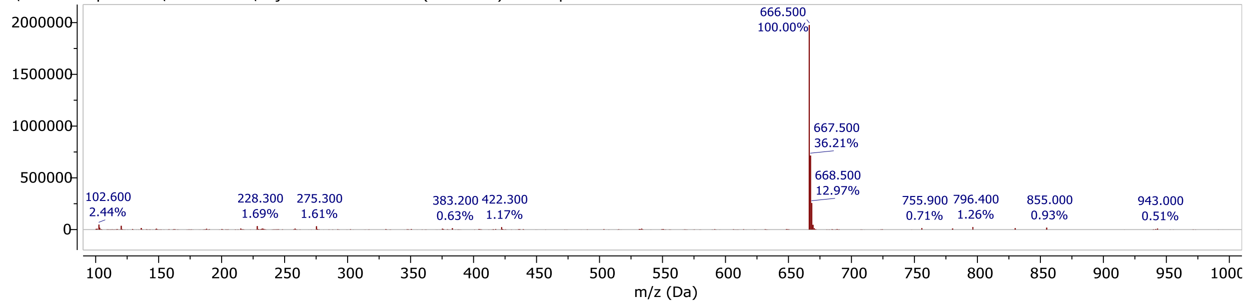


## MM3123 (3)

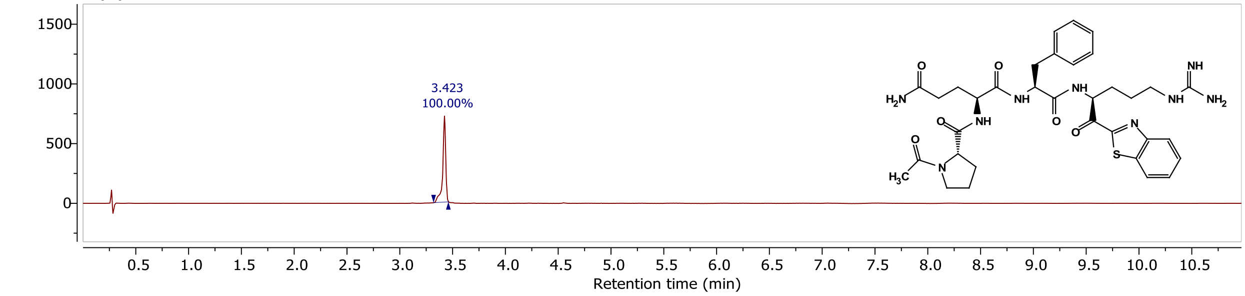


D:\Matt's comp...LCMS\MM3123S.D\ Injection 1 Function 1 (MM3123S) TIC

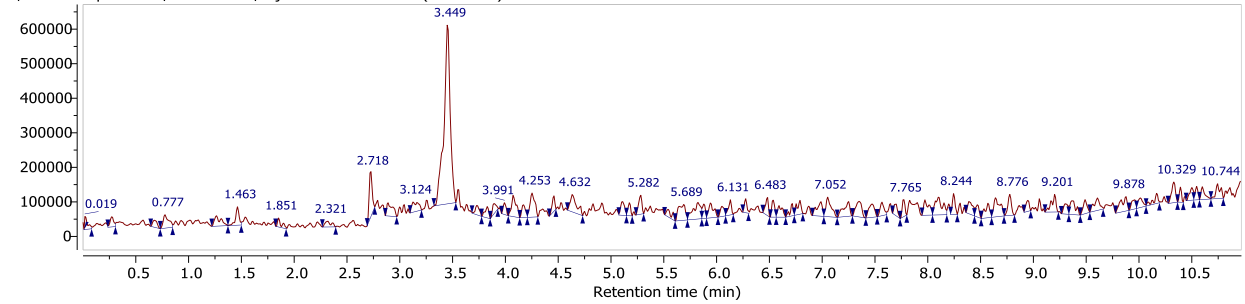


D:\Matt's comp...LCMS\MM3123S.D\ Injection 1 Function 1 (MM3123S) MS + spectrum 3.32..3.49

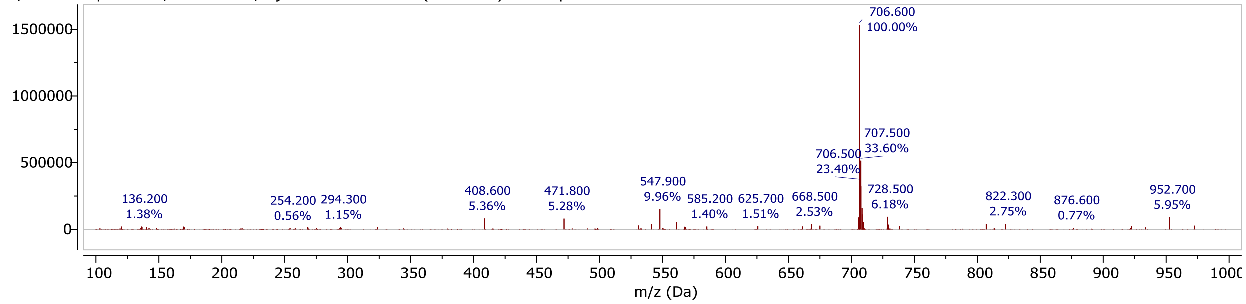

## MM3131 (4)

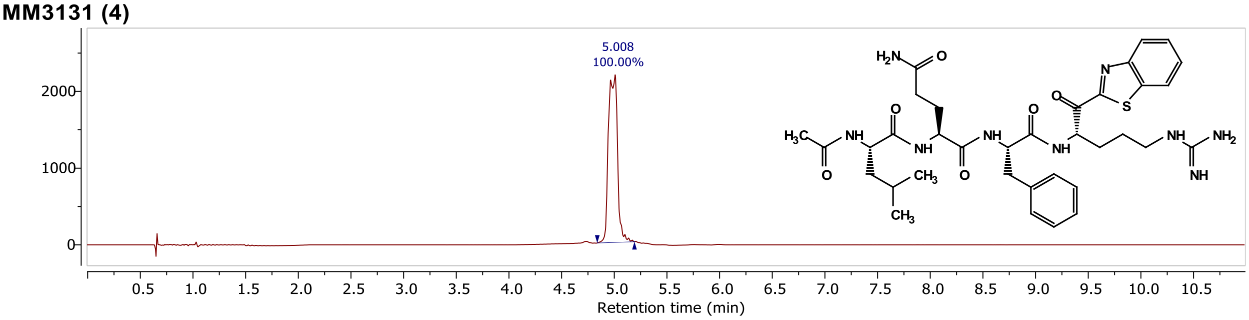


\\quatro-micro...AY2024\MM3131.D Injection 1 Function 1 (MM3131) TIC

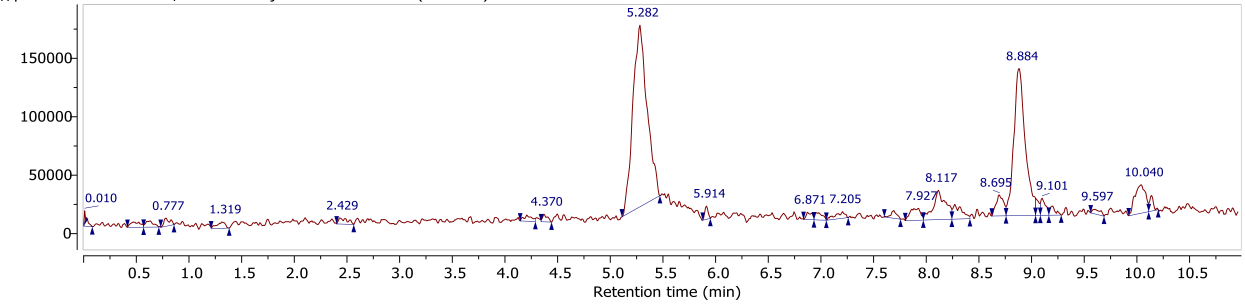


\\quatro-micro...AY2024\MM3131.D Injection 1 Function 1 (MM3131) MS + spectrum 5.07..5.48

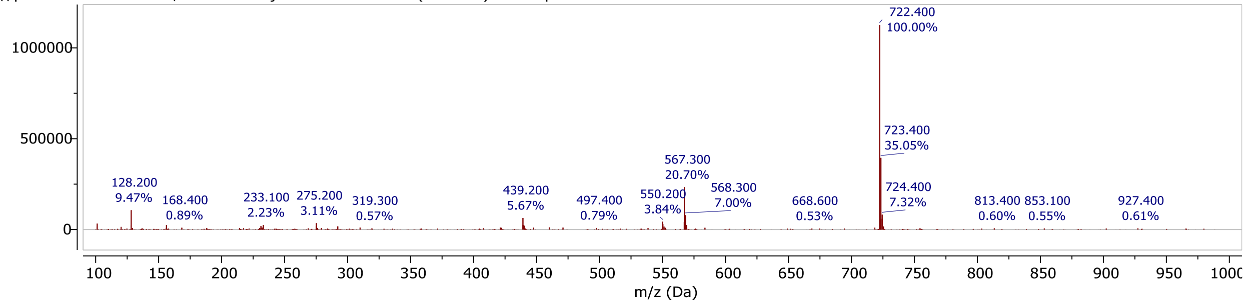


## MM3130 (5)

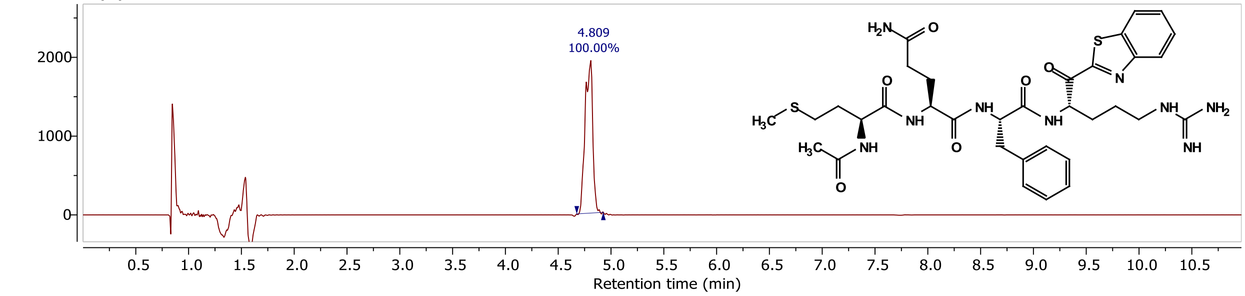


\\quatro-micro...AY2024\MM3130.D Injection 1 Function 1 (MM3130) TIC

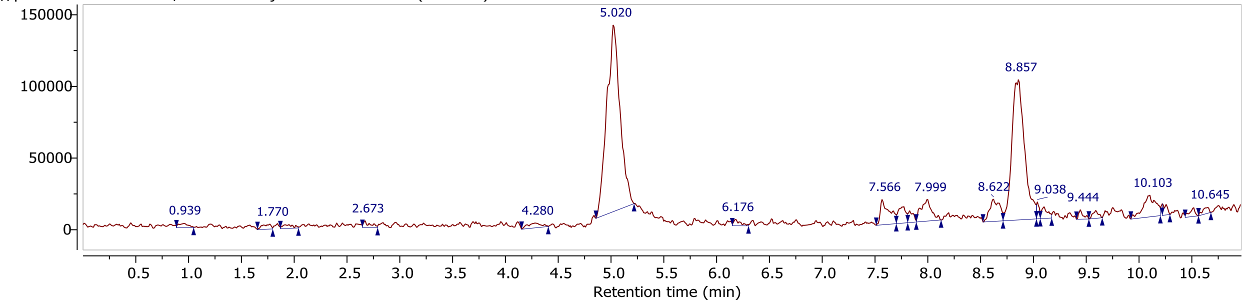


\\quatro-micro...AY2024\MM3130.D Injection 1 Function 1 (MM3130) MS + spectrum 4.85..5.25

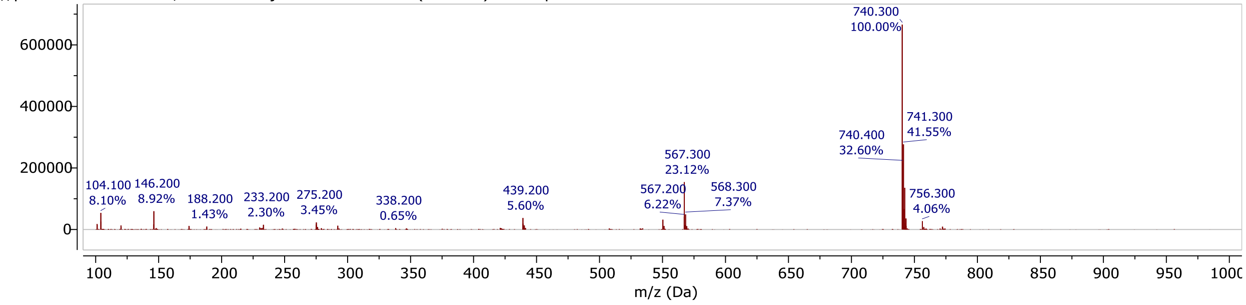

## MM4037 (7)

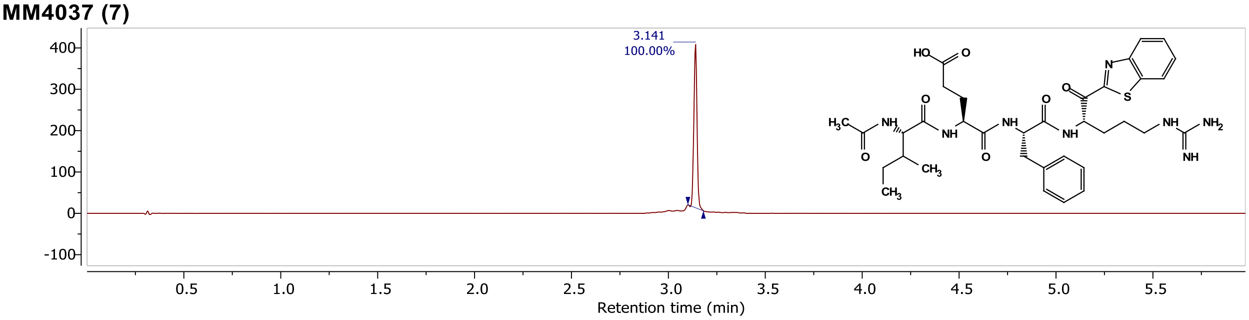


D:/Matt's comp...LCMS/MM4037-2.D Injection 1 Function 1 (MM4028-2) TIC

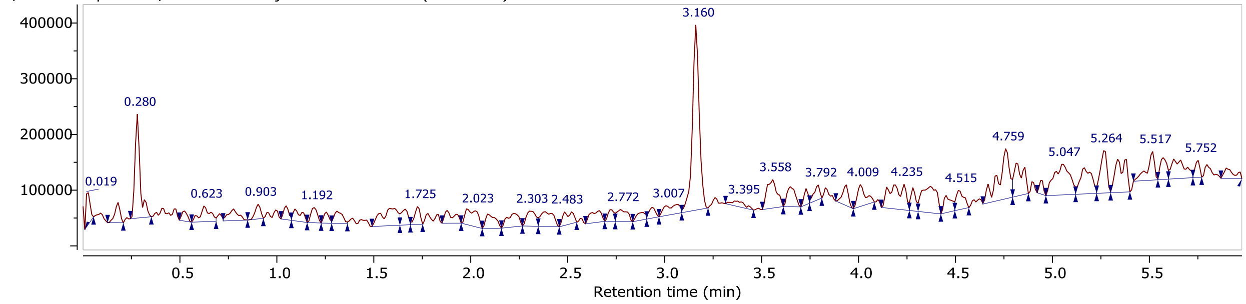


D:/Matt's comp...LCMS/MM4037-2.D Injection 1 Function 1 (MM4028-2) MS + spectrum 3.14

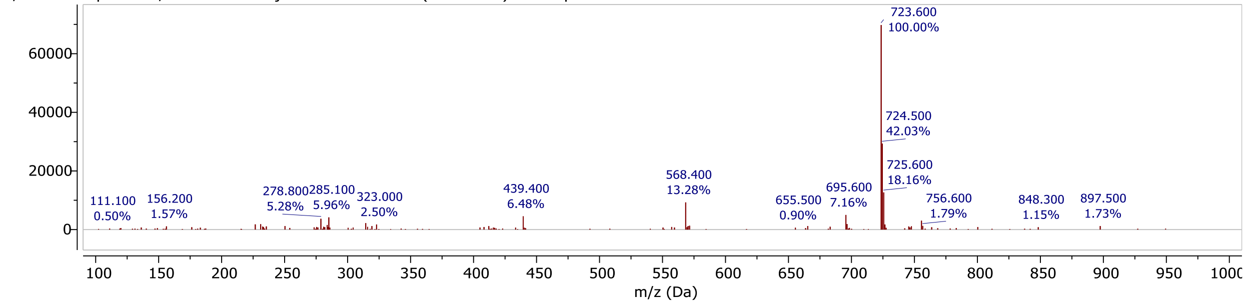


## MM4028 (8)

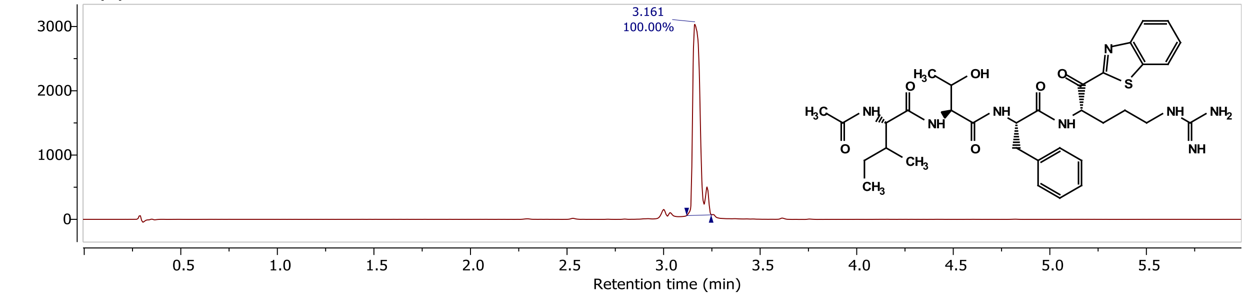


D:/Matt's comp...LCMS/MM4028-2.D Injection 1 Function 1 (MM4037-2) TIC

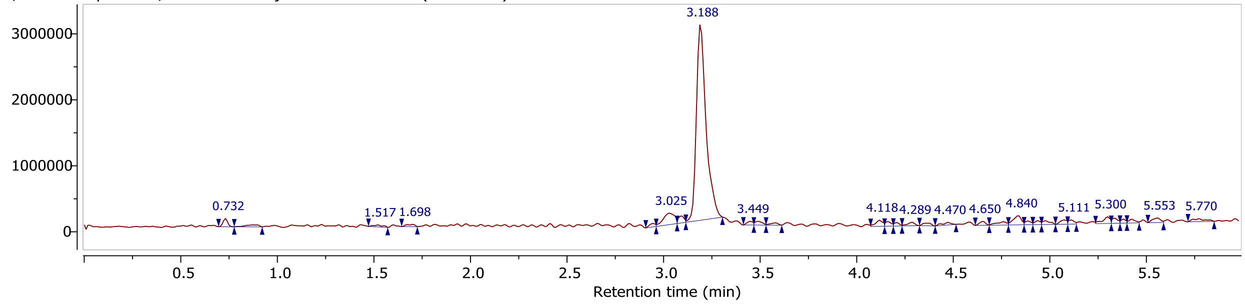


D:/Matt's comp...LCMS/MM4028-2.D Injection 1 Function 1 (MM4037-2) MS + spectrum 3.13..3.30

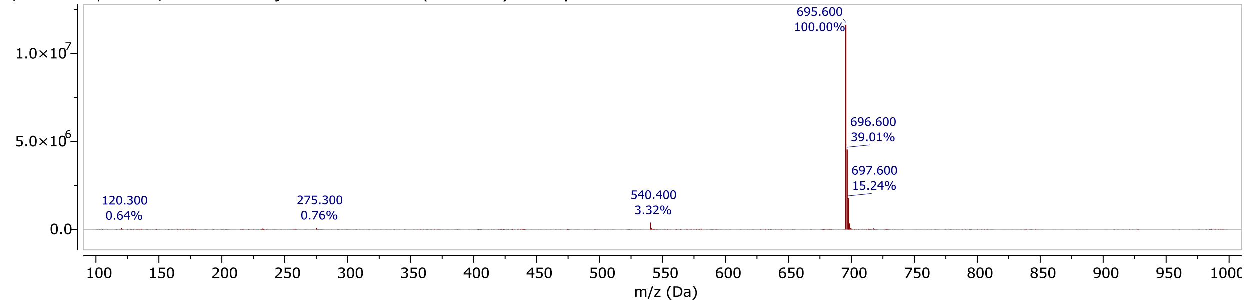

## MM4027 (9)

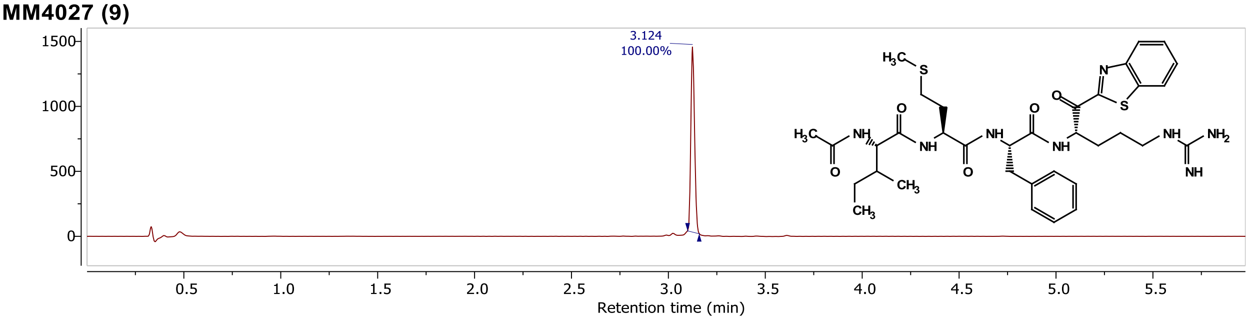


D:/Matt's comp...LCMS/MM4027-1.D Injection 1 Function 1 (MM4027-1F) TIC

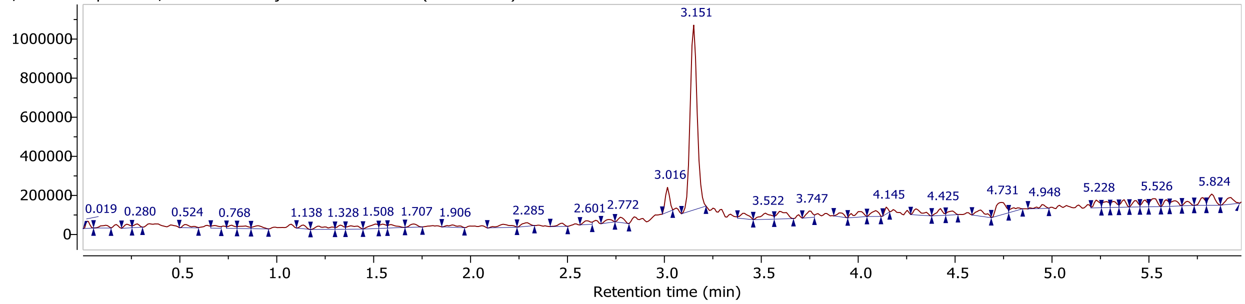


D:/Matt's comp...LCMS/MM4027-1.D Injection 1 Function 1 (MM4027-1F) MS + spectrum 3.12

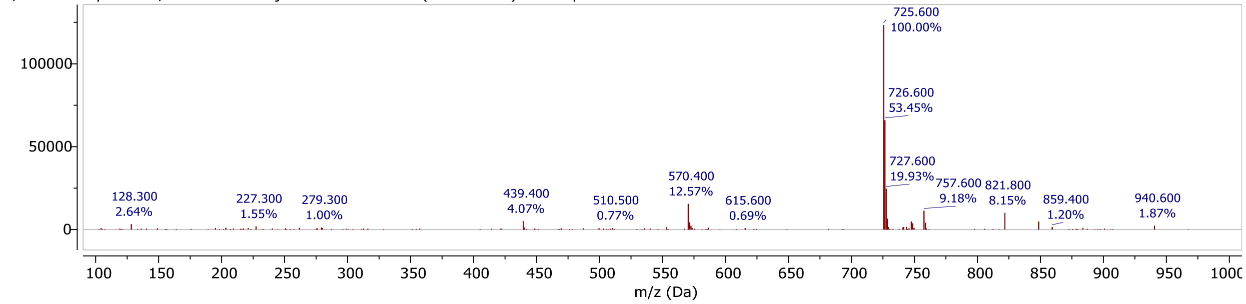


## MM4009 (10)

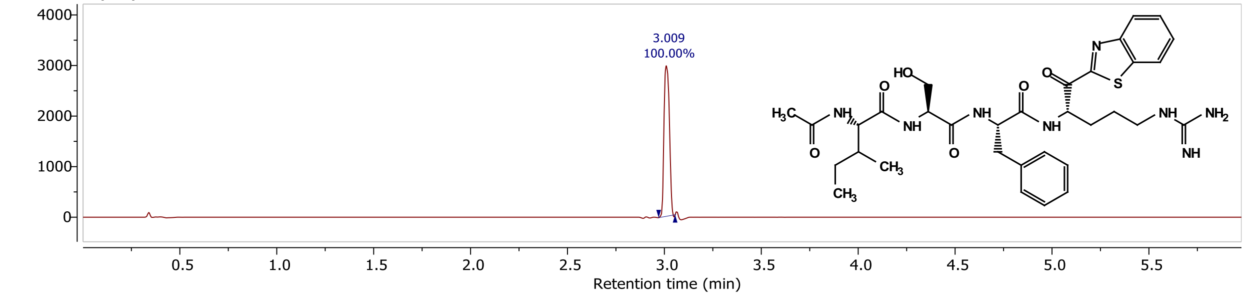


D:/Matt's comp...LCMS/MM4009-2.D Injection 1 Function 1 (MM4009-2F) TIC

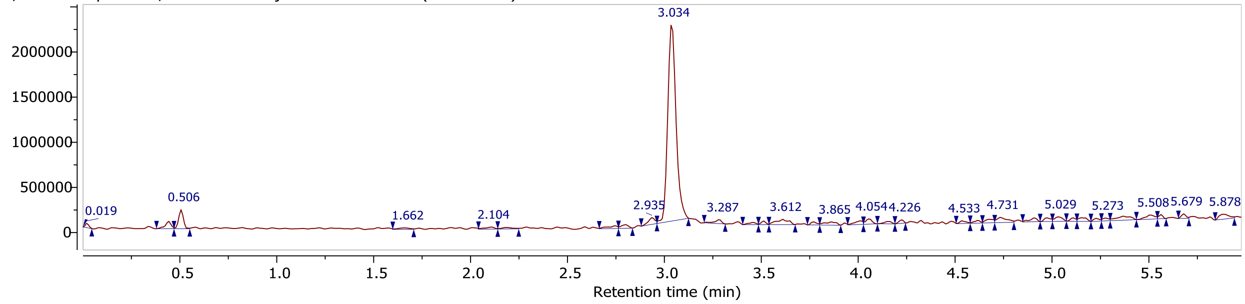


D:/Matt's comp...LCMS/MM4009-2.D Injection 1 Function 1 (MM4009-2F) MS + spectrum 2.98..3.11

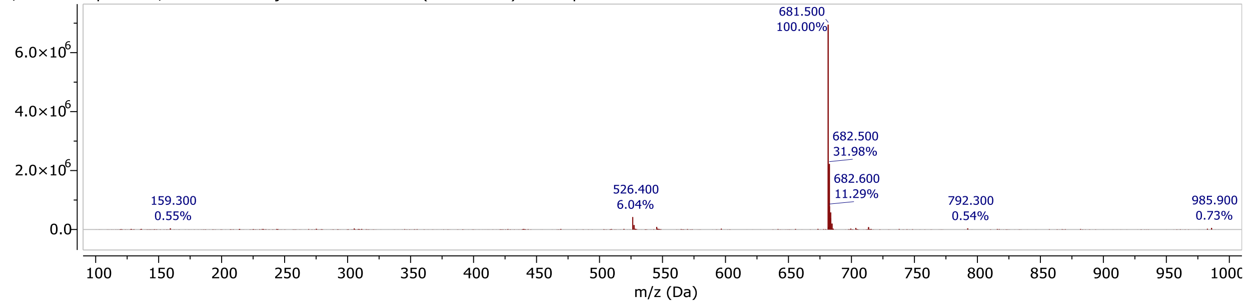

**MM4038 (11)**

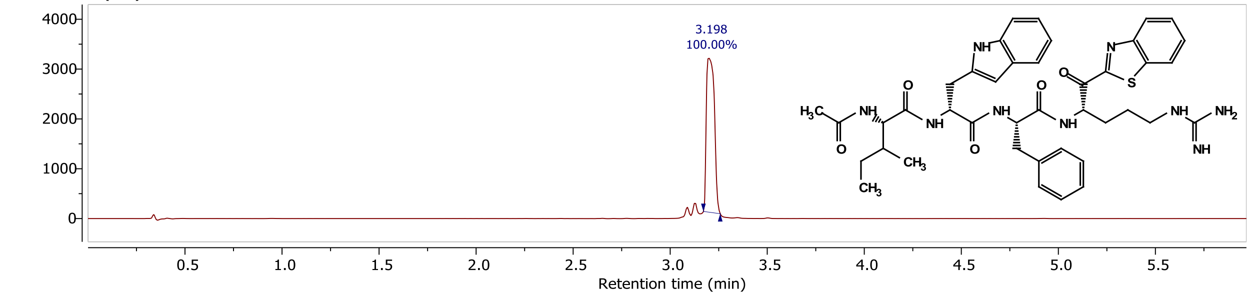


D:/Matt's comp... LCMS/MM4038F.D Injection 1 Function 1 (MM4038F) TIC

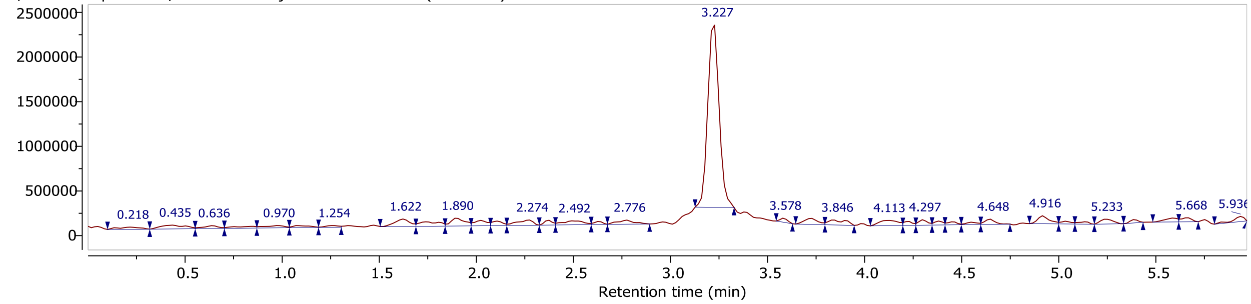


D:/Matt's comp... LCMS/MM4038F.D Injection 1 Function 1 (MM4038F) MS + spectrum 3.14..3.31

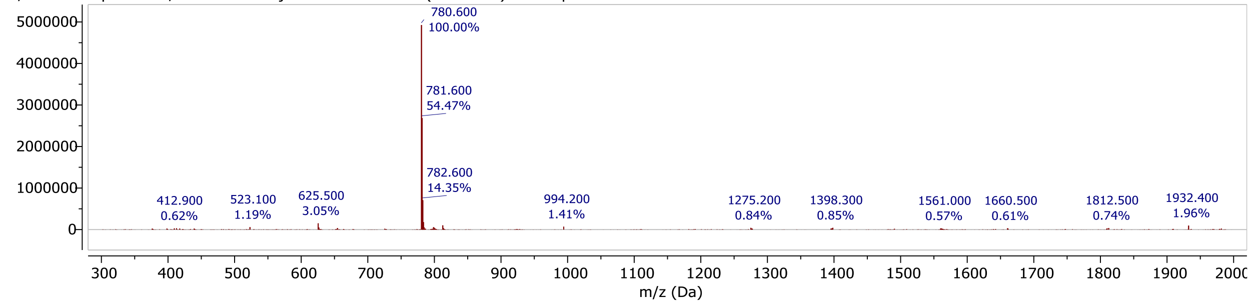


**MM3187 (12)**

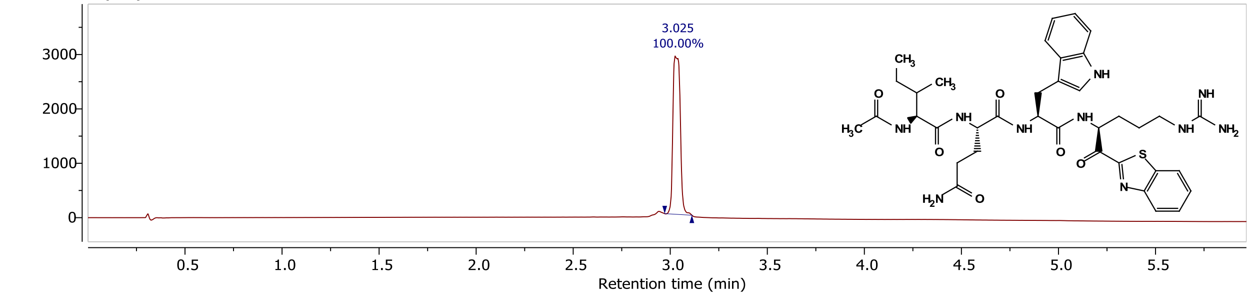


D:/Matt's comp...s LCMS/MM3187.D Injection 1 Function 1 (MM3187) TIC

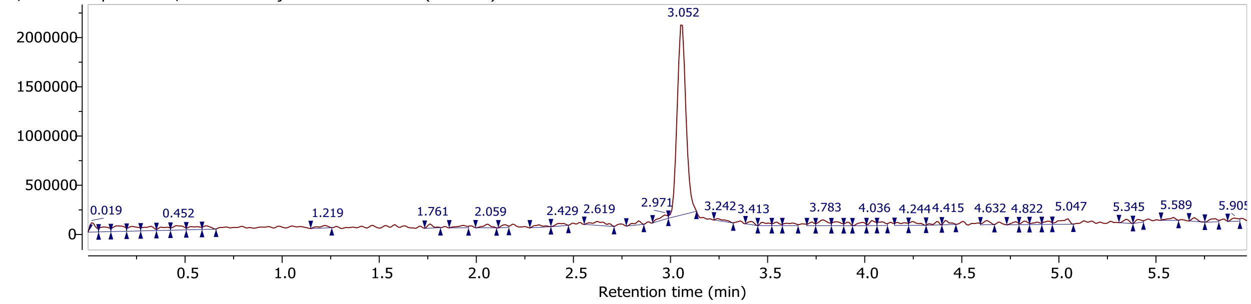


D:/Matt's comp...s LCMS/MM3187.D Injection 1 Function 1 (MM3187) MS + spectrum 3.00..3.14

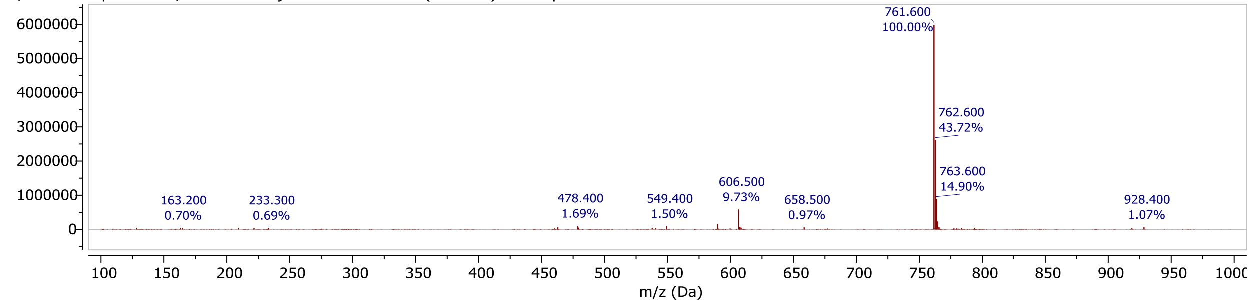

## MM3186 (13)

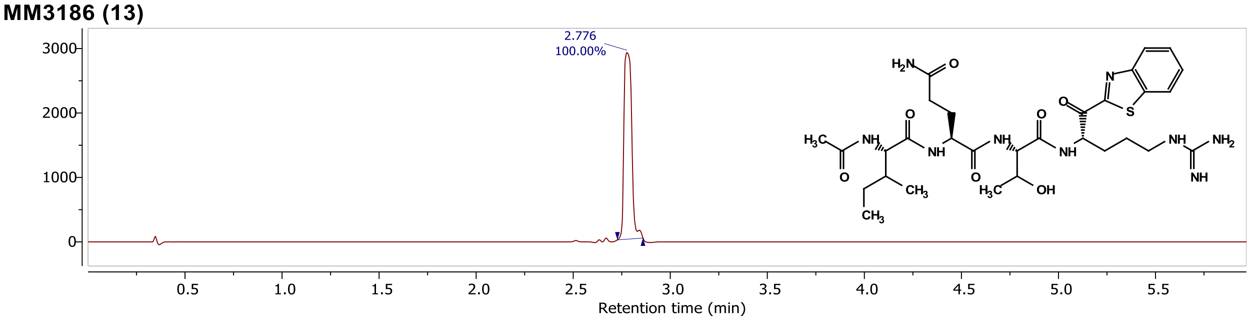


D:/Matt's comp...s LCMS/MM3186.D Injection 1 Function 1 (MM3186) TIC

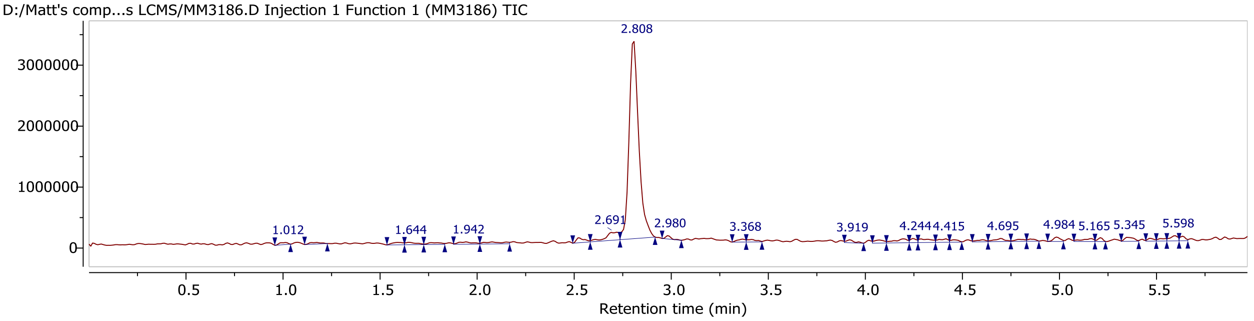


D:/Matt's comp...s LCMS/MM3186.D Injection 1 Function 1 (MM3186) MS + spectrum 2.74..2.91

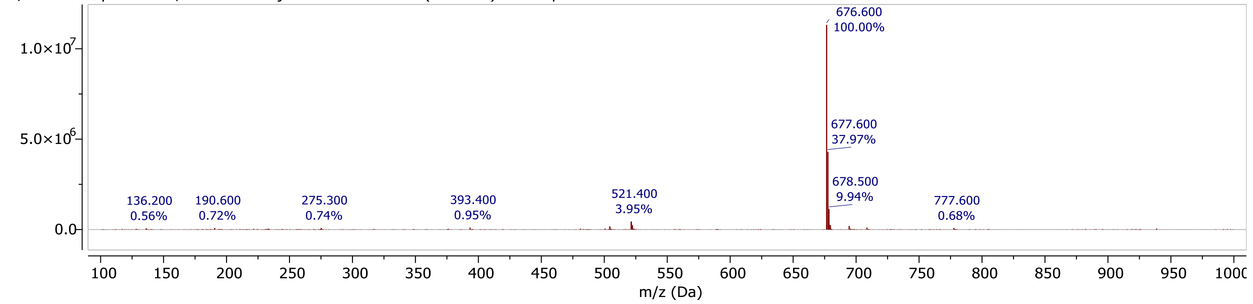


## MM3180 (14)

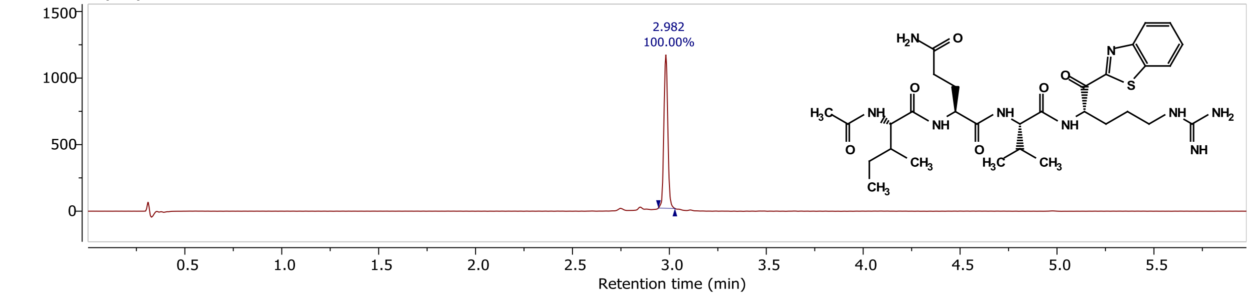


D:/Matt's comp...LCMS/MM3180-2.D Injection 1 Function 1 (MM3180-2) TIC

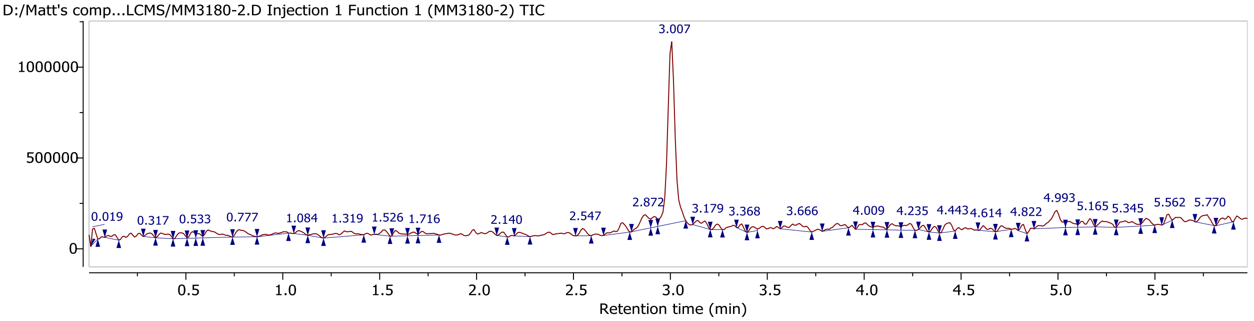


D:/Matt's comp...LCMS/MM3180-2.D Injection 1 Function 1 (MM3180-2) MS + spectrum 2.95..3.06

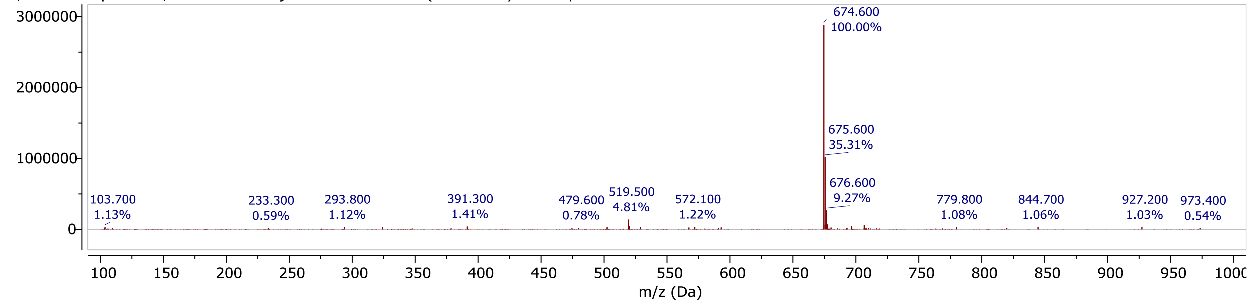

**MM3178 (15)**

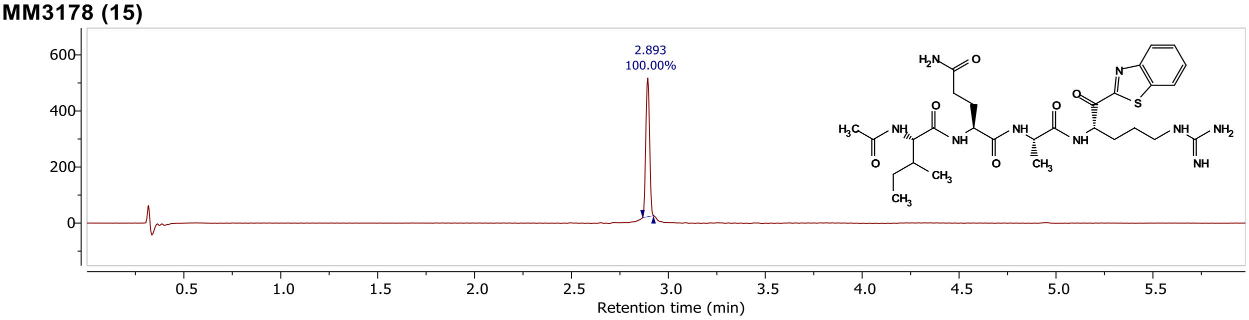


D:/Matt's comp...s LCMS/MM3178.D Injection 1 Function 1 (MM3178) TIC

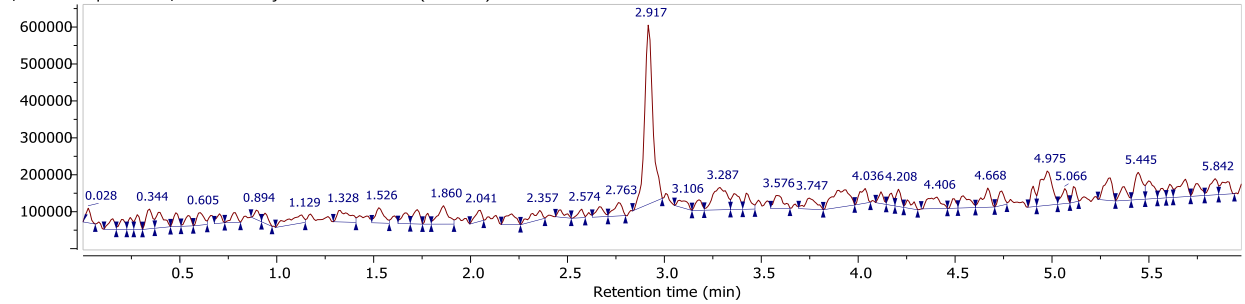


D:/Matt's comp...s LCMS/MM3178.D Injection 1 Function 1 (MM3178) MS + spectrum 2.86..2.99

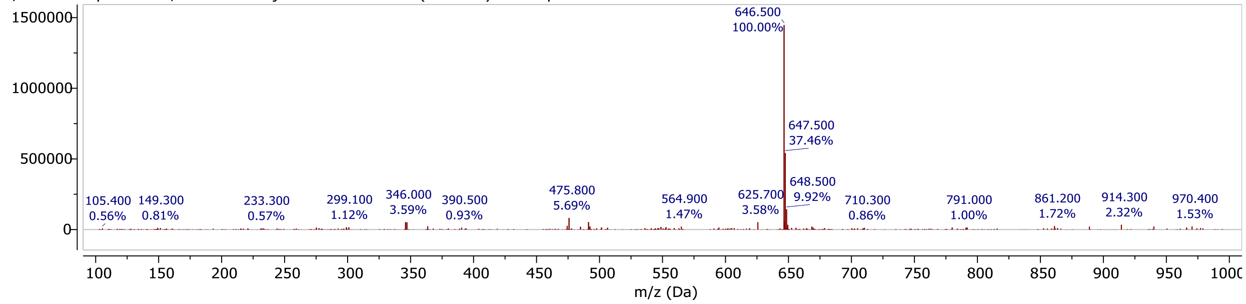


**MM3177 (16)**

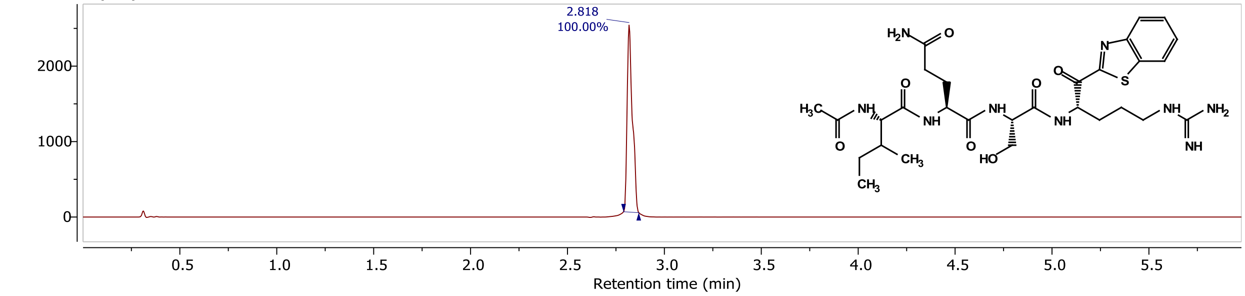


D:\Matt's comp... LCMS\MM3177.D\ Injection 1 Function 1 (MM3177) TIC

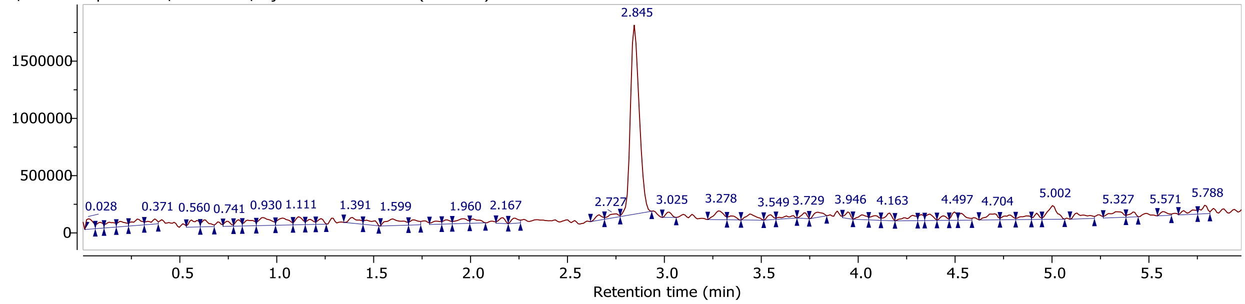


D:\Matt's comp... LCMS\MM3177.D\ Injection 1 Function 1 (MM3177) MS + spectrum 2.79..2.93

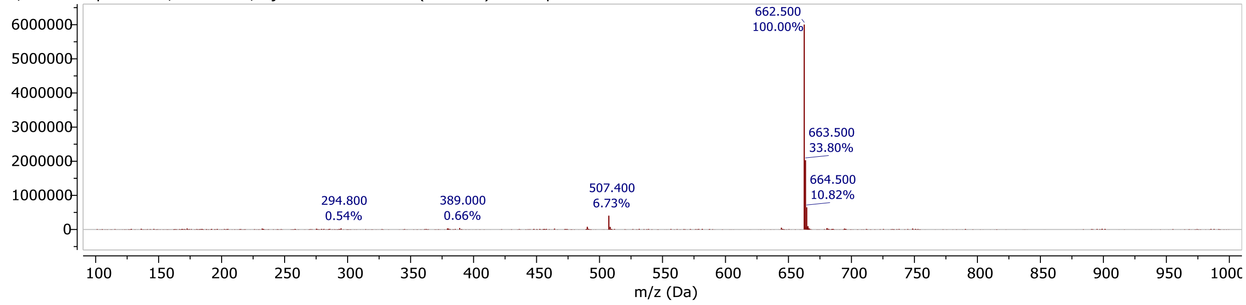

## MM3194 (17)

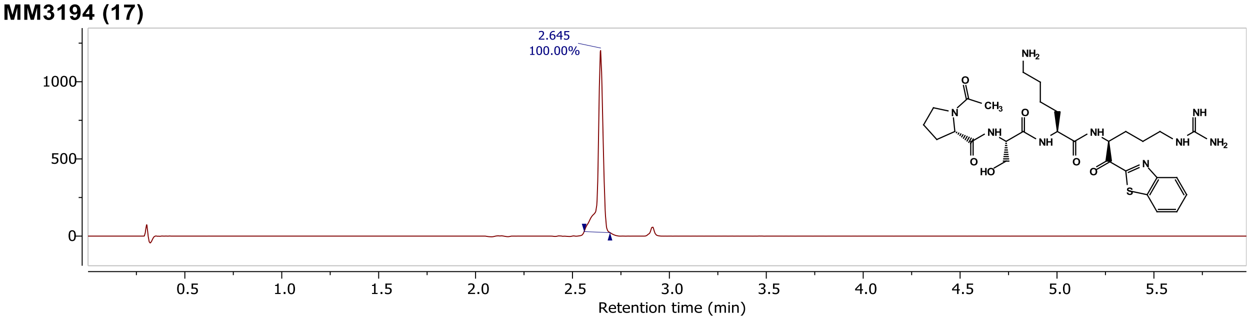


D:/Matt's comp... LCMS/MM3194C.D Injection 1 Function 1 (MM3194C) TIC

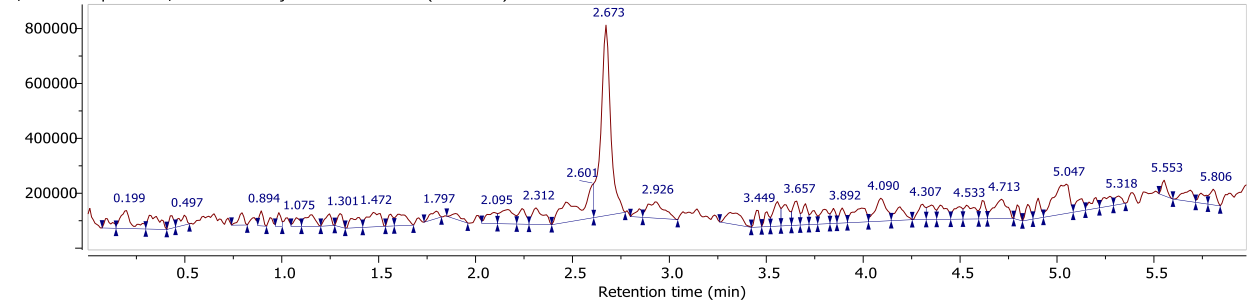


D:/Matt's comp... LCMS/MM3194C.D Injection 1 Function 1 (MM3194C) MS + spectrum 2.60..2.75

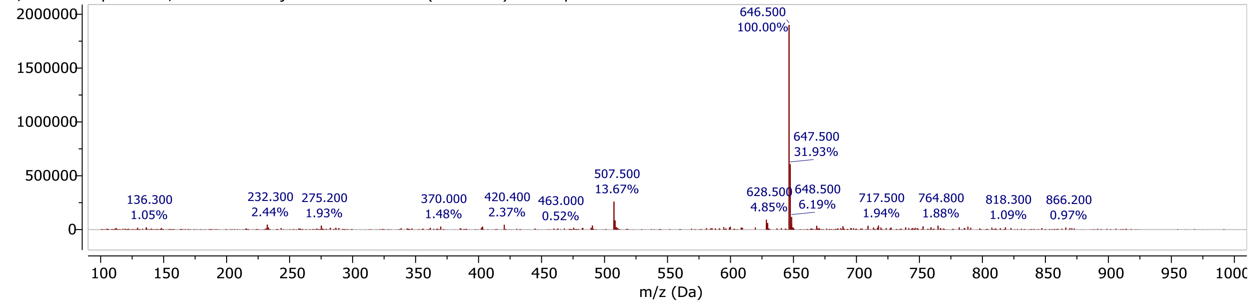


## MM3144 (18)

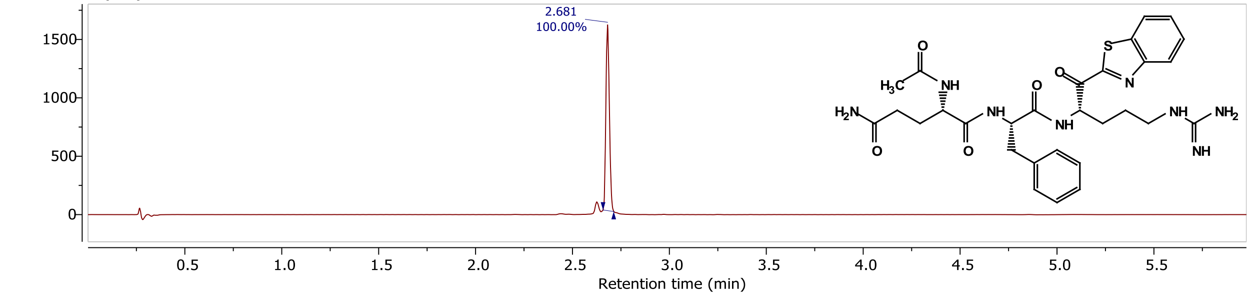


D:/Matt's comp... LCMS/MM3144S.D Injection 1 Function 1 (MM3144S) TIC

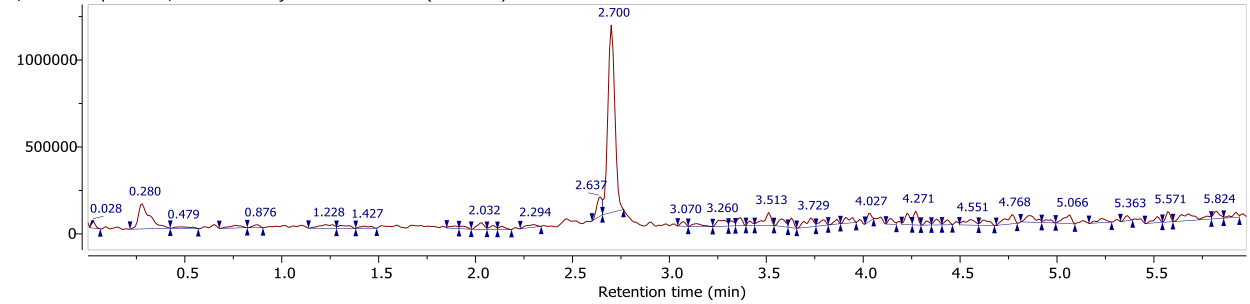


D:/Matt's comp... LCMS/MM3144S.D Injection 1 Function 1 (MM3144S) MS + spectrum 2.70

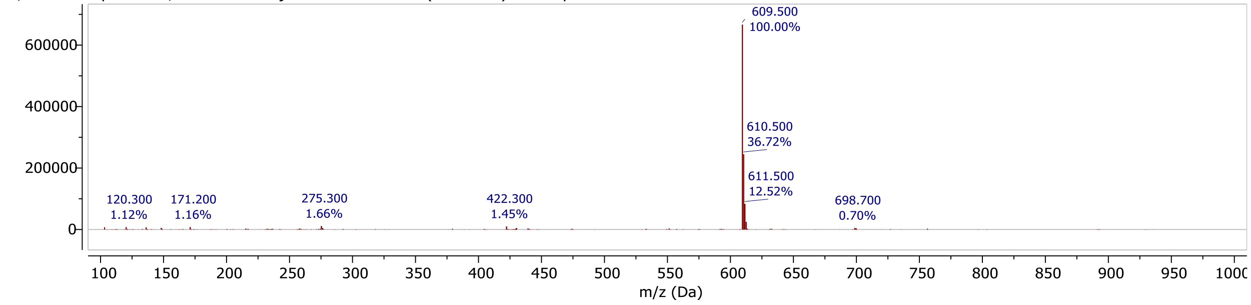

## CA1041 (19)

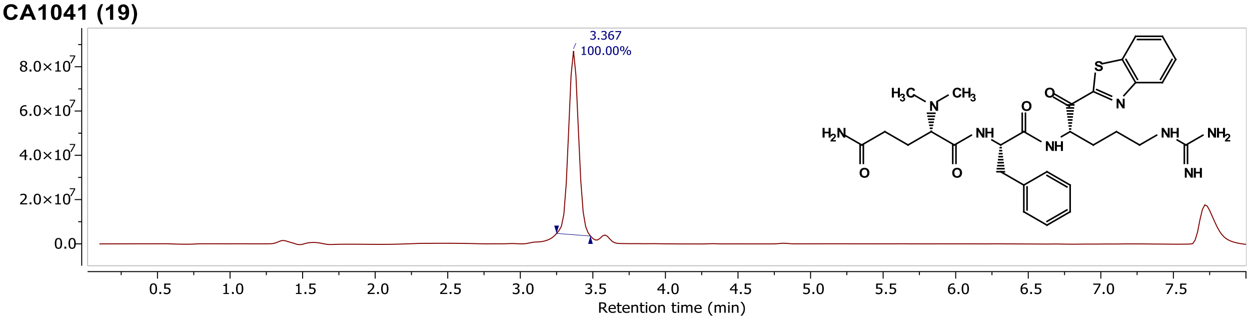


\\waters-lmwb9...\CA1041-2P6.raw Injection 1 MS ES+ TIC

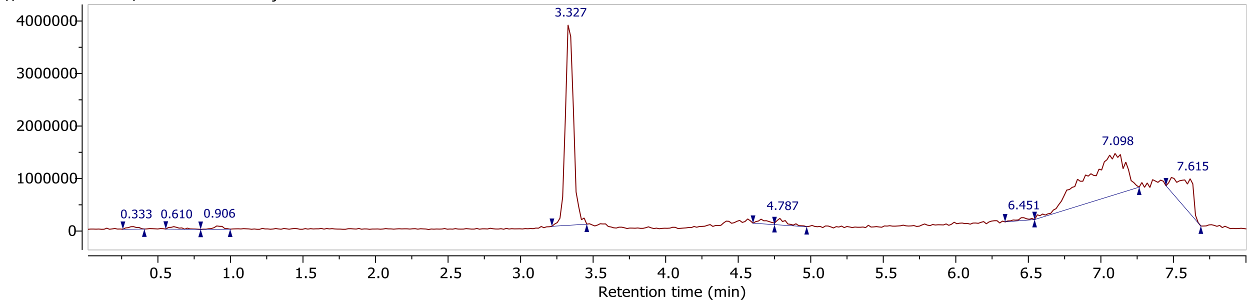


\\waters-lmwb9...\CA1041-2P6.raw Injection 1 MS ES+ MS + spectrum 3.22..3.44

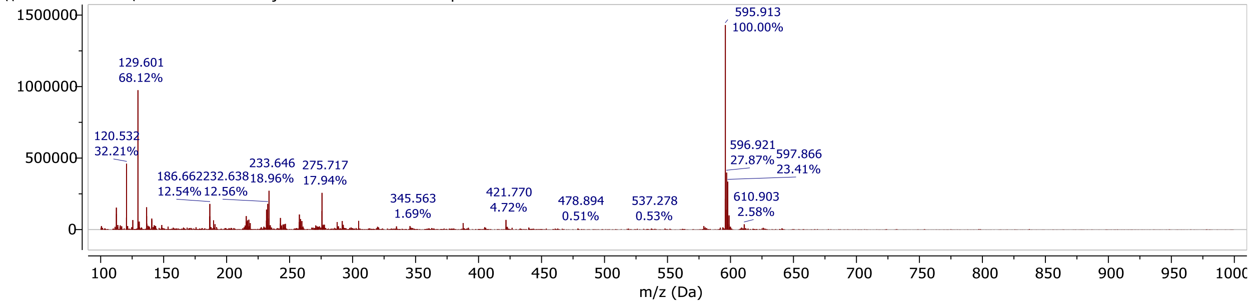


## CA1033 (20)

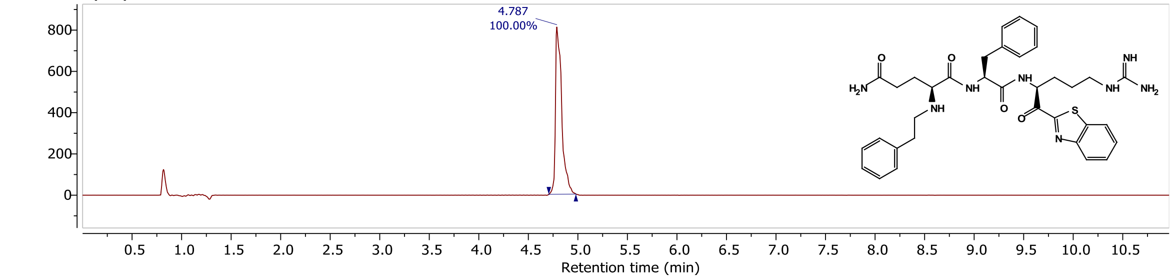


\\quatro-micro...2024\CA1033-2.D Injection 1 Function 1 (CA1033-2) TIC

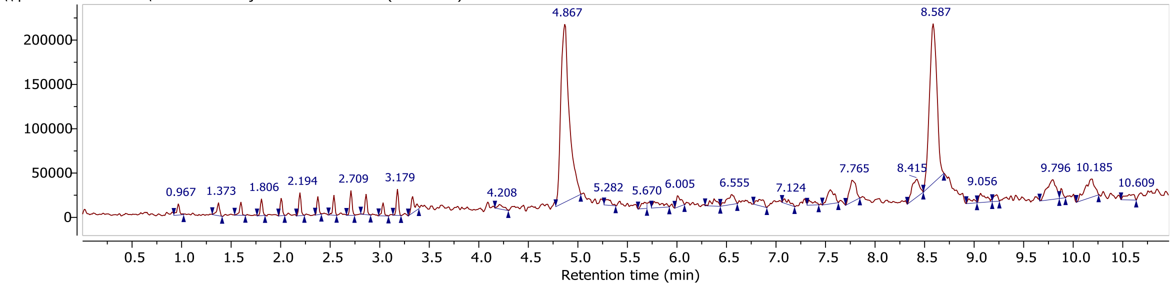


\\quatro-micro...2024\CA1033-2.D Injection 1 Function 1 (CA1033-2) MS + spectrum 4.75..5.00

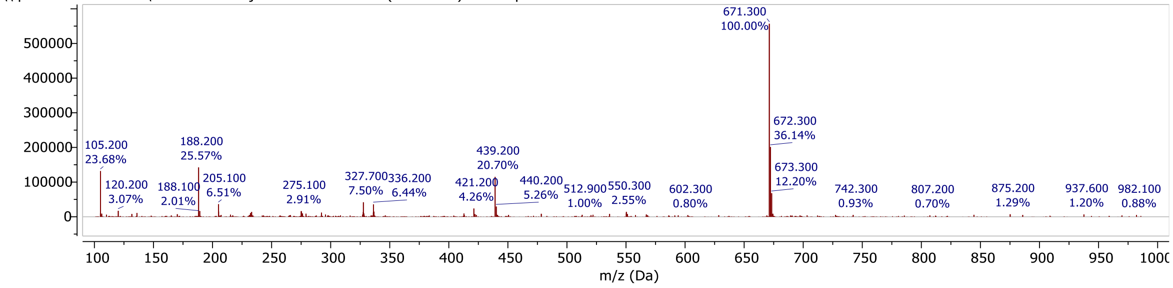

## CA1022 (21)

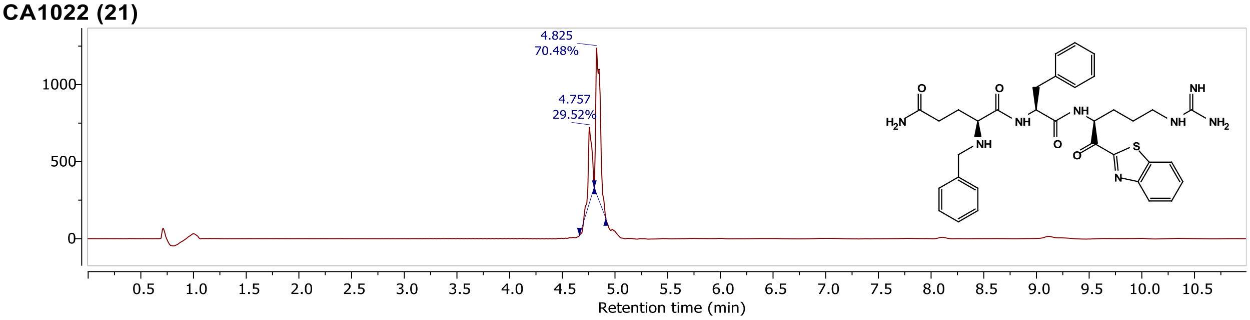


\\quatro-micro...2024\CA1022-2.D Injection 1 Function 1 (CA1022-2) TIC

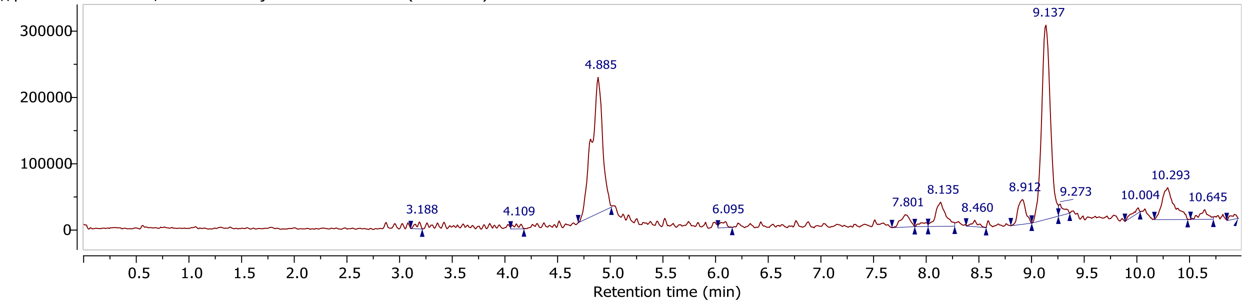


\\quatro-micro...2024\CA1022-2.D Injection 1 Function 1 (CA1022-2) MS + spectrum 4.67..5.02

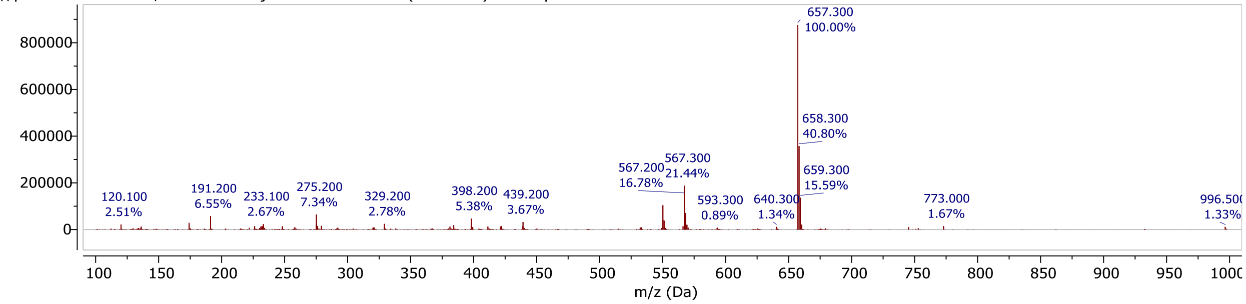


## CA1018 (22)

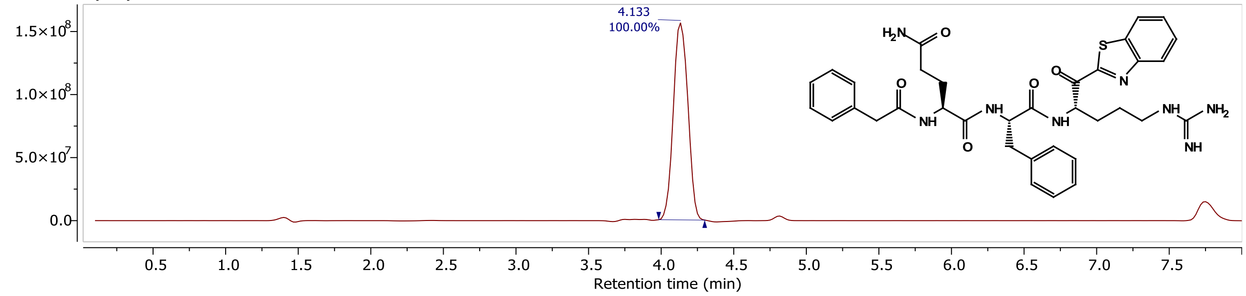


\\waters-lmwb9...a\CA1018PUR.raw Injection 1 MS ES+ TIC

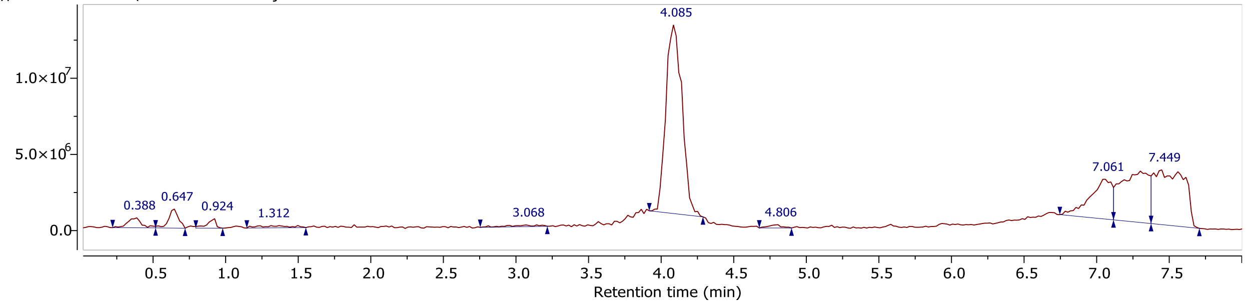


\\waters-lmwb9...a\CA1018PUR.raw Injection 1 MS ES+ MS + spectrum 4.08

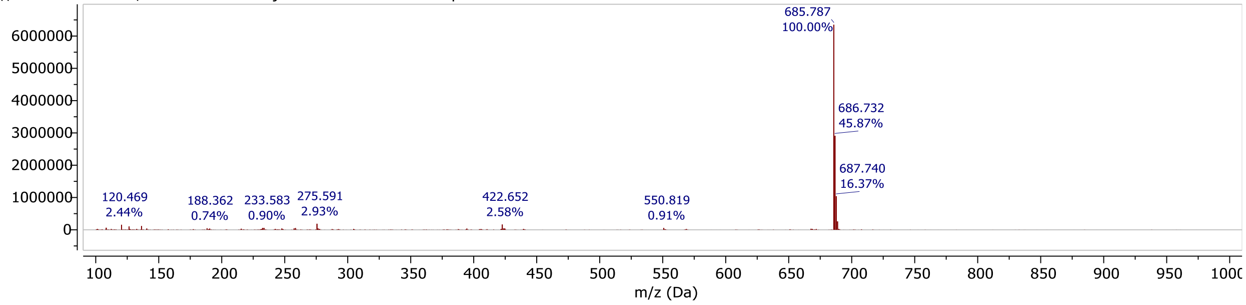

## CA1043 (23)

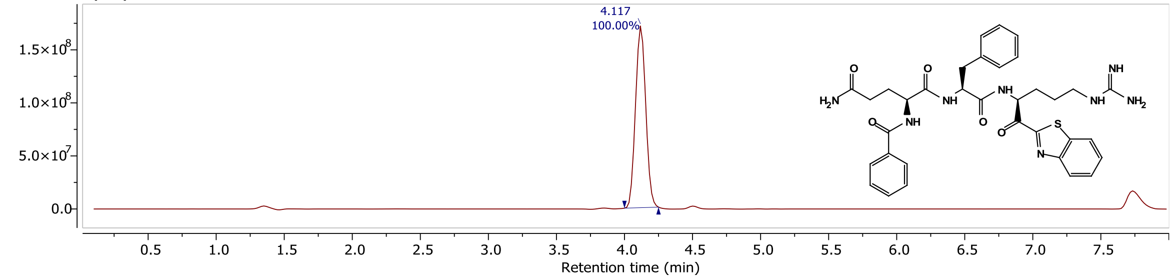


\\waters-lmwb9...\CA1043-252.raw Injection 1 MS ES+ TIC

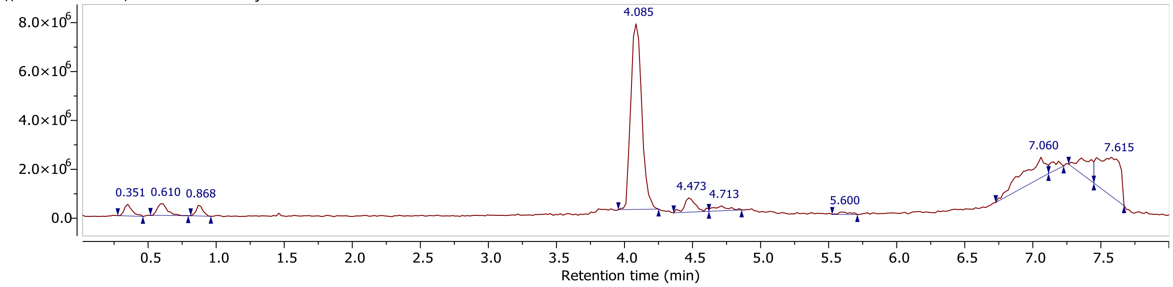


\\waters-lmwb9...\CA1043-252.raw Injection 1 MS ES+ MS + spectrum 4.08

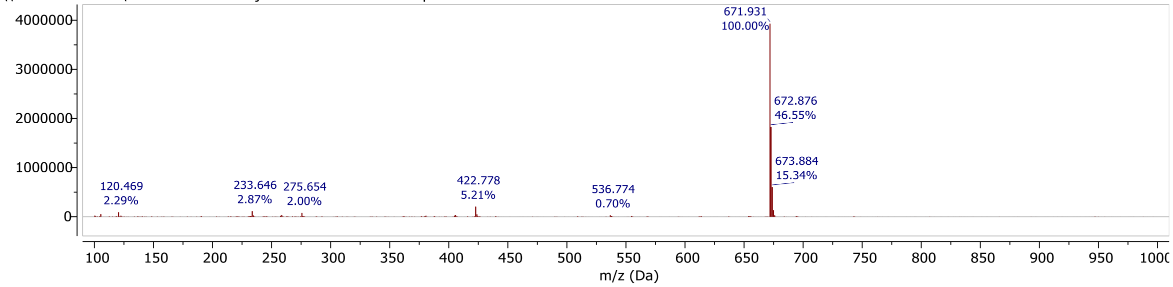


## CA1046 (24)

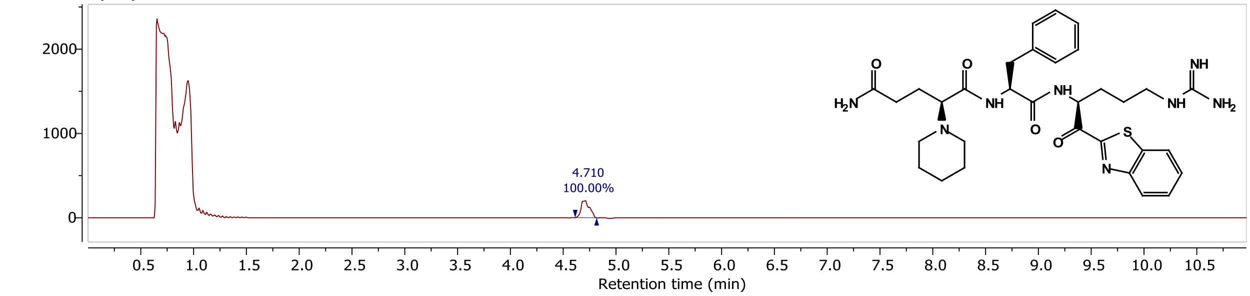


\\quatro-micro...2024\CA1046-1.D Injection 1 Function 1 (CA1046-1) TIC

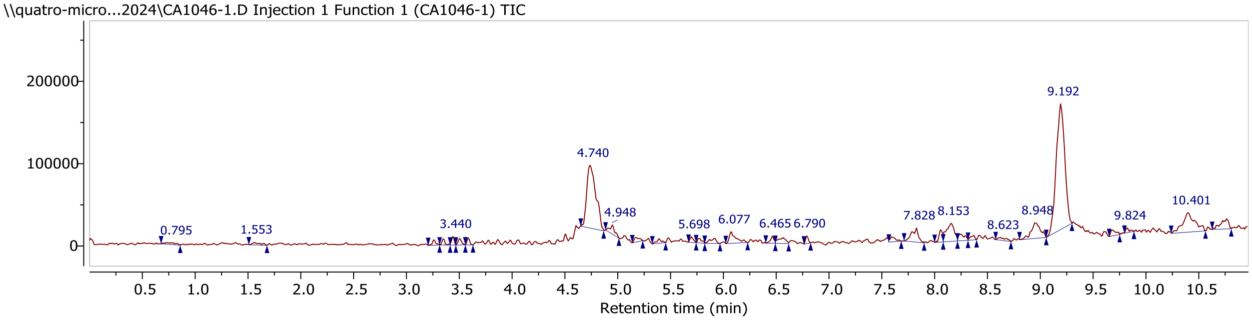


\\quatro-micro...2024\CA1046-1.D Injection 1 Function 1 (CA1046-1) MS + spectrum 4.61..4.80

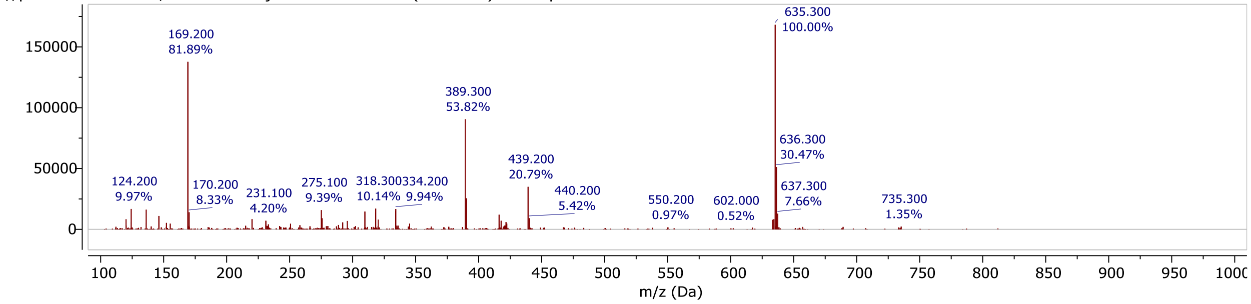

#### ZFH9141 (25)

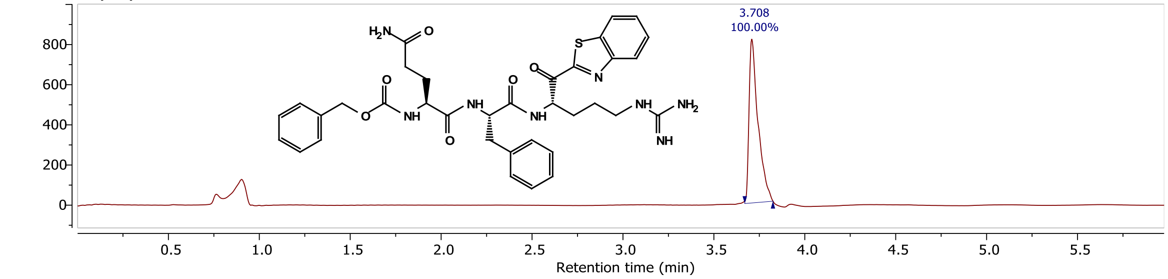


\\quatro-micro...EB2024\9H141P.D Injection 1 Function 1 (9H141P) TIC

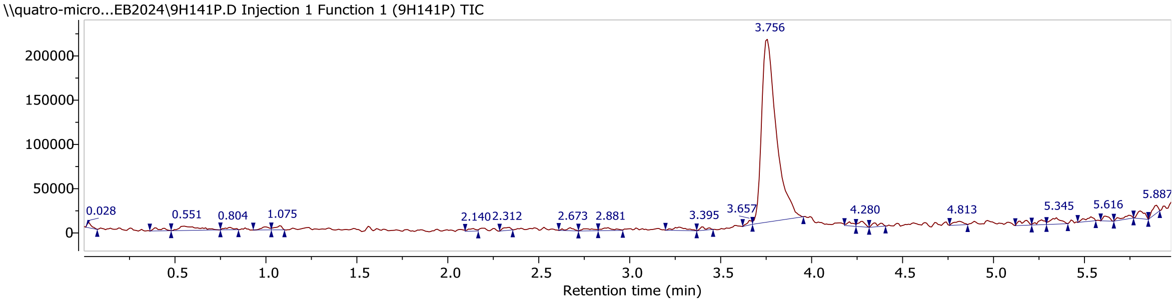


\\quatro-micro...EB2024\9H141P.D Injection 1 Function 1 (9H141P) MS + spectrum 3.76

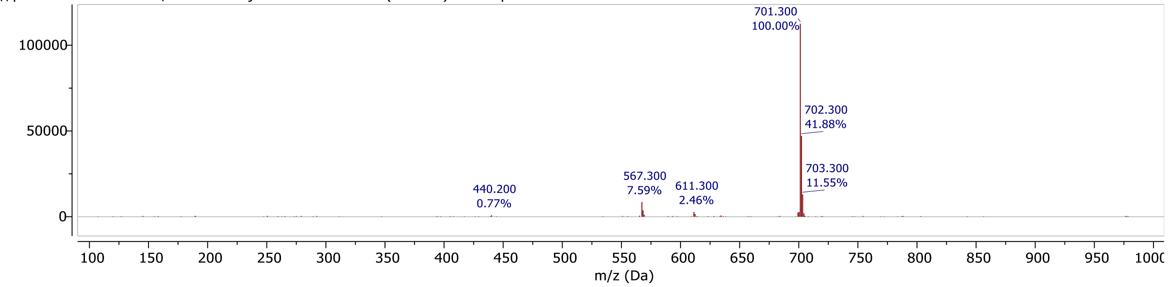


## MF1101 (27)

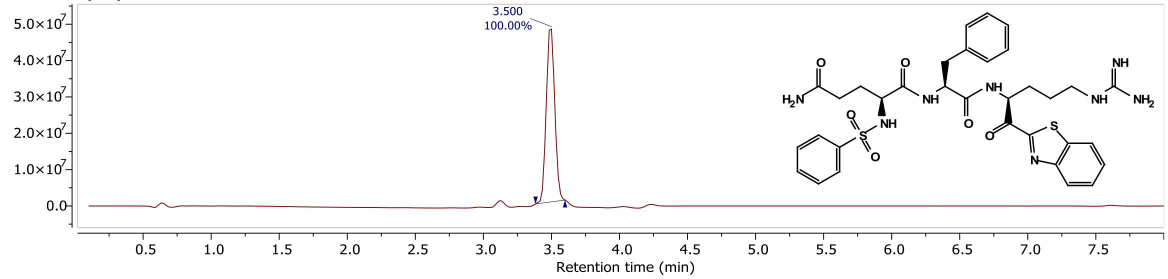


\\waters-lmwb9...a\MF1101F31.raw Injection 1 MS ES+ TIC

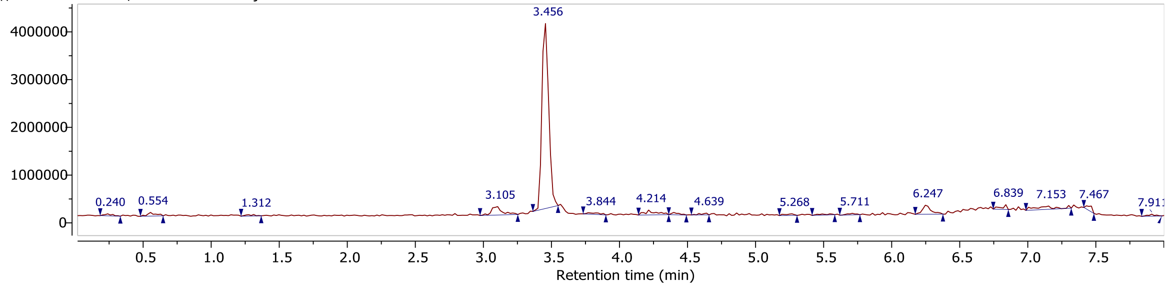


\\waters-lmwb9...a\MF1101F31.raw Injection 1 MS ES+ MS + spectrum 3.46

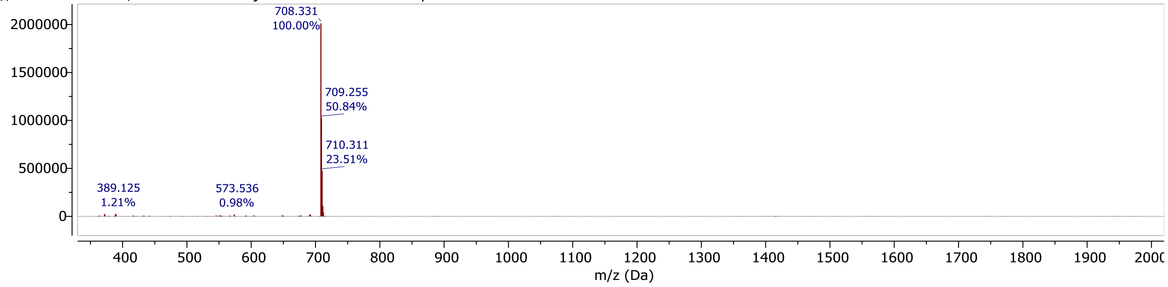

## MF1105 (28)

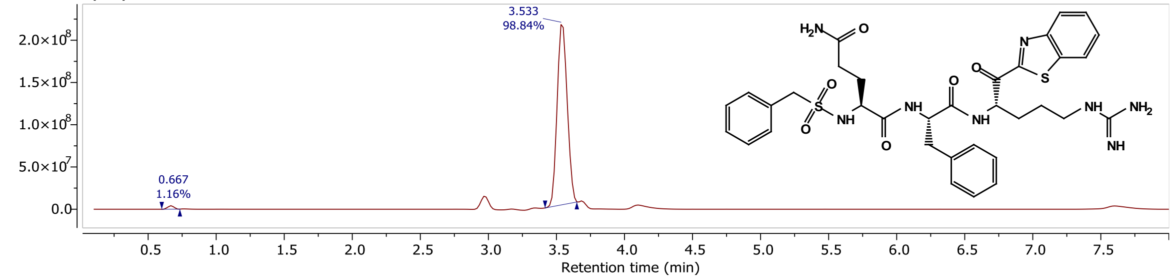


\\waters-lmwb9...a\MF1105F38.raw Injection 1 MS ES+ TIC

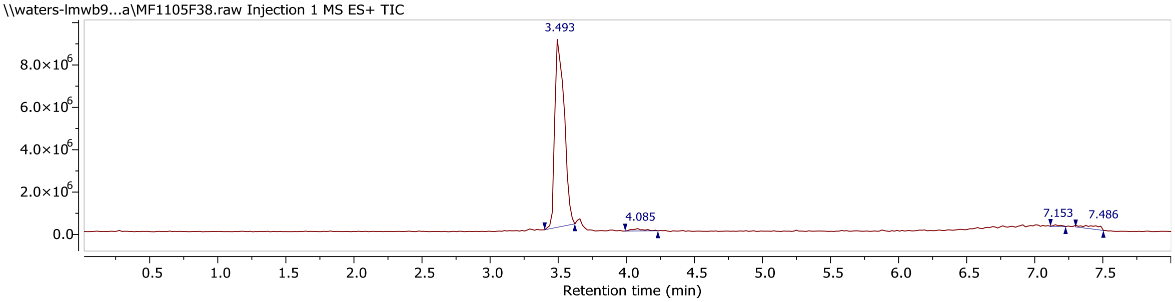


\\waters-lmwb9...a\MF1105F38.raw Injection 1 MS ES+ MS + spectrum 3.49

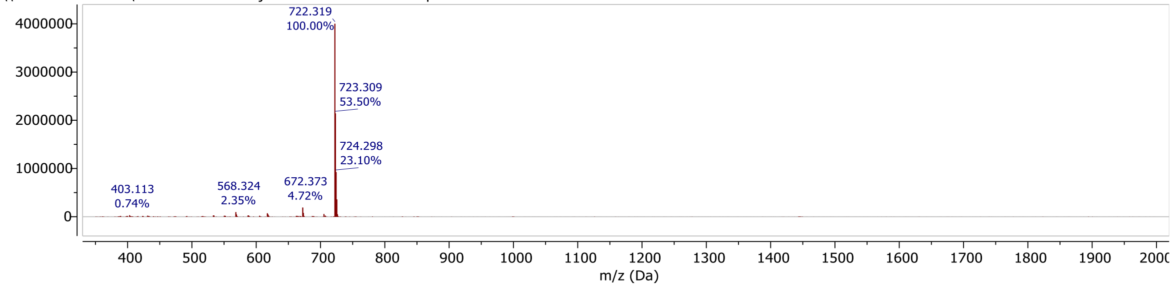


## MF1104 (29)

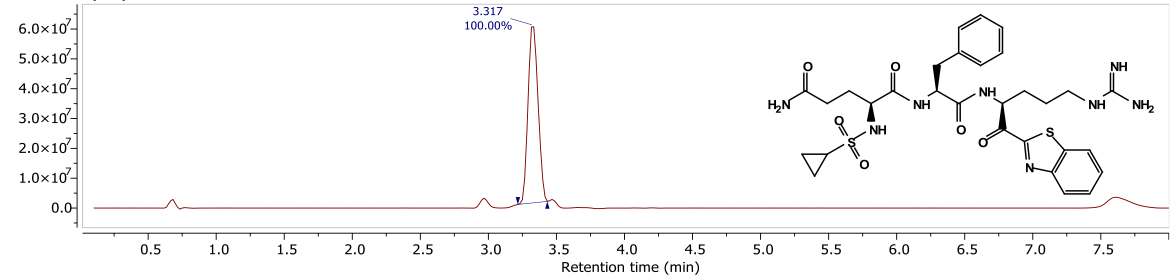


\\waters-lmwb9...a\MF1104F24.raw Injection 1 MS ES+ TIC

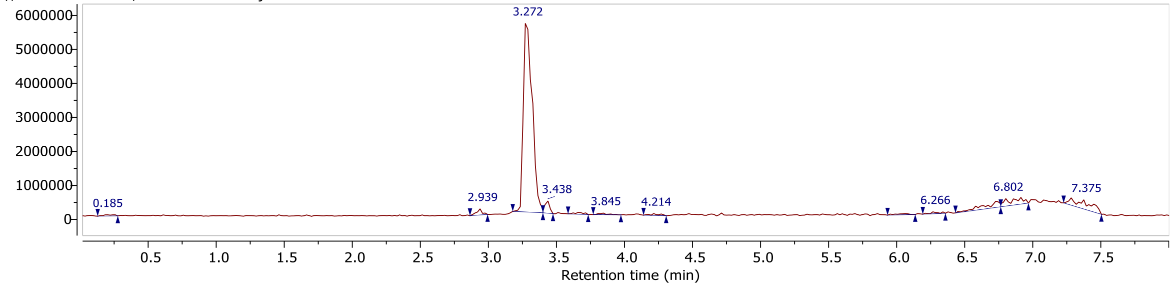


\\waters-lmwb9...a\MF1104F24.raw Injection 1 MS ES+ MS + spectrum 3.27

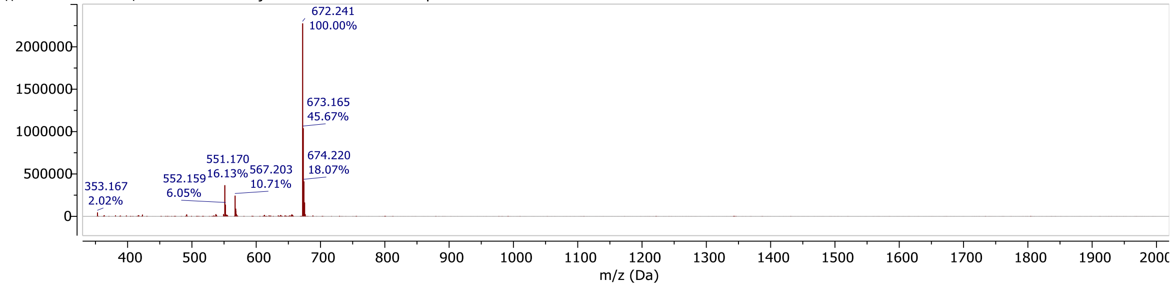

**$K_M$ of Recombinant TMPRSS2.** To calculate $K_M$, Boc-QAR-AMC was serially diluted in DMSO such that the final substrate concentration in the assay ranged from 500 μM to 12.5 μM. Assays were performed in a total volume of 30μL in triplicate wells of black 384-well plates (Nunc),and initial velocity was measured in 35-s intervals for 7 min with excitation of 360 nm and emission of 460 nm on a BiotekHTX plate reader. $K_M$ and $V_{max}$ were calculated from Michaelis-Menten plots (Fig. S1) using GraphPad Prism.

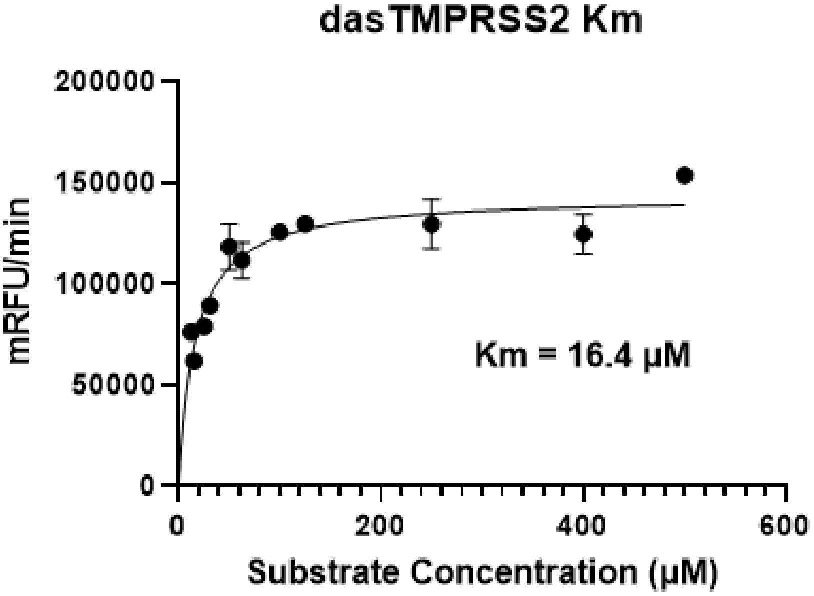


Figure S1: Michaelis-Menten plot of full-length human TMPRSS2, assayed at a final enzyme concentration of 3 nM.
